# Failed allogeneic corneal lenticules show fibrosis and loss of keratocyte phenotype, not rejection

**DOI:** 10.64898/2026.11.04.764222

**Authors:** Hanyu Jiang, Yu Di, Chen Li, Yang Jiang, Ying Li, Yuehua Zhou

## Abstract

**Purpose:** Progressive opacity after corneal stromal tissue implantation is usually attributed to rejection or fibrosis. Across 284 reports, drugs used for opacity were limited to corticosteroids and immunosuppressants, although appearance, diagnosis and steroid response cannot distinguish the two. Tissue evidence that immunosuppression targets the lesion is lacking.

**Methods:** We compared gene expression in six allogeneic corneal stromal lenticules exchanged for progressive opacity in three patients with eight controls, using independent public reference standards also applied to published repair and drug-treatment experiments.

**Results:** Despite differing diagnoses and steroid responses, all six lenticules expressed abundant scar matrix and separated completely from controls (exact P = 0.0061). T cell, cytotoxic and interferon-γ programs were not elevated, identifying fibrosis rather than rejection. Fibrosis involves remodeling, as does recoverable injury. Distinguishing recovery from failure therefore required the keratocyte phenotype, that is, whether keratocyte-characteristic gene expression is preserved. Rabbit corneas that regained transparency and those that scarred showed similarly active remodeling. Phenotype approached uninjured levels with recovery and was significantly reduced with scarring. Failed lenticules had lower phenotype scores than all controls. Tacrolimus and rapamycin both suppressed rejection in mouse grafts, but phenotype increased with tacrolimus and decreased with rapamycin. Suppressing rejection was therefore insufficient to restore phenotype. Resveratrol reduced scar matrix expression, increased phenotype and prolonged graft survival in rats.

**Conclusions:** Failed lenticules lacked T cell–mediated rejection to target. Treatment should shift from intensified immunosuppression to controlling remodeling and preserving phenotype. Phenotype assessment provides a common basis for explaining failure, designing treatment and comparing efficacy.

## Introduction

Corneal stromal tissue implantation is an approach to additive corneal surgery. Stromal lenticules removed during small-incision lenticule extraction (SMILE) can be implanted as donor tissue in another person’s cornea to alter refractive power or increase thickness [1]. The goal is integration of the implant into the recipient cornea as ordered, transparent stroma. Progressive opacity prevents integration and leads to implant removal or exchange [2].

Implant opacity is generally attributed to immune rejection or fibrosis. Rejection is mediated predominantly by CD4 helper T cells, chiefly interferon-γ-secreting Th1 cells, and CD8 cytotoxic T cells [3]. Fibrosis involves abundant, disorganized scar matrix produced by fibroblasts and myofibroblasts [4]. Management follows corneal graft rejection protocols and relies primarily on corticosteroids [5]. Across 284 published reports of corneal stromal tissue implantation, drug treatment for opacity comprised only corticosteroids and immunosuppressants. None used antifibrotic drugs. Yet slit-lamp appearance, clinical diagnosis, steroid response and the course after exchange cannot distinguish rejection from fibrosis. Gene expression was not analyzed in any of the 346 reported specimens from implant removal or biopsy. Tissue evidence is therefore lacking to establish whether opacity reflects rejection or fibrosis and whether immunosuppression targets the lesion.

After stromal injury, both recovery and scarring begin with remodeling as keratocytes become fibroblasts and differentiate toward myofibroblasts. After mild injury, remodeling stops and transparency returns rapidly. With persistent remodeling, myofibroblasts mature and the stroma scars. Once repair is complete, myofibroblasts undergo apoptosis and keratocytes repopulate the stroma. Established scars can resolve over months to years, with permanent scars confined to severe injury [4]. Remodeling and scar matrix thus occur during both recovery and failure. Identifying fibrosis as the cause of opacity does not establish whether an implant will ultimately recover or fail.

Keratocytes express abundant corneal crystallins that minimize cellular light scattering, one condition for corneal transparency [6]. Expression of keratocan, ALDH3 and keratan sulfate proteoglycans falls as keratocytes become fibroblasts and myofibroblasts [7]. Crystallin levels are also much lower in opaque regions of rabbit corneas after freeze injury [8]. The keratocyte phenotype, that is, whether keratocyte-characteristic gene expression is preserved, is therefore associated with corneal transparency. Whether phenotype distinguishes recovery from failure in implanted corneal stromal tissue, and whether suppressing rejection or scarring restores phenotype, remains unknown.

We measured gene expression directly in six allogeneic corneal stromal lenticules exchanged for progressive opacity in three patients, using unimplanted donor lenticules as controls. Reference standards for rejection and keratocyte phenotype were derived from independent public data. We applied the same standards to published data on corneal repair, cell culture and drug treatment after transplantation. We assessed whether opacity reflected rejection or fibrosis, whether phenotype distinguished recovery from failure, and whether suppressing rejection or scarring was sufficient to restore phenotype.

## Materials and methods

### Patients, specimens and clinical assessment

Three women aged 32–37 years underwent bilateral implantation of fresh allogeneic corneal stromal lenticules. The lenticules were subsequently removed and replaced in the same operation for progressive opacity. We analyzed six failed lenticules and four unimplanted lenticules from independent small-incision lenticule extraction (SMILE) donors (Fig. 1D). Four public human lenticule libraries from PRJNA1533946 were processed with the same alignment and counting pipeline, extending the comparison to eight controls. The study was approved by the institutional ethics committee, and all patients provided informed consent. The study followed the Declaration of Helsinki.

**Figure 1.**
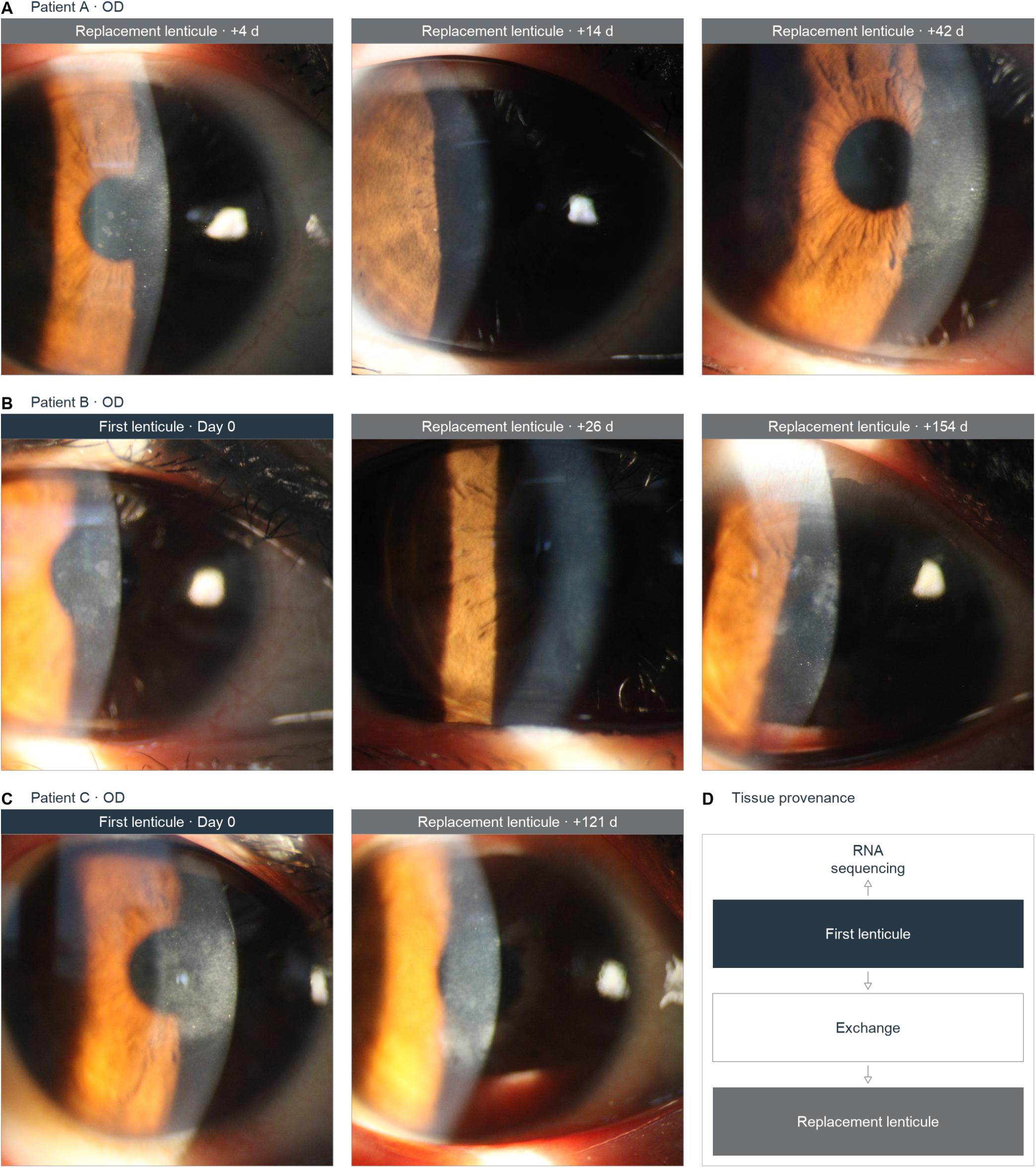
Early clearing after exchange precedes recurrent lenticule opacity. (A–C) Serial right-eye slit-lamp photographs of Patient A at +4, +14 and +42 days (A), Patient B at day 0, +26 and +154 days (B), and Patient C at day 0 and +121 days (C). Day 0 precedes exchange and shows the failed first lenticule; positive days show the replacement lenticule. Dark and gray bands identify the first and replacement lenticules, respectively. Patient A’s band became smoother before granular scattering increased; Patient B’s granular opacity cleared and recurred; Patient C’s replacement lenticule had granular opacity at late follow-up. (D) Tissue provenance. RNA sequencing examined the six failed first lenticules from both eyes of the three patients; clinical follow-up after exchange concerns the replacement lenticules. OD, right eye.

Diagnoses and treatments were transcribed from primary records. Interface reflectivity was measured on Optovue RTVue XR optical coherence tomography (OCT) scans as the mean signal at 50–300 μm divided by that at 400–600 μm. Bilateral values were averaged by patient. Pentacam Scheimpflug densitometry measured the central 0–2 mm zone separately in the anterior 120 μm, intervening stroma and posterior 60 μm [9]. Repeat scans were averaged by eye, then by patient. Clinical intervals and treatments are reported in Table 1; imaging procedures and reference measurements are detailed in the Supplemental methods.

**Table 1.** Failed first lenticules and the clinical course after replacement in Patients A–C.

|  | **Patient A** | **Patient B** | **Patient C** |
|---|---|---|---|
| Eyes implanted / explanted (first lenticules) | 2 / 2 | 2 / 2 | 2 / 2 |
| Interval from first implantation to explantation (d) | 225 | 394 | 90 |
| First documented event leading to exchange (d after implantation) | 148 | 394* | 57† |
| Diagnostic wording in the clinical record | Implant opacity in examination entries; “rejection” appears only in a later summary heading written when exchange was being scheduled | Implant opacity, punctate pocket opacity and low-grade reflectivity; no “rejection” entry | “Rejection” recorded after exchange, at +53 and +121 d, concerning the replacement lenticules |
| Topical corticosteroid treatment | Loteprednol, tapered; fluorometholone after exchange | Fluorometholone and loteprednol; the replacement-lenticule regimen also included ciclosporin | Loteprednol four times daily for approximately two months, on the replacement lenticules |
| Documented treatment response | Dose escalation, tapering and discontinuation documented; effect not stated | Partial improvement of replacement-lenticule opacity, followed by worsening during dose reduction | “No change” in the replacement lenticules |
| Lenticule provenance | Fresh donor tissue (team-reported; written provenance records unavailable) | Fresh donor tissue (team-reported) | Fresh donor tissue (team-reported) |
| Follow-up after exchange (d) | 142 | 154 | 121 |
| Status at last record | No further opacity or rejection entry; the implant was not specifically described | Recurrent punctate opacity and low-grade reflectivity | “Rejection, no change” |
| Mean keratometry at 42 → 142 d after exchange (D) | OD: approximately 50.9 → 59.0; OS: approximately 49.4 → 55.0 | — | — |
| Recorded uncorrected visual acuity at 60 → 142 d after exchange (decimal) | OD: 0.7 → 0.3; OS: 0.8 → 0.1 | — | — |
| Bilateral qOCT visits contributing to Fig. 2B | 7 | 10 | 5 |
| Endothelial cell density at failure recognition or exchange (cells/mm²; OD / OS) | 3,129 / 3,296 | 2,339 / 2,500 | 3,175 / 2,825 |

\* In Patient B, the documented event leading to exchange and the exchange itself occurred on the same day.
† In Patient C, an earlier entry described implant opacity at 36 d; the 57-d entry was the event that led to exchange.
All three patients were women aged 32–37 years at first implantation. RNA sequencing examined the six failed first lenticules. Treatment responses and follow-up concern the replacement lenticules where indicated. Values are transcribed from the clinical records; intervals run from first implantation or from exchange as specified. Endothelial cell densities fall within each report’s printed reference interval. Uncorrected acuity retains the recorded decimal scale, and keratometry retains the reported approximate precision. Em dashes identify items not recovered from the records. The cohort comprises six failed first lenticules from six eyes of three patients and 22 bilateral qOCT visits. OD, right eye; OS, left eye; qOCT, quantitative optical coherence tomography.

### Published implantation reports

Human corneal stromal tissue implantation reports were independently read twice, with disagreements adjudicated by a third reader. We summarized adjudicated procedure, event, author attribution, treatment sequence and specimen-examination records. Reports, eyes and specimens were counted separately; missing quantities remained unreported. Eligibility and source linkage are described in the Supplemental methods.

### RNA sequencing and differential expression

Individual lenticules underwent paired-end RNA sequencing on an Illumina NovaSeq X Plus (2 × 150-bp reads). The sequencing provider aligned reads to GRCh38.p12 with HISAT2 v2.2.1 [10] and counted genes with featureCounts [11]. Independent recounting reproduced the supplied counts (Supplemental Fig. S21B). Salmon provided alternative quantification [12] (Supplemental Fig. S21C).

Bilateral counts were summed by patient; donors remained independent units (Supplemental Fig. S1). Non-host reads were classified separately, and taxon recovery was tested by read-depth resampling (Supplemental Fig. S10). Differential-expression analysis used edgeR v4.6.2 [13,14], trimmed mean of M values (TMM) normalization [15], condition-specific filtering with filterByExpr, a group-only design, dispersion estimation and quasi-likelihood F-tests [16]. Gene-level P values were adjusted by the Benjamini–Hochberg procedure [17].

### Tissue and immune programs

The scar matrix, stromal small leucine-rich proteoglycan and transparency panels contained seven, six and seven nonoverlapping genes, respectively. After mitochondrial genes were removed and counts aggregated by symbol, expression was converted to log₂(CPM + 1), where CPM denotes counts per million. Each gene was standardized across independent patient and control units and centered on controls. Panel scores averaged these values.

Transparency panel scores were displayed in their biological direction and reversed for directional tests. The same definitions were used for both quantifications and the expanded control comparison. Captured-RNA fractions were estimated with BayesPrism and MuSiC against two single-cell references (Supplemental Fig. S2). Spatial associations between myeloid and stromal programs were quantified in 13 public human corneas (Supplemental Fig. S5).

Immune-module enrichment used patient-level edgeR log₂ fold-change ranks; the Hallmark overview [18] used DESeq2 Wald-statistic ranks [19]. Both used multilevel fgsea with eps = 0 [20]. Adjustment families and leading-edge annotation are in the Supplemental methods.

The rejection score combined five immune gene modules: T cell receptor execution, cytotoxic T/NK activity, interferon-γ chemokine, B/plasma cell and antigen presentation. NK denotes natural killer cells. Nineteen one-to-one orthologs were shared by both mouse calibration datasets and both human quantifications. Genome-wide effect ranks were scaled to [−1, 1] and averaged within module. Projection onto the GSE275249 phosphate-buffered saline (PBS)-treated allograft-minus-isograft vector, divided by its squared length, set that contrast to +1. GSE181893 PBS-minus-anti-CD80/86 supplied an independent reference [21].

TRUST4 v1.1.10-r639 reconstructed receptors from FASTQ and aligned BAM files [22]. Clonotypes were compared at a common read depth and required productive, chain-consistent sequences, read support, reproducibility across resampling runs and recovery from BAM input. Full criteria are in the Supplemental methods.

### Keratocyte phenotype and remodeling

Phenotype weights came from two independent human single-cell resources [23,24]. One contrasted quiescent keratocyte-like cells with corneal fibroblasts; the other contrasted corneal stromal fibroblasts with fibroblasts from other ocular tissues. Donor-level expression differences were standardized within each resource and averaged across 12,526 shared genes. The phenotype score was the inner product of these consensus weights and ranked gene effects, divided by the sum of available absolute weights. Both source weightings were analyzed separately.

The failure signature comprised the 300 most upregulated and 300 most downregulated genes in the patient-level comparison with four donor controls. Failure-signature similarity was the mean of forward and reciprocal signed-rank projections, each normalized by its rank-permutation null standard deviation. Cross-species comparisons used one-to-one orthologs. Core matrix expression averaged ranked effects across NABA core-matrisome genes [25] and was compared with phenotype across repair and disease states (Supplemental Fig. S11). Repair and implant-remodeling reference projections, with and without mutual residualization of reference weights, and the expression-matched random-reference test are shown in Supplemental Figs. S4, S6 and S22. Unit-level tests and group-level estimates were calculated separately; full estimators are in the Supplemental methods.

### Public experiments and cell populations

Published datasets were analyzed within study, retaining treatment, comparator, sampling time and animal, culture or pooled-library units. Rabbit stromal pseudobulks represented two injured–contralateral pairs at each freeze-injury time and three fibrotic and three naive animals after alkali burn [26,27]. Culture comparisons used three rabbit libraries per condition [28] and five feline replicates per condition [29]. Optical, structural and survival observations were linked to the corresponding transcriptomic comparisons in eight independent in vivo experiments. Drug and culture contrasts, individual-library projections and outcome linkage are detailed in the Supplemental methods.

We compared anti-Gr-1 plus tacrolimus directly with tacrolimus alone in day-15 mouse allografts [30]. Observed cell-type expression and captured RNA shares were recombined to separate state and composition contributions. Losartan-treated and saline-treated corneas were compared by stromal cluster [31]. In GSE272700, Aire-deficient mice with or without 14 days of rewetting were compared with wild-type mice using single-nucleus data [32].

Stromal pseudobulks were scored before and after library-depth equalization. These analyses, including support-set residuals, are in the Supplemental methods.

### Statistical analysis

Analyses used R 4.5.0 and Python 3.10.12. Patients were the units of inference for failed tissue. Exact tests enumerated all 35 assignments of three failure labels among seven original units and all 165 among eleven expanded-comparison units. Panel tests used directional mean differences, with Benjamini–Hochberg adjustment across the three panels. Phenotype and failure-signature tests recomputed their complete estimators and evaluated mean differences and complete separation separately. Gene-level and enrichment significance was defined as adjusted P < 0.05. Bootstrap units and procedures are in the Supplemental methods. Clinical trajectories were summarized by patient.

## Results

### Diagnoses and steroid responses differed among patients, and early clearing after exchange did not persist

The six initial lenticules in both eyes of three patients were exchanged for progressive opacity 90–394 days after implantation (Table 1). These were termed failed lenticules. Before removal, optical sections in Patient B and Patient C showed central gray-white granular opacity (Fig. 1B and C). After exchange, Patient B’s opacity became a smooth gray-blue band on day 26, but granular opacity recurred on day 154 (Fig. 1B). Patient A’s band was smoother on day 14 than on day 4, but granular scattering increased again by day 42 (Fig. 1A). Patient C still had pronounced gray-white granular opacity on day 121 (Fig. 1C).

On OCT, increased reflectivity was confined to the lenticule interface and adjacent anterior stroma (Fig. 2A). Interface reflectivity was measured as the anterior-to-posterior mean signal ratio, with no corresponding increase in the posterior reference region, Descemet’s membrane or endothelial surface (Fig. 2A and B). Scheimpflug densitometry [33] showed the same depth distribution: central anterior density was 30.07 grayscale units (GSU) at failure versus 15.28 GSU in the preimplantation reference, whereas posterior values were 6.91 and 7.10 GSU, respectively (Fig. 2D). Endothelial cell density was preserved, and photographs showed no neovascularization or ciliary injection.

**Figure 2.**
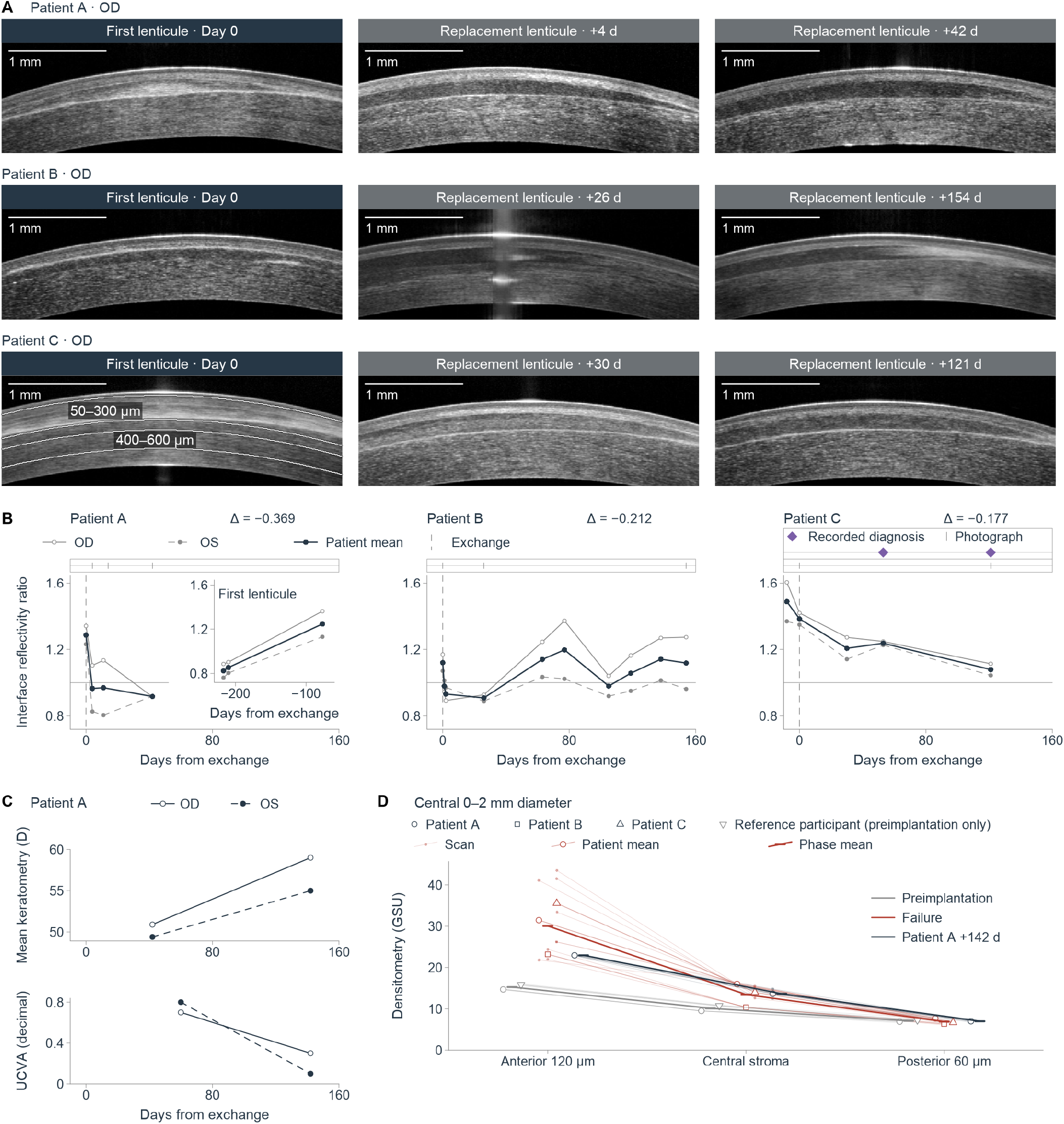
Interface reflectivity falls early after exchange while later corneal shape, vision and anterior density remain abnormal. (A) Nine right-eye optical coherence tomography (OCT) sections from Patients A–C at the indicated days relative to exchange. Bands identify first and replacement lenticules. Patient C’s day-0 image marks the anterior 50–300 μm and posterior 400–600 μm regions used for the reflectivity ratio. Scale bars, 1 mm. Seven sections have same-day photographs in Fig. 1. (B) Interface reflectivity, defined as anterior divided by posterior mean signal, at 44 eye-visits and 22 bilateral patient-visits. Pale open/filled circles and solid/dashed lines denote OD/OS; dark points and lines denote means of unrounded bilateral ratios. The inset contains Patient A’s earlier first-lenticule visits. Dashed vertical lines mark exchange, and horizontal lines mark a ratio of one. Event tracks aligned above the reflectivity plots show photographs (short ticks) and recorded rejection diagnoses concerning Patient C’s replacement lenticules (purple diamonds). Displayed changes compare day 0 with +42, +26 and +30 days for Patients A, B and C, respectively. (C) Patient A’s mean keratometry, in diopters, at +42 and +142 days and decimal uncorrected visual acuity at +60 and +142 days. Open/filled symbols identify OD/OS. (D) Central 0–2 mm-diameter Scheimpflug densitometry in grayscale units (GSU), across the anterior 120 μm, central stroma and posterior 60 μm. Faint trajectories and small points show 15 scans; open shapes show patient means after averaging within eye and then patient; thick lines and short marks show phase means. Circles, squares and triangles identify Patients A, B and C; inverted triangles identify the participant with preimplantation scans only. Gray, red and dark trajectories denote preimplantation (two participants), failure (three patients) and Patient A at +142 days, respectively. OS, left eye.

Diagnostic terms differed, and the records distinguished initial from replacement lenticules (Table 1). Patient A’s examinations described implant opacity, with “rejection” appearing only in a summary heading when exchange was scheduled. Patient B’s records described implant or punctate pocket opacity without documenting rejection. Patient C’s explicit rejection diagnosis was made after exchange and concerned the replacement lenticules (Fig. 2B). Steroid responses were recorded for replacement lenticules. Patient B’s opacity partially improved with corticosteroids and cyclosporine but worsened during tapering. Patient C’s opacity remained unchanged after topical loteprednol four times daily for approximately two months. The first recorded event leading to exchange occurred 77 days before removal in Patient A, on the removal day in Patient B, and 33 days before removal in Patient C.

Interface reflectivity initially decreased in all three patients after exchange (Fig. 2B), followed by recurrent opacity in Patient B and Patient C (Fig. 1B and C). Patient A had no further recorded opacity or rejection within 142 days, but mean keratometry increased bilaterally and anterior density remained 56.1% above preimplantation levels (Fig. 2C and D; Supplemental Fig. S12A and B). Between days 60 and 142 after exchange, uncorrected visual acuity declined from 0.7 to 0.3 in the right eye and 0.8 to 0.1 in the left (Fig. 2C; Table 1). Early clearing after exchange preceded recurrent opacity or continuing changes in corneal shape and vision, without establishing integration (Supplemental Figs. S18–S20). Clinical observations could not distinguish rejection from fibrosis; tissue evidence from failed lenticules was required.

### Failed lenticules expressed abundant scar matrix and had lower phenotype scores than all controls

We compared six failed lenticules with eight controls: four unimplanted donor lenticules and four public human lenticule samples from PRJNA1533946. Scar matrix panel expression increased [34], whereas transparency panel expression, including corneal crystallins, decreased [8]. Both panels separated every patient from every donor control with featureCounts and Salmon quantification (Supplemental Table S1). Complete separation persisted with all eight controls at the patient level (exact P = 0.0061 for each panel; Fig. 3C). Among 871 matrisome and transparency genes, 129 increased and 33 decreased at a patient-level false discovery rate (FDR) < 0.05 (Fig. 3B).

**Figure 3.**
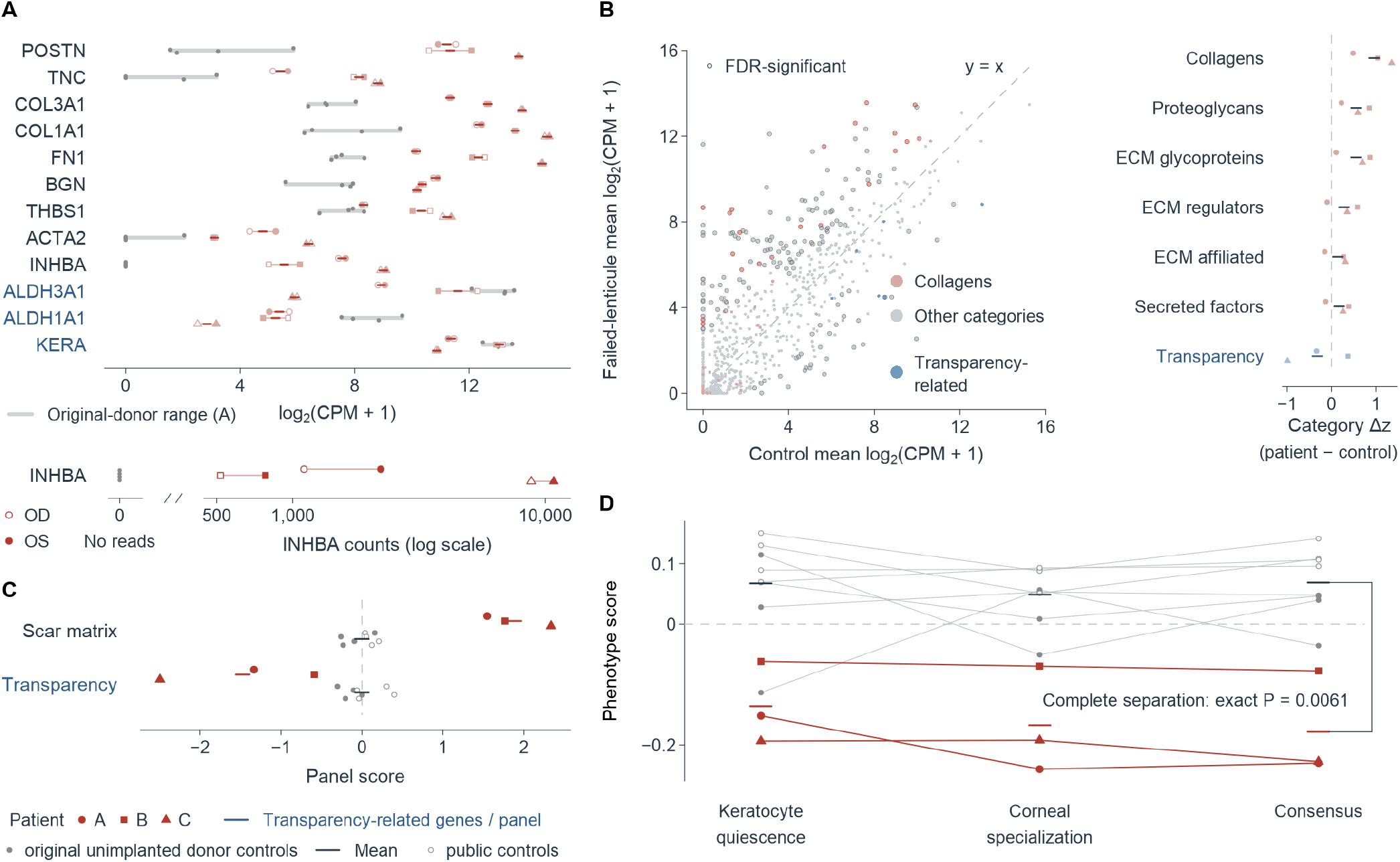
Failed lenticules express abundant scar matrix and lose the keratocyte phenotype. (A) Expression of 12 genes in six failed lenticules from three patients and four original unimplanted donor controls. Open/filled patient symbols denote right/left eyes; lines join paired eyes and short red marks show paired-eye means of log₂(CPM + 1). Gray points and bands show controls and their observed ranges. The separate INHBA track places zero counts in the “No reads” column and positive counts on a logarithmic axis. (B) Mean expression of 871 matrisome and transparency-related genes in controls versus failed lenticules. Red, blue and gray denote collagens, transparency-related genes and other categories; outlines mark patient-level false discovery rate (FDR) < 0.05. The diagonal marks equal expression. Category strips show each patient’s paired-eye mean gene z score minus the control mean and the equal-patient average; standardization uses the 12 original specimens: four donor controls and both eyes of Patients A–D, of whom Patient D supplied two lenticules removed for another indication. (C) Seven-gene scar matrix and seven-gene transparency panel scores in three patient count sums and eight controls. Gene z scores across these biological units were centered on controls and averaged; higher transparency expression is positive. (D) Phenotype scores under keratocyte-quiescence, corneal-specialization and consensus weights. Lines connect the same biological unit across weightings, and short marks show group means. The consensus bracket connects group means; the exact complete-separation test enumerates all 165 assignments of three patient labels among 11 units (P = 0.0061). In A–D, circles, squares and triangles identify Patients A, B and C. Filled/open gray points in C and D identify four original unimplanted donor controls/four public controls from PRJNA1533946. Blue gene and panel labels identify transparency-related expression. CPM, counts per million.

POSTN, TNC and COL3A1 showed large increases, with log₂ fold changes of +8.83, +6.80 and +6.00, respectively (Fig. 3A). INHBA was undetected in all four donor controls, as verified in alignments, but yielded 517–10,800 counts in failed lenticules (FDR = 4.25 × 10⁻⁴). Mean ACTA2 expression followed Patient B < Patient A < Patient C, matching interface reflectivity on the exchange day (Figs. 3A and 2B). Despite different diagnostic labels and clinical courses, all six failed lenticules showed abundant scar matrix production, that is, fibrosis.

The phenotype score measured whether tissue expression favored keratocytes or fibroblasts. Two independent human single-cell resources supplied weightings: quiescent keratocyte-like cells versus corneal fibroblasts [23], and corneal stromal fibroblasts versus fibroblasts from other ocular tissues [24]. The score represented their consensus. All three patients had lower scores than all eight controls, with complete separation (exact P = 0.0061; Fig. 3D; Supplemental Table S2). Failed lenticules thus showed abundant scar matrix production and loss of keratocyte phenotype.

### T cell, cytotoxic and interferon-γ programs were not elevated in failed lenticules

Our analysis of published pig-to-macaque corneal xenotransplant proteomics [35] showed higher scar matrix and lower transparency panel expression in four rejected grafts than in two clear surviving grafts, with complete separation by each panel (Supplemental Table S3). Scar matrix did not distinguish rejection from fibrosis, so immune programs required separate measurement.

We measured rejection using five externally defined immune gene modules [36]: T cell receptor (TCR) execution, cytotoxic T/NK activity, interferon-γ chemokine, B/plasma cell and antigen presentation. The mouse corneal allograft-versus-isograft difference defined a rejection score of +1. In an independent reference experiment, PBS-treated versus anti-CD80/86-treated mouse grafts scored +0.720 [21]. Failed lenticules scored −0.447 with featureCounts and −0.458 with Salmon; scores were negative in all three patients, and all resampling intervals for reference-minus-failure differences were positive (Fig. 4A; Supplemental Table S1).

**Figure 4.**
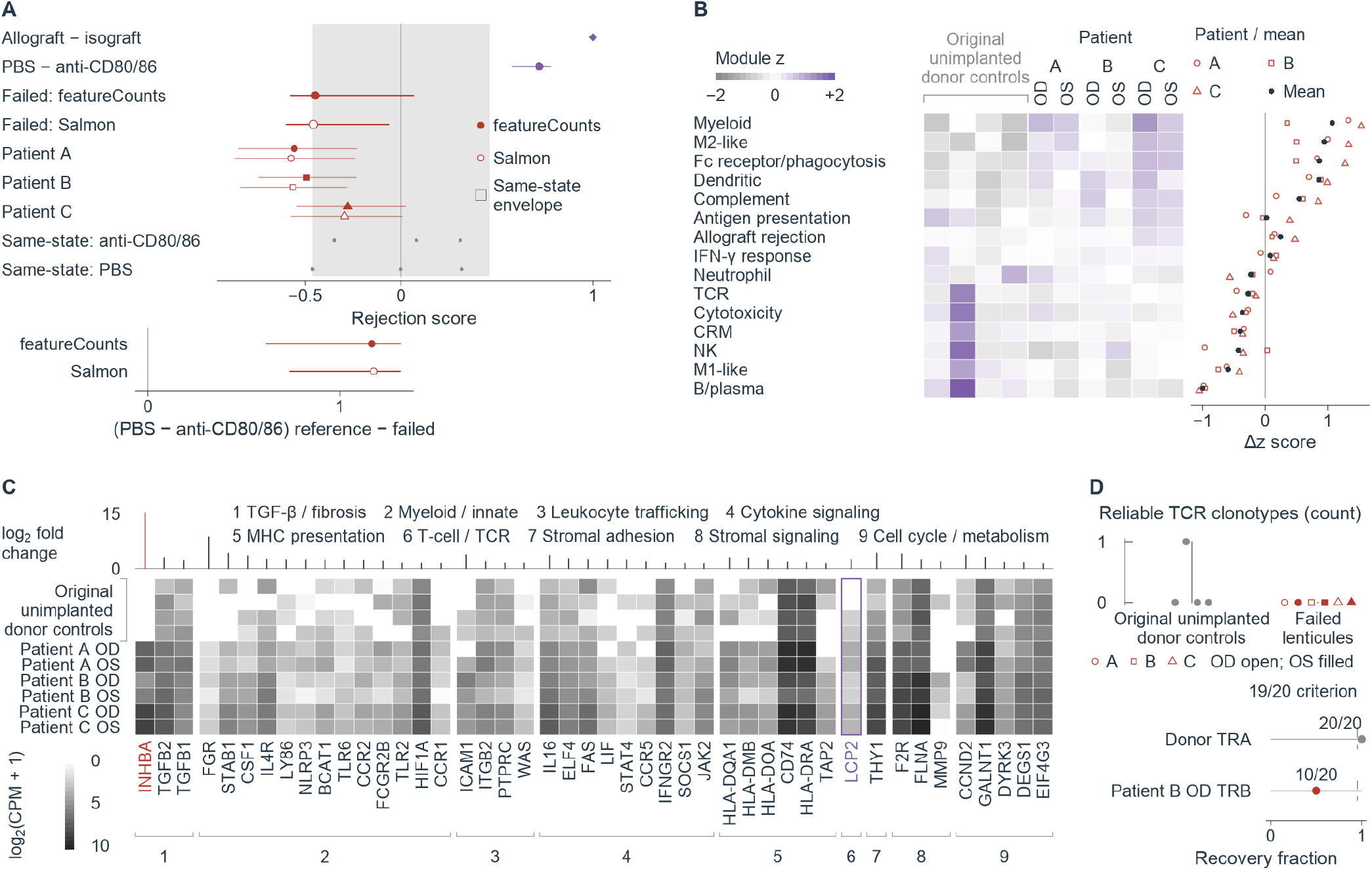
Failed lenticules lack an elevated T cell–mediated rejection program. (A) Five-module rejection scores in three patients and the patient-level comparison with four original unimplanted donor controls. Allograft minus isograft defines +1; phosphate-buffered saline (PBS) minus anti-CD80/86 is the independent positive reference. Filled/open red symbols denote featureCounts/Salmon quantification. Intervals are 95% bootstraps of reference libraries for the positive reference, reference libraries and patient/control units for group estimates and reference-minus-failed differences, and genes within modules for patient estimates. Purple symbols identify references; gray points show six same-state contrasts. Gray shading has a half-width equal to the largest absolute same-state contrast. The lower track shows the independent PBS-minus-anti-CD80/86 reference minus failed lenticules. The independent reference contains three libraries per arm; the defining reference contains one pooled library per condition. (B) Fifteen immune-program z scores in six failed lenticules and four controls, standardized across the 12 original specimens. Gray, white and purple denote −2, 0 and +2, with endpoint saturation. The adjacent strip shows bilateral patient-minus-control differences and their equal-patient mean. (C) Expression and patient-level log₂ fold changes of 45 Hallmark allograft rejection leading-edge genes, grouped into nine processes. White-to-black shading represents log₂(CPM + 1). INHBA is red; LCP2, assigned to T cell/T cell receptor (TCR) signaling, is outlined in purple. (D) Reliable TCR clonotypes in four controls and six failed lenticules; vertical segments span observed group ranges. Circles, squares and triangles identify Patients A–C; open/filled symbols denote right/left eyes. The lower track shows candidate recovery in 20 technical resamples at 16,028,223 read pairs: donor TRA, 20/20; Patient B OD TRB, 10/20; reproducibility criterion, 19/20. TRA/TRB, TCR α/β; NK, natural killer; IFN-γ, interferon-γ; CRM, common rejection module; MHC, major histocompatibility complex; TGF-β, transforming growth factor-β; Fc, fragment crystallizable; CPM, counts per million; OD/OS, right/left eye.

Myeloid, Fc receptor/phagocytosis, dendritic cell and complement programs increased in failed lenticules, whereas TCR and cytotoxic programs decreased (Fig. 4B). The interferon-γ response was not enriched by any of five scoring methods (Supplemental Fig. S3C). The broader Hallmark allograft rejection gene set [18] was enriched, with normalized enrichment score (NES) = 1.46 and adjusted P = 0.0325. INHBA ranked highest among 45 leading-edge genes; only LCP2 represented T cell/TCR signaling. The remaining genes were involved in innate immunity, cytokines, antigen presentation, adhesion, repair and metabolism (Fig. 4C), reflecting a response broader than T cell– mediated rejection. No failed lenticule exceeded 5 counts for CD3D, CD3E, CD8A, TRBC2, GZMB or IFNG (at most 0.30 CPM); GZMB and IFNG were undetected in every original specimen, whereas one donor control had 217 TRBC2 counts (Supplemental Data 1).

Receptor reconstruction gave concordant results. At matched sequencing depth, the six failed lenticules yielded no TCR clonotype meeting the prespecified reproducibility criterion, whereas one donor control yielded a reproducible TRA clonotype (Fig. 4D). Each failed lenticule had fewer reliable B cell receptor clonotypes than every donor control (Supplemental Fig. S13C). The three analyses agreed: failed lenticules lacked a T cell–mediated rejection program, and elevated immune programs were predominantly myeloid.

### Phenotype distinguished repair with restored transparency from scarring and was lower in failed lenticules than in scarred rabbit corneas

Successful repair also involved remodeling, so distinguishing recovery from failure required another measure. Remodeling activity was measured with the failure signature, comprising the top 300 genes in each direction of expression change in failed lenticules versus four donor controls. Greater similarity between a tissue-versus-control expression change and this signature denoted remodeling closer to that in failed lenticules.

Rabbit corneas healed without scarring after freeze injury [26], with backscatter falling toward normal between days 7 and 28. Failure-signature similarity increased from +3.62 to +12.99, while the phenotype score approached the uninjured level, changing from −0.0414 to −0.0146 (Fig. 5A). Rabbit corneal fibrosis on day 14 after alkali burn [27] likewise showed high failure-signature similarity (+10.15), but a much lower phenotype score (−0.1405; Fig. 5A). Remodeling was similarly active during repair and fibrosis; phenotype distinguished them.

**Figure 5.**
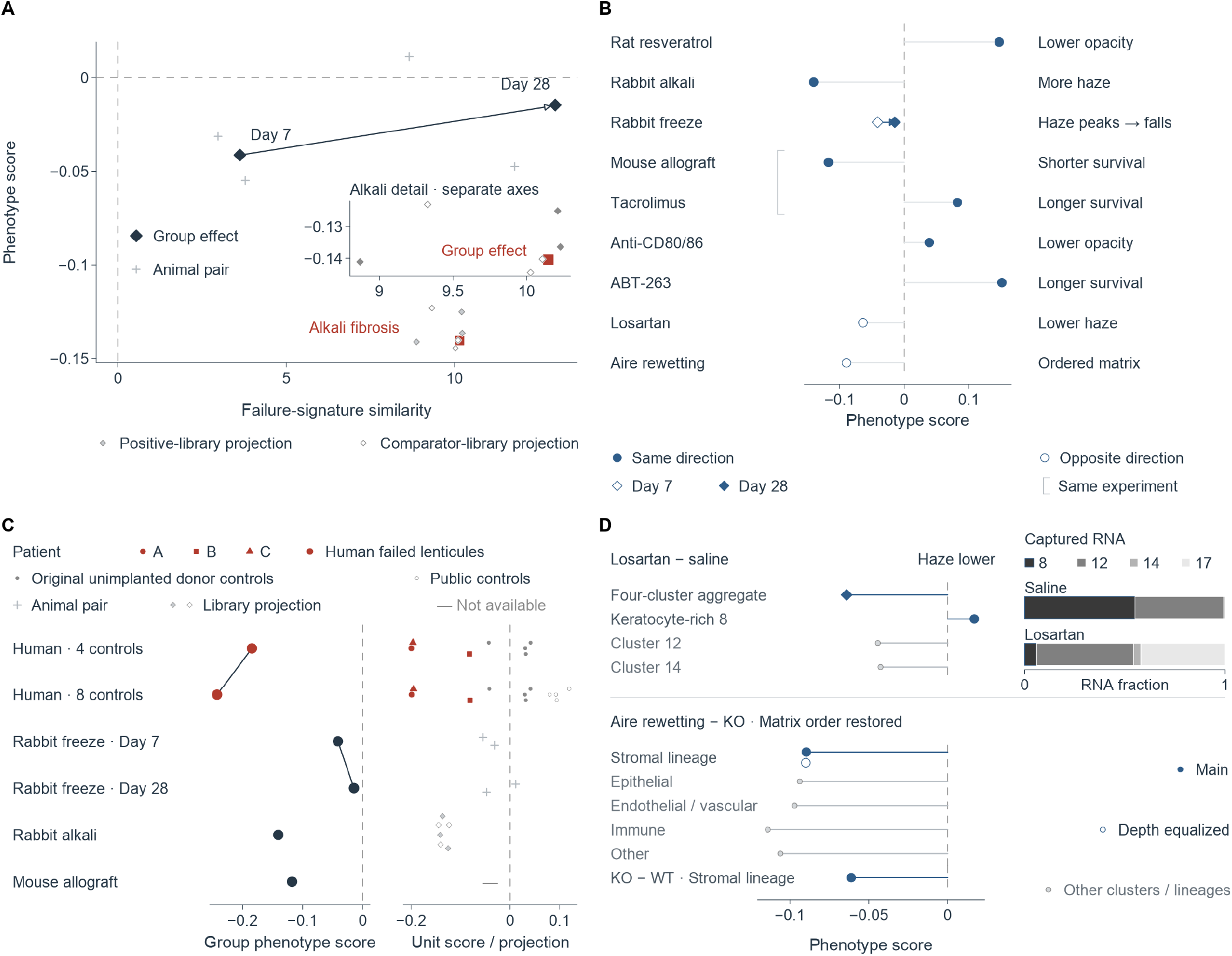
Keratocyte phenotype distinguishes clearing repair from fibrosis and diverges from matrix organization. (A) Failure-signature similarity versus phenotype score during rabbit freeze-injury repair and alkali-burn fibrosis. Large diamonds and the square show separately calculated repair and fibrosis group effects. Crosses show two injured–contralateral animal pairs per freeze-injury time; small filled/open diamonds show three fibrotic/three naive library projections. The arrow connects day-7 and day-28 estimates from different animals. The inset enlarges fibrosis on separate axes. (B) Phenotype direction and reported outcomes for ten comparisons from eight independent in vivo experiments, displayed on nine tracks. Freeze-injury times share a trajectory; brackets link comparisons from the same experiment. Filled/open circles denote agreement/disagreement between phenotype and outcome direction; open/filled diamonds denote freeze days 7/28. Six experiments have matching directions. (C) Six group-level phenotype estimates, with separately calculated patient/control scores, animal-pair scores or library projections alongside. Human comparisons use three patient count sums versus four original unimplanted donor controls or all eight controls (four original and four public controls). The mouse reference uses one pooled library per condition; a dash identifies its unavailable unit layer. Patient symbols distinguish Patients A–C; filled/open gray points distinguish original/public controls. (D) Losartan minus saline phenotype scores for a four-cluster stromal aggregate and three shared clusters; stacked bars show captured-RNA fractions for clusters 8, 12, 14 and 17. Below, Aire-deficient corneas after rewetting are compared with untreated deficient corneas across five lineages; the stromal-lineage comparison with wild type is shown separately. Filled/open stromal points denote main/depth-equalized analyses. Each condition in these two experiments has one library; Aire libraries are pooled. In A and C, positive-library projections compare each positive-arm library with the comparator mean, and comparator-library projections compare the positive mean with each comparator library. Group estimates retain their separate estimator. KO, Aire knockout; WT, wild type.

Using the same group-level estimate as in public experiments, failed lenticules had phenotype differences of −0.1843 versus four donor controls and −0.2424 versus all eight controls, below alkali-burn fibrosis (−0.1405) and far below repair with restored transparency after freeze injury (−0.0146) (Fig. 5C; Supplemental Table S4). Their phenotype was therefore lower than in scarred rabbit corneas.

We applied the same standard to eight independent public in vivo experiments reporting outcomes (Supplemental Table S4). Phenotype direction agreed with outcomes in six (Fig. 5B). Better outcomes accompanied higher phenotype scores with resveratrol in rat allografts (+0.1473; better clarity, edema and rejection index scores, longer survival) [37], tacrolimus in mouse allografts (+0.0823; longer survival) [30], anti-CD80/86 in mouse grafts (+0.0388; less opacity, greater survival) [21], and ABT-263 in mouse allografts (+0.1514; maintained transparency, longer survival than vehicle) [38] (Supplemental Fig. S17).

Worse outcomes accompanied lower scores in rabbit alkali-burn fibrosis on day 14 (−0.1405) and at peak haze on day 7 after freeze injury (−0.0414). PBS-treated mouse allografts scored −0.1176 versus isografts (allograft median survival approximately 15 days). After freeze injury, phenotype approached the uninjured state when transparency returned on day 28.

The other two experiments showed discordance. After nitrogen mustard injury, losartan-treated corneas [31] had 32% less opacity than saline controls on day 3, yet lower phenotype across four stromal clusters (−0.0641); keratocyte-rich stromal cluster 8 had higher phenotype (+0.0169; Fig. 5D; Supplemental Fig. S8). Single-nucleus RNA sequencing in Aire-deficient dry-eye mice [32] used one pooled library per condition. Stromal-lineage phenotype was lower than in wild-type mice (−0.0611). After 14 days of PBS rewetting, collagen alignment and interfibrillar spacing returned to wild-type-like values, and stromal thickness was lower than in untreated deficient mice. Yet stromal-lineage phenotype was 0.0896 lower than in untreated deficient mice (0.0899 after library-depth equalization; Fig. 5D; Supplemental Fig. S16). Whole-tissue phenotype could therefore diverge from cellular composition or matrix structure.

### Suppressing rejection or scar matrix expression did not determine phenotype recovery

In the same mouse corneal allograft experiment [30], tacrolimus and rapamycin both reduced rejection scores versus PBS (−0.792 and −1.007), but phenotype increased (+0.0823) and decreased (−0.1131), respectively (Fig. 6A). Both weightings reproduced these opposite changes (Supplemental Data 1). The original study reported that tacrolimus extended median graft survival by 11 days.

**Figure 6.**
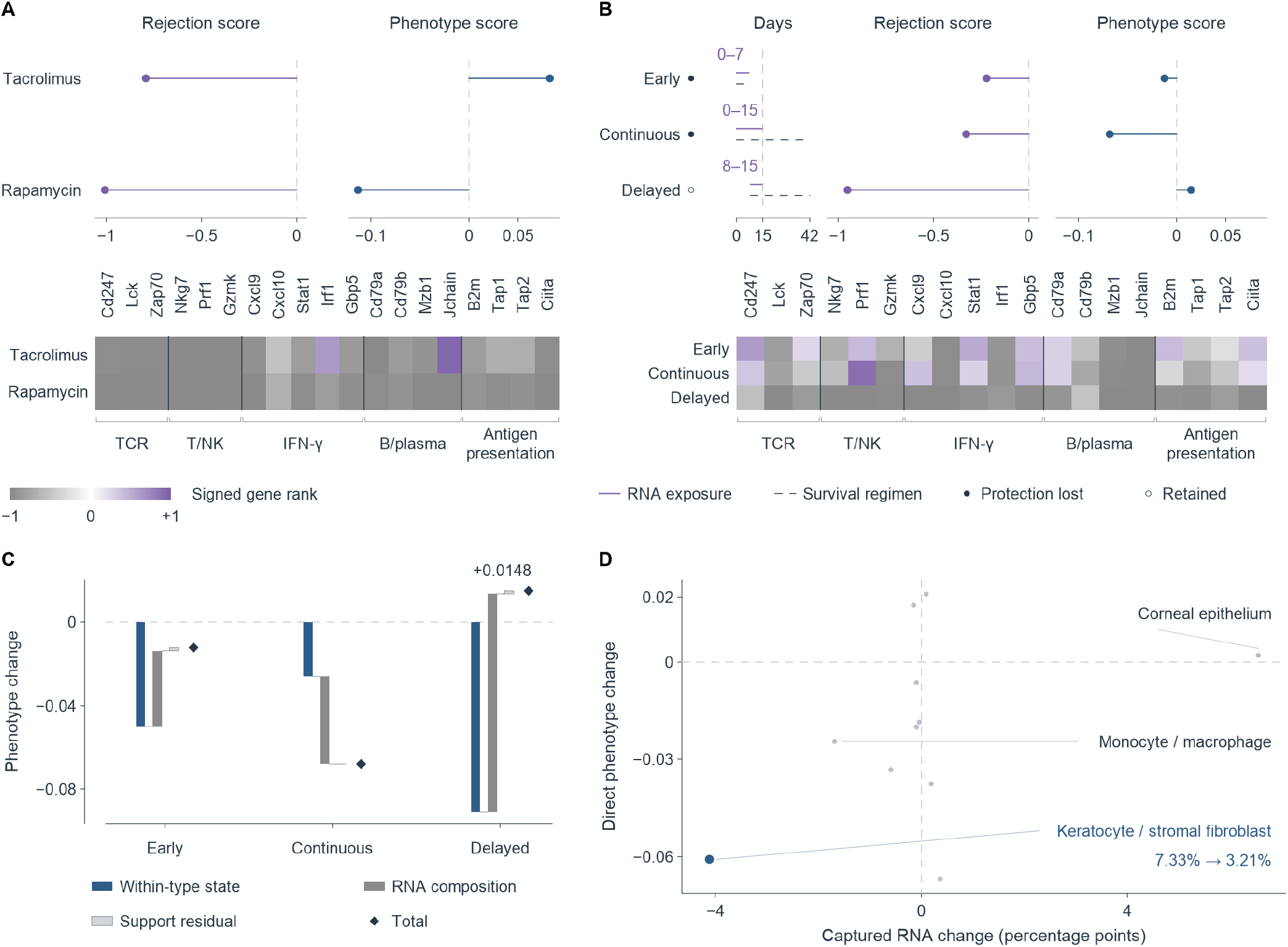
Suppressing rejection does not determine phenotype recovery, and altered RNA composition can mask declining cellular phenotype. (A) Rejection and phenotype scores for tacrolimus and rapamycin versus PBS in mouse corneal allografts. (B) Early, continuous and delayed anti-Gr-1 plus tacrolimus versus tacrolimus alone at day 15. Solid purple/dashed dark timelines show RNA-exposure/survival regimens: days 0–7/0–7, 0–15/0–42 and 8–15/8–42. The vertical line marks RNA collection on day 15. Filled/open regimen symbols denote lost/retained protection. In A and B, heatmaps show the same 19 signed gene ranks in five modules: TCR execution, cytotoxic T/NK activity, IFN-γ chemokine, B/plasma cell and antigen presentation. Gray, white and purple denote −1, 0 and +1. Each RNA condition is one pooled library from 4–6 corneas. (C) Fine-cell-type decomposition of whole-tissue phenotype change for the three depletion windows. Blue, gray and pale gray segments add within-type state, captured-RNA composition and support residual; diamonds mark totals. For delayed depletion, +0.014796 = −0.091148 + 0.104561 + 0.001383. (D) Direct within-type phenotype change versus captured-RNA-share change, in percentage points, for 11 common cell types with calculable scores under delayed depletion. The blue point identifies keratocytes/corneal stromal fibroblasts, whose RNA share fell from 7.33% to 3.21%; gray points identify the other types. Dashed lines mark zero change. PBS, phosphate-buffered saline; TCR, T cell receptor; NK, natural killer; IFN-γ, interferon-γ.

Early and continuous anti-Gr-1 depletion abolished tacrolimus-associated protection, whereas delayed depletion preserved it. Compared with tacrolimus alone on day 15, all three depletion arms had lower rejection scores, but phenotype increased only with delayed depletion and decreased with early and continuous depletion (Fig. 6B). Rejection scores fell with both lost and retained protection, whereas phenotype followed protection. Whole-tissue phenotype depended on expression state and RNA share. Its increase with delayed depletion (+0.0148) arose from RNA composition (+0.1046), while expression state within common cell types decreased (−0.0911; Fig. 6C). Within the keratocyte/corneal stromal fibroblast cell type, phenotype decreased (−0.0608) and RNA share fell from 7.33% to 3.21% (Fig. 6D; Supplemental Table S5; Supplemental Fig. S15B).

Rabbit corneal cells seeded at 50,000 or 100,000 cells per collagen matrix had much lower scar matrix expression in 3D culture in fibrillar collagen than in 2D coated culture. Mean log₂ fold changes across seven scar genes were −3.25 at 50,000 and −2.84 at 100,000 [28] (Fig. 7A). Both densities reduced failure-signature similarity and increased the seven-gene transparency panel, but phenotype increased at 50,000 (+0.0422) and decreased at 100,000 (−0.0407; Fig. 7B). Directions agreed across three libraries per condition. Comparing 100,000 with 50,000 showed phenotype lower by 0.1461, with agreement between weightings (Fig. 7C; Supplemental Fig. S7A and B). Large reductions in scar matrix expression accompanied opposite genome-wide phenotype changes. Suppressing rejection or scar matrix expression did not predict phenotype recovery, which required separate measurement (Supplemental Fig. S9).

**Figure 7.**
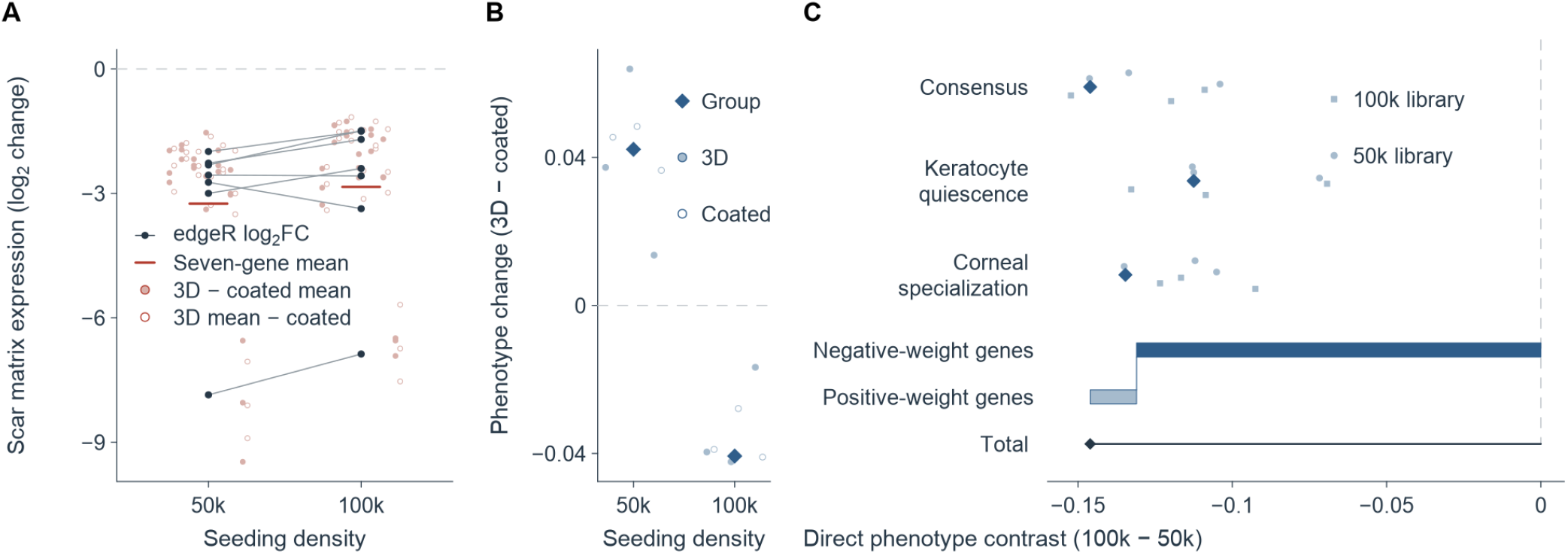
Large reductions in scar matrix accompany opposite phenotype responses at two culture densities. (A) Seven scar matrix gene effects in rabbit corneal cells cultured in three-dimensional (3D) fibrillar collagen versus two-dimensional coated culture. The labels 50k and 100k denote 50,000 and 100,000 cells seeded per collagen matrix. Dark points and connecting lines show each gene’s edgeR log₂ fold changes; red marks show seven-gene means (−3.25 and −2.84). Pale filled/open points show 3D-library minus coated-mean/3D-mean minus coated-library log₂ counts-per-million projections. The two comparisons reuse nine libraries, three per condition. (B) Phenotype group estimates and library projections for the same contrasts. Diamonds denote group estimates; filled/open points denote 3D/coated libraries. Twelve projections reuse nine libraries because the coated comparator is shared. (C) Direct 100k-minus-50k phenotype under consensus, keratocyte-quiescence and corneal-specialization weights. Diamonds show group estimates; circles and squares show the same three 50k and three 100k libraries, respectively, under each weighting. Below, dark/pale blue segments show negative-/positive-weight contributions, using a common absolute-weight denominator, and the dark diamond marks their sum. Group estimates and library projections are calculated separately. Dashed lines mark zero change.

### Drug treatment reduced the remodeling program and increased phenotype

INHBA was much higher in failed lenticules. TGF-β induction increased INHBA and SRF/MRTF target gene expression in human, rabbit and cat corneal cells [29,39,40] (Fig. 8A). Both measures also increased in rabbit alkali-burn fibrosis and declined as transparency returned during repair after freeze injury (Fig. 8A; Supplemental Table S6). INHBA and SRF/MRTF target gene expression thus measured remodeling activity.

**Figure 8.**
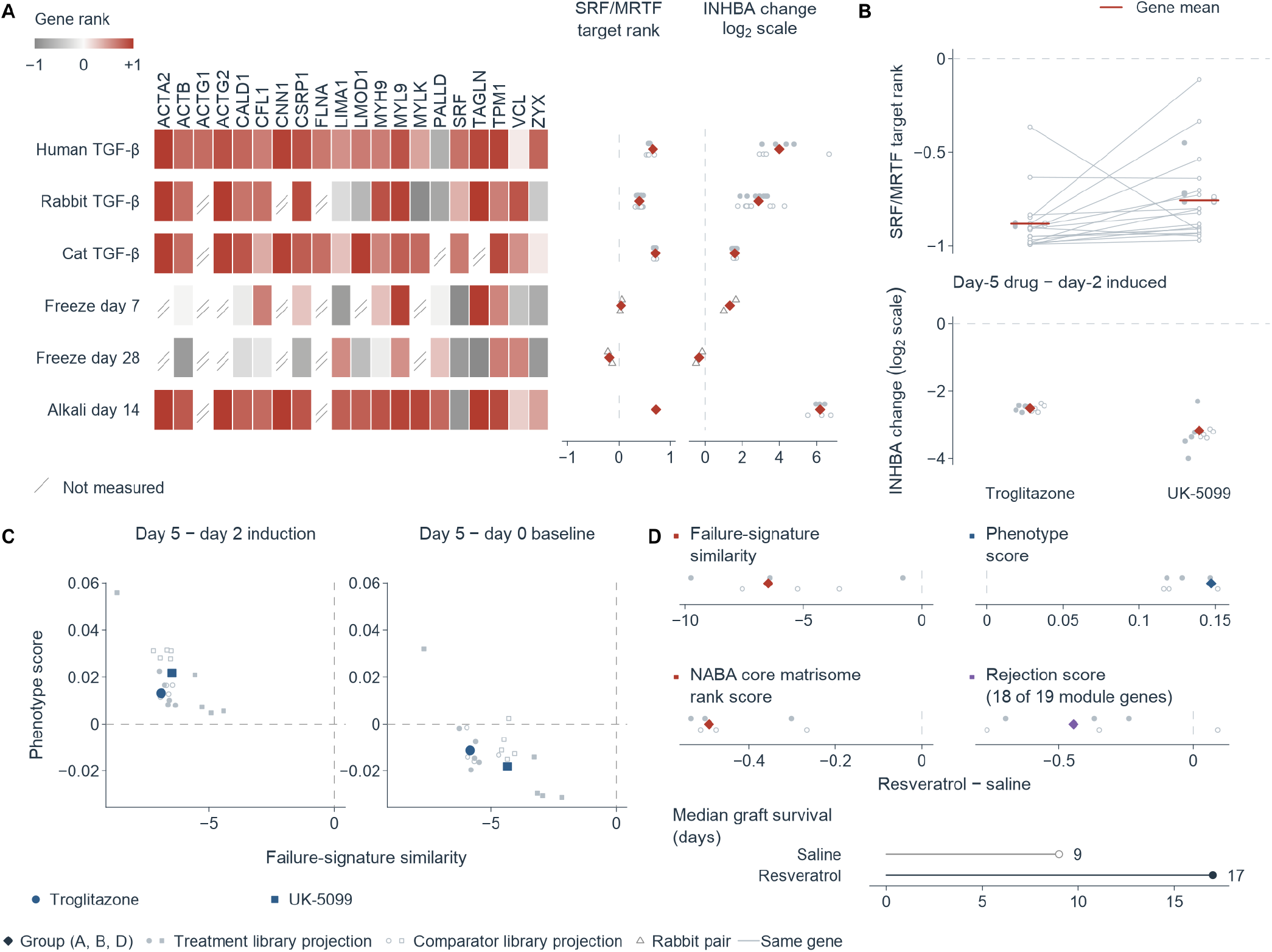
Drug treatment reduces the remodeling program and increases keratocyte phenotype. (A) INHBA expression change and 20 prespecified serum response factor/myocardin-related transcription factor (SRF/MRTF) target-gene ranks across six contexts: TGF-β induction in human, rabbit and feline corneal cells, rabbit freeze-injury repair at days 7 and 28, and day-14 alkali-burn fibrosis. Induction comparisons have 4, 8 and 5 libraries per arm, respectively; freeze-injury times have two injured–contralateral animal pairs each; fibrosis has three libraries per arm. Gray, white and red denote target ranks of −1, 0 and +1; slashes denote unmeasured genes. Adjacent tracks show mean target ranks and INHBA effects with library projections or animal-pair scores. Human INHBA uses differences in log₂(author-normalized count + 1); rabbit and feline induction group effects use edgeR log₂ fold change and library projections use Δlog₂CPM; in vivo INHBA uses Δlog₂CPM. (B) Day-5 troglitazone and UK-5099 versus day-2 induction in feline corneal cells. Lines join the same 17 measurable target genes across drugs; red marks show their mean ranks. The INHBA track retains group and library estimators. Each condition has five libraries; two comparisons generate 20 projections from 15 unique libraries. (C) Failure-signature similarity versus phenotype for both drugs, relative to day-2 induction or day-0 baseline. Large circles/squares identify troglitazone/UK-5099 group estimates; small filled/open symbols show treatment-/comparator-library projections. (D) Resveratrol versus saline in rat allografts: failure-signature similarity, phenotype, NABA core matrisome rank score and rejection score from 18 of 19 module genes. Each arm contains three corneas, one library per cornea. The NABA score averages signed effect ranks of measurable core matrisome genes; the rejection readout uses native-expression group means. The lower track reports median graft survival of 17 versus 9 days, with ten grafts per arm. In A, B and D, diamonds denote group estimates; filled/open small symbols denote treatment-library minus comparator-mean/treatment-mean minus comparator-library projections, and triangles denote rabbit pairs. Red, blue and purple identify remodeling, phenotype and rejection readouts. TGF-β, transforming growth factor-β; CPM, counts per million.

Treatment after induction tested drug responsiveness. Troglitazone or UK-5099 in cat corneal myofibroblasts [29] reduced INHBA and SRF/MRTF target expression on day 5 relative to the induced state on day 2 (Fig. 8B). Failure-signature similarity decreased (−6.907 and −6.471, respectively), and phenotype increased (+0.0132 and +0.0217) (Fig. 8C; Supplemental Fig. S14; Supplemental Table S7).

Resveratrol reduced the remodeling program and increased phenotype in rat corneal allografts [37]. Compared with saline, failure-signature similarity decreased (−6.50), core matrix gene expression decreased (−0.493), and phenotype increased (+0.1473; Fig. 8D). The rejection score from 18 available module genes also decreased (−0.445). Both weightings reproduced the phenotype increase (Supplemental Data 1). The original study reported less opacity at RNA sampling on day 7 and thinner corneas on day 14. Graft survival was longer (median 17 versus 9 days; 10 grafts per arm; Fig. 8D). Reduced rejection and remodeling with increased phenotype accompanied better graft appearance and longer survival.

## Discussion

Failed allogeneic corneal stromal lenticules expressed abundant scar matrix, whereas T cell, cytotoxic and interferon-γ programs were not increased. Opacity was therefore caused by fibrosis rather than rejection. Keratocyte phenotype distinguishes recovery from failure and was lower in failed lenticules than in all controls. Phenotype responses differed in direction among treatments suppressing rejection or scar matrix expression, showing that neither effect is sufficient to restore phenotype.

Our analysis of published pig-to-macaque corneal xenotransplant proteomic data [35] showed that rejected grafts also had high scar matrix and low transparency panel expression. Scar matrix cannot distinguish rejection from fibrosis, so the cause must be determined from immune programs. Failed lenticules had lower T cell receptor and cytotoxic programs than controls. None of the six lenticules had reliable T cell receptor clonotypes. Myeloid, phagocytosis and complement programs were increased. Among 17 human corneal allografts removed 4 weeks to approximately 20 years after rejection, 15 had chronic stromal infiltrates dominated by T lymphocytes and macrophages, whereas two further grafts replaced for reasons other than rejection had none. Infiltration was greater in grafts removed earlier and was not significantly correlated with pre-removal topical corticosteroid dose [41].

Examination records described only opacity in all three patients’ initial implants. Even if the first recorded event leading to exchange marked rejection onset, the longest interval to removal was 77 days, overlapping the period of greatest infiltration in these allografts. Failed lenticules therefore showed neither ongoing T cell–mediated rejection nor a residual T cell signature of rejection, leaving no T cell–mediated lesion for intensified immunosuppression to target.

Fibrosis involves abundant scar matrix formation as implanted tissue remodels, yet remodeling and scar matrix accompany both recovery and failure. Myofibroblast markers can be induced without crystallin loss in cultured rabbit corneal cells [42]. Rabbit corneas that regained transparency by day 28 after freeze injury [26] and those scarred after alkali burns [27] shared the failed-lenticule remodeling program. Phenotype was near uninjured levels in the former, significantly lower in the latter, and lower still in failed lenticules. Across eight independent public in vivo experiments reporting transparency, graft survival or stromal structure, phenotype direction agreed with reported outcomes in six. Phenotype therefore measures the outcome of remodeling rather than its activity, distinguishing recovery from failure. This distinction relates to stromal structure and function: after stromal opacity developed in rats, injecting quiescent keratocytes reduced opacity, restored ordered collagen fibers and improved vision, whereas injecting stromal fibroblasts did not [43]. The two discordant experiments showed that whole-tissue phenotype can diverge from cell composition or stromal structure. Losartan-treated corneas had less opacity after nitrogen mustard injury but lower phenotype across four stromal clusters combined, despite higher phenotype in the keratocyte-rich cluster [31]. Rewetting restored wild-type-like collagen organization in Aire-deficient dry-eye mice, yet stromal-lineage phenotype remained below that in untreated deficient mice [32]. Integration therefore requires assessing both phenotype and stromal organization.

Tacrolimus and rapamycin suppressed rejection in the same mouse transplantation experiment but increased and decreased phenotype, respectively [30]. In rabbit corneal cells, 3D culture in fibrillar collagen greatly reduced scar matrix gene expression at both cell densities, but phenotype rose at one density and fell at the other [28]. Neither suppressing rejection nor reducing scar matrix expression therefore predicts phenotype recovery. Phenotype must be measured separately and should itself be a treatment target.

Targeting phenotype requires drugs that control remodeling. INHBA, significantly increased in failed lenticules, encodes the βA subunit of activin A, which induces myofibroblast markers in corneal fibroblasts [44]. Expression of INHBA and SRF/MRTF target genes increased in TGF-β-induced human, rabbit and feline corneal cells and in rabbit alkali-burn fibrosis, but declined during clearing after freeze injury, providing readouts of remodeling activity. Two drugs converted feline corneal myofibroblasts to a fibroblast-like state [29], with lower expression of these genes and higher phenotype than in the induced state. In rat allografts, resveratrol reduced rejection and scar-related programs, increased phenotype and prolonged median graft survival from 9 to 17 days [37]. Failed tissue lacks a T cell–mediated rejection program, and drug treatment can reduce the remodeling program. Treatment should therefore shift from intensified immunosuppression toward controlling stromal remodeling and preserving phenotype.

Clinical diagnoses differed among the three patients, and corticosteroids did not consistently or durably reverse opacity in replacement lenticules. Opacity in the initial implants was confined to the lenticule interface and immediately adjacent anterior stroma, without endothelial loss, neovascularization or ciliary injection, consistent with the absence of tissue T cell–mediated rejection. Progressive opacity should therefore not be assumed to represent rejection based on slit-lamp appearance or clinical diagnosis. Removed lenticules should be tested directly for immune programs, scar matrix and phenotype. Interface reflectivity decreases significantly within months in successful implants [45]. It initially decreased after exchange in all three patients, before opacity recurred or keratometry continued to increase and uncorrected visual acuity declined. Early clearing after exchange does not establish integration. Integration should be assessed by sustained transparency, stable corneal shape and visual acuity, and removed tissue by phenotype.

Previous studies described failed implants structurally using light microscopy, electron microscopy and immunohistochemistry, without expression data. We used reference standards from independent public data to measure immune programs, scar matrix and phenotype, addressing rejection, fibrosis and tissue acceptance, respectively. The weights and code are publicly available, allowing the same measurements in any removed implant or intervention experiment. Three directions follow. First, applying these measurements to failed specimens from other implant types can establish whether fibrosis and phenotype loss are common processes in stromal implantation failure. Transparent implants also warrant testing: transparent monkey implants still showed disordered fibers and fewer keratocytes after two years [46]. Second, implantation models should test inhibition of INHBA/activin and SRF/MRTF pathways and resveratrol, with phenotype as the primary endpoint. Third, cellular and spatial analyses should identify which cells lose phenotype, because whole-tissue measurements combine cell states with their RNA shares. Increased whole-tissue phenotype after delayed anti-Gr-1 depletion arose from altered RNA shares, while phenotype within the keratocyte cell type declined.

Recovery or failure of implanted stroma depends on whether remodeling produces an ordered matrix and cells return to a keratocyte phenotype. Integration is defined in tissue by ordered stroma and keratocyte phenotype, and confirmed clinically by sustained transparency, corneal shape and visual acuity. Evaluating implanted tissue by phenotype gives a common basis for explaining failure, designing treatment and comparing efficacy, with the goal of a persistently transparent cornea with stable shape for patients undergoing stromal implantation.

## Declarations

### Data availability

Gene-level count and normalized expression matrices, analysis code, figure source data and derived tables are available in Zenodo (https://doi.org/10.5281/zenodo.24127731). The main external datasets are listed in Supplemental Table S8, and the intervention, panel, phenotype-test and decomposition results are supplied in Supplemental Tables S1, S2, S5, S7 and S9 and Supplemental Data 1. Raw sequencing reads, clinical records and anterior-segment images contain patient information and are not publicly available; deidentified data can be obtained from the corresponding authors on reasonable request.

### Author contributions

HJ, YJ, YL, and YZ conceived and designed the study. YZ and CL performed clinical care and acquired the explanted lenticules and clinical records. YD arranged the sequencing of the specimens and acquired the sequencing data. HJ analyzed the data, prepared the figures, and wrote the original draft. HJ, YD, CL, YJ, YL, and YZ interpreted the data and reviewed and edited the manuscript. YJ, YL, and YZ supervised the study. The order of the co-first authors (HJ, YD, and CL) was determined by their relative contributions to the work.

### Declaration of generative AI use

The authors used Claude 5 series (Anthropic) and Codex GPT-6.1 series (OpenAI) from July to November 2026 to improve the language of the manuscript and to assist with writing analysis code. The authors assume full responsibility for ensuring the integrity and accuracy of AI output presented in this article.

### Declaration of competing interest

The authors have declared that no conflict of interest exists.

## Supporting information

Supporting Data Values

## Acknowledgments

The authors thank the patients whose explanted tissue made this study possible.

## Notes

### Competing Interest Statement

The authors have declared no competing interest.

### Summary of Updates

The title, abstract, analyses, figures and supplementary materials are revised relative to the first version. The specimens are unchanged: six explanted allogeneic corneal lenticules from three patients, compared with eight unimplanted donor lenticule controls. The title and abstract now state the central result directly. Failed lenticules express abundant scar matrix and have lower keratocyte phenotype scores than all controls, while T cell, cytotoxic and interferon gamma programs are not elevated, so the opacity reflects fibrosis and not rejection. The term corneal identity used in the first version is replaced throughout by keratocyte phenotype, defined as whether keratocyte characteristic gene expression is preserved. Rejection and keratocyte phenotype are read against reference standards taken from independent public data. The same standards are applied to published experiments of rabbit corneal repair, cultured corneal cells, mouse corneal grafts treated with tacrolimus or rapamycin, and rat corneal grafts treated with resveratrol. These analyses show that phenotype separates recovery from failure and that suppressing rejection does not by itself restore phenotype. The introduction now documents, from 284 published reports of corneal stromal tissue implantation, that drugs used for opacity were limited to corticosteroids and immunosuppressants and that none of 346 reported removed or biopsied specimens had gene expression analyzed. Reports were read independently by two readers with adjudication by a third. The paper now contains eight main figures and 22 supplemental figures. Supplemental tables are numbered in order of first citation. Supplemental Data 1 and the Supporting Data Values workbook give the values behind the tables and figures. The Data availability statement points to the updated Zenodo record with gene level matrices and code.

