## Supplemental Information for "Failed allogeneic corneal lenticules show fibrosis and loss of keratocyte phenotype, not rejection"

### Supplemental methods

#### Quantitative anterior-segment OCT

Serial anterior-segment OCT was acquired at 22 bilateral visits, giving 44 single-eye scans on an Optovue RTVue XR using the bilateral Cornea Line OU protocol with an 8.00-mm scan size. Reflectivity was summarized within each scan as the ratio of mean signal intensity between two fixed depth intervals in the same eye: 50–300 µm, which spans the lenticule and the implant interface, over 400–600 µm, which lies in the posterior stroma below it. The patient-level value was the mean of the two eyes' unrounded ratios. Because the ratio is referenced internally within a single scan, it reduces sensitivity to multiplicative differences in overall signal gain between visits. Measured central corneal thickness across the 44 scans ranged from 654 to 901 µm, so the 400–600 µm window serving as the denominator lay entirely within the stroma in every scan. Forty-three of the 44 scans passed the prespecified quality gate; the single exception, one eye of Patient A at a visit that was not the most recent scan, failed on corneal thickness falling outside the 450–900 µm acceptance window. Visit schedules differed between patients, and the magnitude of change is reported within each patient.

#### Scheimpflug corneal densitometry

Layered corneal densitometry was exported from a Pentacam system running software v1.26r28 on the instrument's native 0–100 grayscale scale, where 0 denotes maximum transparency and 100 maximum opacity; the export header labels this column %, and the same scale is conventionally reported as grayscale units. No unit conversion was applied. The instrument reports four depth measurements, three depth layers (the anterior 120 µm, the posterior 60 µm, and the intervening mid-stroma) and the full thickness, across four concentric annular zones (0–2, 2–6, 6–10 and 10–12 mm) and the full diameter. This gives 20 values per scan. The layered comparisons reported here use the central 0–2 mm zone as the primary estimand. Repeated scans were averaged within an eye, the two eyes were then averaged within a patient, and patients were weighted equally; we computed no eye-level statistic and ran no test at eye level. Point-value exports and matrix-average exports are different estimands and were never pooled.

Three features of these data bear on their interpretation. First, the failure-state scans were obtained on the day of exchange, immediately before surgery, in Patients B and C, but 77 days before exchange in Patient A, so the densitometry is not contemporaneous with the transcriptome in that patient. Second, the preimplantation reference is the patient-equal-weight mean of preimplant scans from two patients of this series, with replicate scans averaged within eye and then OD and OS averaged within patient. It is not an age-matched normative database, so all comparisons are within-cohort and no layer is described as normal in absolute terms. Third, the instrument's quality specification flag did not return OK for the failure-state scans, which is consistent with the reduced transparency being measured; these scans are retained and the flag is reported rather than used as an exclusion criterion.

#### Corneal topography and recorded uncorrected visual acuity

Uncorrected visual acuity was transcribed from the existing clinical record on its decimal scale, with OD and OS retained separately and visits expressed in days after exchange. For Patient A, TMS videokeratoscopy tracked corneal shape over the interval in which the clinical record contained no further entry describing the implant. Mean keratometry is reported for each eye at the available acquisition dates without interpolation.

#### Specular microscopy

Endothelial cell density and morphology were read from four bilateral specular microscopy reports acquired on NIDEK instruments. Density and morphology indices were assessed separately against the reference ranges printed on each report.

#### Anterior segment photography

One hundred and seven anterior segment photographs were available (43, 41 and 23 for Patients A, B and C). Of these, 41 had previously been reviewed for signs of rejection and were scored for corneal neovascularization and ciliary injection; those scores are reported over the 41 reviewed photographs.

#### Pathway and module scoring

Hallmark pathway-overview enrichment used fgsea [1] with the multilevel method and eps = 0, which permits estimation of very small P values, on a ranked list built from the patient-level DESeq2 Wald statistic [2]. That ranking is used for the pathway overview only. Immune-module enrichment used fgsea with the same multilevel settings and a minimum set size of three on the patient-level edgeR [3,4] log fold-change ranking. Benjamini–Hochberg [5] adjustment ran jointly over the 13 gene sets that retained at least three ranked genes, eleven immune modules and the M1-like and M2-like polarization components; the cytotoxicity, natural-killer and IL-17/Th17 modules retained zero, one and one ranked gene and were scored without an enrichment test. For Figure 4C, the leading-edge annotation and enrichment statistics were taken from the same Hallmark analysis, with BH adjustment across the Hallmark pathways. The pathway-overview, immune-panel and leading-edge annotation analyses each use their own adjustment family. Signature comparisons within the state-space analyses instead used fgseaSimple, whose P values are bounded by the permutation count and are therefore not reported at extreme magnitudes. We scored immune, matrisome and transparency programs as gene modules prespecified before the present analyses. The broader corneal transparency and keratocyte phenotype module used in the category calibration comprises thirteen genes selected from primary literature; it is distinct from the seven-gene crystallin/transparency panel used for patient-level testing. The panel uses independent patient/donor units, while the broader module summaries below use the specified specimen or cell units; extracellular matrix categories follow the NABA gene sets [6] retrieved through msigdbr. For bulk data, mitochondrial rows were removed, counts were summed by gene symbol and converted to log2(CPM + 1); within each module, genes detected and variable across samples were z-scored across samples and averaged. For external single-cell references, we computed module activity with UCell [7] on within-cell raw-count ranks and averaged by cell type and condition; pseudobulk scoring instead aggregated raw counts by sample and cell type before averaging gene percentile ranks of log2(CPM + 1). The two single-cell scoring conventions are used descriptively and compared only within a module across conditions, never between modules. For each module, we compared the observed failure-minus-control difference against an expression-matched null distribution (2,000 draws per module), and matching quality was reported as the standardized mean difference between each drawn null set and its target (Supplemental Data 1).

#### Pathogen screening

Non-host reads were classified with Kraken2 [8] v2.17.1 against the PlusPF database (2026-06-26 build) after host subtraction. An empirical stability floor was established by downsampling real read pairs, not by constructing or spiking any sequence. A taxon had to appear in at least 9 of 10 resamplings and to hold a top-10 rank in at least 9 of 10. The conservative floor was 5,158 read pairs (Figure S10A). No between-group test is reported for this screen; it assesses signal stability within specimens and recurrence across specimens.

#### External reference systems and state-space projection

External reference states were assembled from published datasets and are listed in full in Supplemental Table S8. For each dataset the table gives the accession, species, the exact contrast used here, and the number of biological or sequencing units together with what those units are (libraries, pools, animals, donors, culture replicates or matched pairs), so that single-cell counts are never read as biological replicates.

We used three groups of references. Corneal allograft rejection was represented by two murine datasets, one supplying the primary rejection axis and one providing directional calibration together with an isograft arm in which surgery occurs without allogeneic mismatch. Injury and repair were represented by a murine corneal time course, which also defines the injury–repair corridor against which orthogonal deviation was measured, by a rabbit scar-free repair model and by a rabbit alkali-burn fibrosis model. Material-associated remodeling was represented by a murine implant dataset contrasting a synthetic with a biologically derived scaffold. Human references supplied a TGF-β1-driven corneal myofibroblast state, two independently prespecified phenotype weightings derived from separate donor panels, single-cell and single-nucleus references for composition modeling, and a spatial transcriptomic dataset used to localize the failure signature within corneal tissue.

Cross-species comparisons were made on one-to-one orthologs at the level of ranked program scores. Expression matrices from different species or platforms were never merged.

Human-to-mouse mapping used strictly one-to-one orthologs (babelgene, min_support = 3, top = FALSE). Expression matrices were never concatenated and we applied no cross-study batch correction; each external system contributed only a within-study signed effect, which was converted to a genome-wide continuous rank before projection. Axis scores are therefore comparisons of the direction of state change, not of absolute expression across platforms.

#### Genome-wide keratocyte phenotype weights

Phenotype weights were prespecified from the integrated human cornea meta-atlas [9] and GSE199013 [10] before application. The meta-atlas contrast was quiescent keratocyte-like cells versus corneal fibroblasts. The GSE199013 contrast was corneal K_Fibro cells versus fibroblasts from other ocular tissues. Donor pseudobulks required at least ten cells in each compared class. Within-donor $\log_2$ CPM differences were averaged across donors, standardized within source and averaged over 12,526 shared genes. The original source weightings and their consensus were retained. The in-house data were used to apply these prespecified weights and to match baseline abundance for the random-set null.

Robustness was assessed by deleting each of 1,622 HGNC gene families in turn, by deleting the top one to three contributing genes, and against 1,000 random gene sets matched on external expression, gene length and in-house control abundance quartile.

The costimulation-blockade-sensitive allorejection axis (A1) was defined by the GSE181893 PBS-minus-anti-CD80/86 contrast and directionally calibrated against the GSE275249 allograft-plus-PBS-minus-isograft contrast. The resolution axis (A2) combined the GSE303805 all-cell D2-minus-Day40, stromal D5-minus-Day40 and stromal D10-minus-Day40 contrasts. Only genes with the same sign in all component contrasts were retained, and their component-specific rank effects were averaged. The implant-remodeling axis (A3) was constructed analogously from PCL-minus-ECM effects in the CD45e_old_1wk, CD45e_young_1wk and Fibroblast_young_1wk strata of GSE175889. A1 used its primary single-contrast readout, whereas A2 and A3 used their respective same-direction consensus readouts.

Each axis and query contrast was converted to a continuous within-contrast gene rank. At N = 300 per signed arm, the forward projection scored the axis's positive and negative genes in the query rank, and the reciprocal projection scored the query's positive and negative genes in the axis rank. Each signed mean was divided by its rank-permutation null standard deviation. The symmetric score was the arithmetic mean of the forward and reciprocal normalized enrichment scores. Sample scores were centered by subtracting the mean score of the four donor controls.

Axis non-redundancy was tested in the 1,431 genes shared by all primary axes. Each axis-weight vector was standardized across this common gene space and regressed by ordinary least squares, with an intercept, on the other two standardized axes. The residual weights defined the mutually residualized axes and were projected with the same symmetric N = 300 procedure. Model $R^2$ was the ordinary coefficient of determination from the corresponding gene-level regression. A patient score was the median of the paired-eye scores. The patient-level separation margin was minimum patient score - maximum donor-control score.

The random-axis null comprised 1,000 iterations in the same common-gene space. Within each of ten in-house logCPM strata, the observed weights of each raw external axis were permuted, preserving the genes and the complete weight distribution. Each random triplet was mutually residualized and projected as above. The null statistic was the sample-level separation gap, minimum failed-sample score - maximum control score, rather than the patient-level margin. Its upper-tail empirical P value was (1 + number of null gaps at least as large as the observed gap) / 1,001.

For each state, the core matrix expression score was the mean continuous ranked effect over NABA core matrisome genes. The phenotype score was the weighted inner product of the same ranked effect and the externally prespecified consensus phenotype weights, divided by the sum of the available absolute weights. Core matrix expression and the phenotype score were standardized separately against the GSE303805

[11] stromal D2, D5, D10 and D40 reference states. The reported matrix-mis-specification statistic was core matrix expression z - phenotype score z; it was therefore a signed difference between standardized scores, not a geometric distance or a separately fitted score.

The injury–repair trajectory was the chronological polyline joining those four stromal reference states in the core-matrix-expression-by-phenotype plane. Orthogonal deviation was the shortest Euclidean distance from a state to this polyline. Corridor width was the larger held-out interpolation error obtained by omitting D5 or D10 and projecting the omitted state onto the polyline through the other anchors. For the matched-set test, high-information phenotype genes were replaced by genes matched on external atlas abundance, gene length and in-house control baseline abundance. The prespecified high-information weights were transferred to the matched genes before the phenotype score and polyline distance were recomputed. The upper-tail P value for orthogonal deviation used (1 + number of matched-null distances at least as large as observed) / (B + 1), where B was the number of matched random sets.

#### Cell-composition deconvolution

Bulk composition was estimated against two independent human corneal single-cell references [10,12] using two methods (BayesPrism [13] and donor-aware MuSiC [14]), giving four reference–method pipelines. Estimates are RNA contribution fractions, not histological cell counts. We measured reliability directly: per-sample reconstruction $R^2$, pairwise Jensen–Shannon distance between pipelines, and a signature-separability gate for distinguishing activated keratocytes from myofibroblast-like cells. A direction was reported only if it was concordant across both methods and both references and if both references supplied at least 25 cells from at least 2 donors for that class.

The bulk contrast was decomposed as observed effect = reference-mixture composition effect + residual, which holds exactly per sample. Because the residual absorbs unmodeled composition as well as genuine within-lineage state, results are reported as robustness to composition adjustment, not as cell-intrinsic attribution.

#### Spatial analysis

External human spatial data [15] were analyzed by spot-level deconvolution using spacexr RCTD [16] v2.2.1 in full mode with a minimum of 100 UMIs per spot, against the same two human corneal references. Adjacency was evaluated on the Visium six-neighbor lattice with 999 permutations that preserve the composition of each in-tissue connected component, and inference was performed with each cornea as an equal-weight independent unit: spot-level fits were summarized per sample before sign-flip or label permutation. Empirical P values from 999 permutations have a floor of 0.0010 and are reported at that floor.

The failure signature was taken from the patient-level failure-versus-control edgeR contrast. Among genes detected in the spatial dataset, the 200 most positive and 200 most negative log-fold changes were scored separately by UCell. The signed failure signature score for each spot was defined as (up-score - down-score) / 2. Epithelial, keratocyte/stromal and corneal endothelial marker programs were fixed before localization. Within each cornea, Spearman correlation was calculated between the signed failure score and each marker-program score. Corneas, rather than spots, were the units of inference: the per-cornea correlations were averaged with equal weight across the 13 corneas and tested against all sample-level sign assignments. Benjamini–Hochberg adjustment was applied across the marker programs within the correlation family.

The deconvolution-based spatial sensitivity analysis used macrophage/monocyte abundance as the index field. Activated-stromal abundance was the sum of the activated-stroma, activated-keratocyte and myofibroblast-candidate RCTD weights. High-abundance spots were defined within each cornea by the top 20% of each field. Four spatial metrics were evaluated. Inclusive adjacency was the fraction of directed edges from macrophage-high spots that ended at an activated-stroma-high spot, minus its within-cornea permutation mean. Strict adjacency applied the same calculation after excluding spots high for both fields from both endpoint classes. Residual cross-Moran correlation was the mean product of standardized residual abundances across directed neighboring spots after each abundance had been residualized against library size and the epithelial, stromal and endothelial marker percentiles. The conditional-neighbor statistic was the standardized regression coefficient for index-spot macrophage abundance in a model of mean neighboring activated-stromal abundance, adjusted for same-spot activated-stromal abundance, library size and the three marker percentiles.

Thresholded adjacency nulls used 999 permutations within each cornea, connected component and dominant marker-defined layer. Each spatial metric was then summarized as the equal-cornea mean and tested by exact sample-level sign flipping. Benjamini–Hochberg adjustment was applied within each reference across the four spatial metrics. A metric passed the cross-reference consistency gate only when its mean was positive and its adjusted P value was below 0.05 under both human corneal references. For either thresholded adjacency metric, attenuation after deconvolution was defined as 100 × (1 - RCTD effect / gene-score effect). Attenuation was summarized as the range of this quantity across the two references for the inclusive metric.

These analyses localized a transcriptomic signature and tested program neighborhoods in external human corneal tissue. They were reference-localization analyses, not measurements of cellular composition in the failed lenticules, and spot-level co-occurrence was not treated as same-cell expression.

#### Clinical-record confidence grading

Diagnostic terms, treatment and follow-up were transcribed directly from the primary clinical record: 86 scanned clinical and examination documents (31, 19, 20 and 16 for Patients A, B, C and D) and a 132-column clinical spreadsheet. Because much of this record is handwritten, every extracted item carries a confidence grade, and items that could not be read reliably were recorded as not retrievable. Diagnostic wording was translated verbatim and not harmonized into a single term. Surgical dates were derived. Two interval conventions are used. The primary convention runs from implantation to explantation, which places the three patients on a common scale for time in situ. Where post-exchange observation is compared against the course of the first lenticule, the comparator runs from implantation to the first documented event that led to exchange. In Patient C an earlier entry describing implant opacity appears at 36 days, and in Patient B the first documented event was the exchange visit itself. Formal operative notes for the three exchange procedures were not located, so lenticule dimensions and interface handling are reported as not retrievable.

Patient B's planned preoperative correction was +8.00 D in the right eye and +8.25 D in the left eye.

Sample-to-patient correspondence of the sequenced specimens was confirmed against the submission record and the provider's sample-receipt dates; each patient contributed both eyes.

Surgical dates were taken from operative signatures where these exist and otherwise from the date of the corresponding first-postoperative-day entry, corroborated against the specimen collection date.

Printed fields and machine-generated timestamps were treated as high confidence, as were handwritten dates corroborated by a second independent entry. General handwritten diagnostic and medication entries were treated as medium confidence. Items that could not be read reliably were recorded as not retrievable rather than inferred. Preservation status was established from the operating team and is not among the missing variables.

#### Transcriptome-wide convergence analysis

Convergence of the failed lenticules was measured on a fixed core feature set of 17,605 genes (mitochondrial rows removed, counts aggregated by gene symbol, log2(CPM+1), genes retained at CPM ≥ 1 in at least two of the ten primary specimens). The primary metric was the genome-wide Pearson correlation distance $d = 1 - r$; Spearman distance was computed as a same-direction sensitivity only. The convergence statistic was $\Delta$ = mean d(failure, control) − mean d(failure, failure), so $\Delta > 0$ indicates that failed lenticules lie closer to one another than to controls. A technically adjusted view regressed each gene on logit(mitochondrial read fraction) and log(Shannon effective genes), fitted on the ten primary specimens without protecting group labels, a conservative correction that removes any biological signal collinear with these covariates. The two Patient D specimens were projected onto the fixed feature set and excluded from gene selection, model fitting and all group-level statistics. Significance used the complete label-permutation space: all C(10, 6) = 210 assignments of the ten primary specimens exactly enumerated per view, with 10,000 Monte Carlo draws as a check, and the exact upper-tail P reported as (rank of the observed $\Delta$)/210. To remove the paired-eye contribution, the analysis was repeated for all eight one-eye-per-patient choices, each with its complete C(7, 3) = 35 exact permutation space.

#### Count reproducibility and orthogonal quantification

Libraries were unstranded: Salmon's automatic library-type detection returned IU for all twelve samples, with compatible-fragment ratios of 1.0 and strand-mapping bias between 0.491 and 0.526. Sequencing consumed all available tissue, leaving no blocks, sections, residual RNA or cDNA libraries.

The ten-gene glucocorticoid-receptor activation fingerprint comprised *FKBP5*, *TSC22D3*, *PER1*, *KLF15*, *ZBTB16*, *DDIT4*, *SESN1*, *CEBPD*, *ANGPTL4* and *TXNIP*. The receptor gene *NR3C1* was evaluated separately.

Count reproducibility was assessed by independent recomputation before any biological analysis. A like-for-like featureCounts [17] recomputation (v2.1.1; -p --countReadPairs -B -Q 10 -s 0 -t exon -g gene_id) met the prespecified reproducibility threshold of per-sample Spearman and log1p-Pearson ≥ 0.995, with total-count ratios within 0.98–1.02. An orthogonal Salmon v1.4.0 [18] quantification (index -k 31 --keepDuplicates; quant -l A --validateMappings --seqBias --gcBias) was run against a transcriptome built from the same genome and GTF; agreement was lower, consistent with a different assignment model. Library-complexity and composition metrics that failed quality thresholds were retained at their observed values.

An eye-level differential-expression analysis was run only as a sensitivity check. It gave essentially identical effect sizes (genome-wide Spearman $\rho = 0.9995$ across 15,338 matched genes against the patient-level result) but roughly twice as many genes below FDR 0.05 (2,144 versus 1,055), consistent with the inflated power expected from treating paired eyes as independent. Of the 1,055 patient-level significant genes, 1,039 were also tested at eye level and all 1,039 were significant there.

#### Clonotype reliability gate (full criteria)

A clonotype was counted as reliable only if it satisfied a six-part gate: (i) a complete CDR3 nucleotide sequence over the canonical alphabet with a TRUST4 [19] CDR3 score above 0.01 and canonical start and end codons; (ii) a productive, in-frame rearrangement translating without stop or ambiguous residues; (iii) chain-consistent V and J assignments carrying the prefixes expected for that locus, each with non-zero identity; (iv) support from at least two independent read fragments, counted after TRUST4 merges paired mates; (v) recovery in at least 19 of 20 common-depth resampling replicates; and (vi) recovery of the same chain and CDR3 nucleotide sequence when the same sample was reconstructed from aligned BAM input.

Original unimplanted donor control 1 defined the common-depth floor: its native depth equaled the target of 16,028,223 read pairs. It was therefore not subsampled, and all 20 of its runs contained the same 41,054 selected candidate read pairs. Criterion (v) is consequently trivially satisfied for that specimen and is not interpreted as independent resampling evidence for it; its reliable calls additionally required criteria (i) to (iv) and recovery of the same chain and CDR3 nucleotide sequence from BAM input. Common-depth repertoire reconstruction was performed separately from the count-level depth-sensitivity analysis.

#### Patient-level optical-matrix panels

The same 35 label allocations were used for all panels in the original cohort. The prespecified ordering statistic was the smaller of the scar-minus-SLRP and SLRP-minus-transparency differences. The three individual panel tests and this joint ordering test were reported separately. KERA reassignment, DCN deletion and leave-one-patient-out calculations were carried out as sensitivity analyses; the prespecified three-level ordering was not established. Salmon was an alternative quantification of the same specimens.

#### Public perturbation readouts

The failure signature comprises the 300 most upregulated and 300 most downregulated genes in the patient-level contrast of failed lenticules versus the four donor controls. Positive failure-signature similarity indicates concordant gene-expression changes with that contrast. The rejection score used five immune modules and was scaled so that mouse allograft PBS minus isograft equaled +1.

Each public comparison retained its processed gene effects and original biological design. Failure-signature similarity and A1–A3 scores used the prespecified symmetric signed-rank procedure at $N = 300$; $N = 50$, 100, 200 and 500 results remain in the complete table. The phenotype score was the weighted inner product of ranked effects and prespecified external phenotype weights, normalized by the sum of absolute weights. Core matrix expression used the core-matrisome list. Twenty-eight comparisons had all 19 members of the rejection score; 41 lacked at least one member. Complete projections and available-member subsets occupy separate columns. Forty-three comparisons had at least two libraries in both arms. Library-minus-opposite-arm-mean projections were summarized separately for the positive and negative arms. Mechanism readouts retained the source effect scale for INHBA, FST and FSTL3, and the existing 20-gene SRF/MRTF target set was summarized by its within-comparison mean rank. The accession, contrast, experimental time, biological or pooled-library unit, member coverage and original projection P values accompany every row.

For the group-level SRF/MRTF target readout, ranks were scaled as $(r - (G + 1)/2)/((G - 1)/2)$, where r is the average-tie gene rank and G is the number of ranked genes, and averaged across available members of the prespecified 20-gene set. Per-library target readouts used $2r/(G + 1) - 1$, retaining their original estimator. INHBA, FST and FSTL3 retained the original gene-effect scale of each source contrast. Supplemental Table S6 assembles the existing day-7 and day-28 freeze-injury contrasts and day-14 alkali-burn contrast with their corresponding phenotype scores, core matrix expression and remodeling readouts.

#### Original-scale effects in the culture comparisons

The rabbit GSE337743 and feline GSE231796 culture comparisons retained the original edgeR filtering, TMM normalization, dispersion estimation and quasi-likelihood models. The rabbit experiment contained 15 libraries in five conditions and the feline experiment 25 libraries in five conditions; the seven and ten original pairwise comparisons were recovered, respectively. Original model $\log_2$ fold changes, FDR values, gene availability and rank-based readouts were retained together. Scar-matrix expression was summarized over the prespecified seven-member panel, with core-matrisome effects summarized separately. Gene-specific effects are listed alongside the panel means in Supplemental Table S10 and as gene-level blocks in Supplemental Table S7.

For the rabbit experiment, an additional model used joint filtering and TMM normalization across all 15 libraries with a five-condition design, followed by the same seven contrasts. This common model supplied original-scale comparisons across the culture conditions. The original pairwise effects and phenotype ranks remained the primary recorded readouts. Here coated denotes two-dimensional coated culture, and 50k and 100k denote the two cell densities of three-dimensional culture. ACTA2 was present in both contrasts with coated culture and absent after filtering in the original direct 100k-versus-50k contrast; the common model retained it. The direct comparison therefore has six original panel members and seven common-model members. The feline analysis retained its original pairwise models and separated the three day-5 drug-versus-drug contrasts from the day-5-versus-day-2 or day-0 contrasts [20].

Phenotype contributions were calculated as the prespecified weight multiplied by the gene's original within-contrast scaled rank, divided by the sum of available absolute weights. Positive- and negative-weight contributions used the same denominator and summed to the total score. For the fixed-coverage comparison, the contributing genes and weight denominator were restricted to the genes available in all three focal rabbit contrasts; ranks remained those of each complete original contrast. Existing HGNC families and core-matrisome membership were used to group the contributions. Family annotations retain their overlapping membership. Supplemental Table S10 reports the original and common-model expression effects, direct phenotype decompositions and all three same-day feline drug comparisons.

#### Extended control comparisons

Four public controls (human corneal lenticules) from PRJNA1533946 were aligned to the original GRCh38.p12 reference and counted with the original featureCounts settings. Prespecified-axis projection retained the original reference sample, TMM effective library sizes [21], control center, pseudocount and signed arms. The original ten-specimen scores were reproduced before the public controls were projected. Patient scores were summarized by the median of paired-eye scores, as in the original axis analysis. A separate analysis summed bilateral counts before projection; both estimators are provided in Supplemental Table S1. Patient-level exact tests of the prespecified-axis scores enumerated every assignment of the three patient labels (35 among the original seven profiles; 165 with the four public controls added). In each assignment the gene-wise control center was rebuilt from the control-labeled profiles of the original cohort, and the query gene ranks, the 300 signed genes per arm and both projections were recomputed; the public controls were projected onto that center and did not contribute to it. For the patient-median estimator, a patient profile entering a rebuilt center contributed the mean of its two eyes' normalized log expression, one weight per patient. Exact P was the number of assignments with a mean difference at least the observed difference divided by the number of assignments. In the comparison of the three patients with the four public controls the center remained the original four controls.

The panel extension summed bilateral counts before scoring, removed mitochondrial rows, aggregated counts by gene symbol and calculated $\log_2(\mathrm{CPM} + 1)$. Gene standardization and control centering were repeated separately for the original-control, public-control and combined-control comparisons. The seven scar, six SLRP and seven transparency members and their panel directions remained unchanged. The original cohort was analyzed with featureCounts and Salmon; the public-control extension used featureCounts. Exact tests enumerated the 35 assignments for three patients versus four donors and the 165 assignments for three patients versus eight controls. The mean patient-minus-control score was the test statistic, and BH adjustment covered the three panels within each comparison.

#### Patient-level phenotype tests

Patient profiles were formed by summing bilateral counts. Prespecified phenotype weights were applied to the complete within-profile gene ranks after centering expression on the reference controls. The common-gene consensus, meta-atlas and GSE199013 vectors each contained 12,526 genes; the full z-standardized source vectors contained 15,108 and 21,015 genes, respectively. All 57,132 gene rows entered ranking before the available prespecified weights were selected and normalized by their absolute-weight sum. The original featureCounts and Salmon comparisons each comprised three patients and four controls. The expanded featureCounts comparison added the four public controls.

Exact tests enumerated all 35 or 165 assignments of the three patient labels. For every assignment, the reference center was rebuilt from control-labelled units within the original seven-unit cohort, and the gene effects, complete gene ranks and phenotype scores were recomputed. The public controls contributed to the tested label allocations and group means and were projected onto the rebuilt center. The original and expanded enumerations used four and four to seven reference units per assignment, respectively. The weights remained fixed throughout enumeration. For the mean-difference test, the statistic was the mean patient score minus the mean of all control-labelled scores in that assignment; its one-sided lower-tail P value was the fraction of assignments at least as negative as observed, using the original $10^{-12}$ comparison tolerance. The observed allocation was included in the denominator.

Complete separation was evaluated separately with the binary statistic S = 1 when the highest patient score was below the lowest control score, and S = 0 otherwise. Its exact upper-tail P value was the fraction of assignments with S at least as large as the observed S. Thus, an observed separated allocation uses the number of separating assignments divided by all assignments; an observed nonseparated allocation has P = 1. Supplemental Table S2 reports the mean-difference P value, observed separation, number of separating assignments and separation-statistic P value in separate columns. It also retains the observed group means and the fixed-score tests. These statistics answer the mean-difference and complete-ordering questions separately.

#### Group-level phenotype estimation in human and public comparisons

For the human group-level estimate, bilateral counts were first summed within each patient and converted to $\log_2(\mathrm{CPM} + 1)$. The CPM denominator included all 57,132 nuclear-gene rows before gene selection. The gene-wise effect was the mean patient expression minus the mean control expression. Effects were ranked with average ranks for ties and scaled as $2r/(G + 1) - 1$. The ranked universe comprised genes with an available prespecified weight and a positive summed count across the design, giving G = 12,326 for three patients versus four donor controls and G = 12,348 for three patients versus eight controls or seven controls after summing counts from two public controls (libraries A1 and H1) before normalization. The phenotype score was the inner product of the consensus weights and scaled ranks divided by the sum of available absolute weights. The two source weightings were applied separately over the same common-gene universe. Public comparisons used the same group-effect ranking and weighting within their own mapped gene universes. Rabbit units were animal-level stromal pseudobulks, with two animals per arm at each freeze-injury time point and three per arm in alkali-burn fibrosis. The mouse allograft PBS-minus-isograft comparison used one pooled library of 4–6 corneas per condition. Human lower-tail exact tests recomputed the group effect, ranks and score for all 35, 165 or 120 label assignments while keeping the gene universe and weights fixed. A sensitivity analysis retained all 12,377 genes shared by the count matrix and weights, including genes with zero counts across the design.

#### Per-animal reconstruction of rabbit repair and fibrosis

Stromal pseudobulks were reconstructed from GSE328779 for two injured–contralateral pairs at each of days 7 and 28, and from GSE302936 for three fibrotic and three naive corneas. Reconstructed group effects reproduced the retained effects across 9,287 genes at day 7, 9,339 genes at day 28 and 11,285 genes in fibrosis, with maximal absolute differences of $5.55 \times 10^{-15}$, $5.11 \times 10^{-15}$ and $5.33 \times 10^{-15}$. Panel ranges were calculated using each injury/fibrosis library minus the comparator-arm mean and the injury/fibrosis-arm mean minus each comparator library. The two sets preserve the original biological arm and treatment direction. Group-level failure-signature similarity and genome-wide phenotype scores use their retained group contrasts. The panel ranges are listed in Supplemental Table S11. Direct time contrasts used joint filtering and TMM normalization of the eight GSE328779 stromal pseudobulks, with gene-level contrasts ranked and projected as for the single-time contrasts.

#### Linking public tissue outcomes to transcriptional comparisons

Published optical, structural and survival observations were linked by dataset, treatment arm, comparator and tissue, with RNA-collection and outcome-assessment times recorded separately. For GSE236626, RNA and graft appearance were assessed at day 7; OCT thickness was assessed at day 14 [22]. For GSE328779, stromal backscatter and the day-7/day-28 tissue states were taken from the freeze-injury study [23]; GSE302936 supplied day-14 opacity and lamellar structure in alkali-burn fibrosis [24]. These observations identify the tissue outcomes attached to the source comparisons.

GSE275249 anti-Gr-1 treatment arms were matched to the corresponding published tacrolimus experiment using shared investigators, transplant model, day-15 sampling and the three antibody-exposure windows [25]. Direct comparisons used anti-Gr-1 plus tacrolimus minus tacrolimus alone. Each paired count matrix was TMM-normalized and converted to logCPM with prior count 2; gene ranks and all projections were computed directly from that contrast. The prespecified five-module rejection score retained all 19 members. Scar, SLRP and transparency coverage was 7/7, 6/6 and 6/7. The consensus phenotype weights and both source weightings were applied separately. Early RNA exposure was days 0–7, delayed exposure days 8–15 and continuous exposure days 0–15. GSM9020665 was assigned to continuous and GSM9020666 to delayed exposure from the submitted sequencing-file and processed-matrix names, which also match the immune-cell counts reported for each arm; the GEO sample titles list these two windows in reverse. Published survival follow-up used early days 0–7, delayed days 8–42 and continuous days 0–42 dosing. Each RNA arm comprised one pooled library of 4–6 corneas; the source survival experiment retained its own animal units.

#### Cell-population-specific readouts during losartan-associated clearing

GSE291819 supplied day-3 nitrogen-mustard-injured corneal tissue including the limbus and part of the conjunctiva, treated with saline or losartan [26]. Raw counts were summed using the existing cluster assignments: clusters 0, 3 and 7 for myeloid cells and clusters 8, 12, 14 and 17 for the original stromal aggregate. All-cell, myeloid and stromal pseudobulks reproduced the existing count matrices. Each arm comprised one sequencing library. TMM-normalized $\log_2$ CPM effects, with prior count 2, were mapped by the original one-to-one ortholog procedure and projected using the prespecified programs.

Stromal clusters were then scored separately. Cluster 8 was Kera-enriched. Kera-positive cells and Kera-positive, Krt12-negative cells were also scored within this cluster using transcript detection to define the subsets. Consensus and source-specific phenotype weights were retained together. The meta-atlas weighting contrasts quiescent keratocytes with corneal fibroblasts, while the GSE199013 weighting contrasts corneal K_Fibro cells with fibroblasts from other ocular tissues. Source-specific scores use each source's eligible genes; consensus scores use the common genes and standardized weights. Cell fractions use all quality-controlled cells in the relevant library as the denominator.

Functional programs used Gene Ontology Biological Process sets with mouse annotation from org.Mm.eg.db and GO.db version 3.21.0. Gene effects were ranked within the native mouse pseudobulk contrast and summarized as mean scaled ranks. The retained sets cover inflammatory activity, epithelial junctions and migration, collagen organization, glycosaminoglycan metabolism and transport. The source study's day-3 haze and flow-cytometric measurements and day-14 structural observations retain their acquisition times. Source-linked phenotype and cell-count readouts are provided in Supplemental Table S11.

##### Decomposition of within-population state and captured-RNA composition

The anti-Gr-1 and losartan contrasts were decomposed using the original cell annotations, expression matrices, ortholog mappings and scoring functions. Anti-Gr-1 comparisons used T/NK, myeloid, stromal and other cells, with tacrolimus as the reference. The losartan stromal analysis retained clusters 8, 12, 14 and 17; clusters 8, 12 and 14 supplied the shared-cluster comparison. Captured cell fractions and captured RNA fractions were calculated separately from their corresponding cell counts and UMI totals. Each RNA condition retained its sequencing-library unit.

For populations present in both arms, four reconstructed pseudobulks combined reference or treated within-population expression with reference or treated captured-RNA shares. Each was scored against the same reference using the original normalization, complete gene ranks and prespecified weights. F00 denotes reference state and reference composition, F10 treated state and reference composition, F01 reference state and treated composition, and F11 treated state and treated composition. The state contribution was $[(F10 - F00) + (F11 - F01)]/2$ and the composition contribution $[(F01 - F00) + (F11 - F10)]/2$. For the phenotype score and rejection score, these contributions sum to $F11 - F00$. In the losartan analysis, the cluster-17 entry term was the original four-cluster score minus the score for the shared clusters, both relative to the same shared-cluster reference. Supplemental Table S5 retains the shared-cluster and full-aggregate results together.

The rejection-score analysis retained all 19 members and the complete 32,285-feature mouse rank universe. A1 retained its forward, reciprocal and symmetric projections at each corner as separate readouts. Consensus and both source weightings were retained for the phenotype score; common-gene and all-eligible source variants are labelled separately from the full z-standardized source variants used in Supplemental Table S2. Kera-positive and Kera-positive, Krt12-negative subsets retained the original raw-UMI detection definitions. The decompositions preserve the existing condition libraries and population assignments.

##### Corneal xenograft proteomics

For PXD031320, six corneal TMT channels were assigned to rejected and surviving pig-to-macaque grafts and donor genotypes using the source Methods section 2.4.2 and Table 1 [27]. IDPicker normalized reporter values were retained once per protein group; the public search result carries macaque-database proteins only. Macaque genes with a unique NCBI human ortholog were mapped to human genes, and multiple protein groups mapping to the same gene were summarized by median $\log_2$ intensity per channel. Each gene was standardized across the six channels using its sample standard deviation. Panel and module scores were the mean member z score; the phenotype score was the sum of prespecified consensus weights multiplied by member z scores divided by the sum of the available absolute weights. Comparisons used rejected minus surviving grafts across all donors and within WT donors, with independent two-sample t tests as in the source study (Supplemental Table S3).

##### Specimen allocation and clinical reference measurements

The six failed lenticules and four independent donor controls comprised the cohort for differential-expression, rejection-score, immune-repertoire and phenotype analyses. Two lenticules removed from an additional participant for another indication were sequenced separately as descriptive references outside group inference. Specimen assignments were checked against submission records and the sequencing provider's receipt dates. Original donor controls and public control libraries retained separate source labels. All three patients with failed lenticules were women; sex was not analyzed as a biological variable. The ethics approval covered protocol 1.0.

The preimplantation densitometry reference comprised Patient A and the additional reference participant. Repeat scans were averaged by eye, then by participant, with equal weight given to the two participants. Clinical intervals were calculated from dates established by operative signatures or corroborated entries from the first postoperative day. The first documented event leading to exchange was the primary-record entry prompting lenticule exchange; the interval before removal was the difference between those dates. Uncorrected visual acuity retained the recorded decimal scale.

##### Eligibility and extraction of published implantation reports

Eligible reports in humans covered intrastromal lenticules, surface and patch procedures, stromal rings or additions, historical procedures and xenogeneic tissue. Comparator arms involving penetrating or deep anterior lamellar keratoplasty were excluded, as were synthetic rings, biosynthetic corneas, and endothelial or limbal stem-cell grafts. Rejection affecting a subsequent penetrating graft was assigned to that graft. Reports were independently read twice and adjudicated by a third reader. Programmatic summaries used the adjudicated event category, author attribution, treatment sequence, procedure and specimen-examination fields. Reported eye and specimen counts were retained; missing quantities remained unreported.

##### Gene counting and clinical expression summaries

The sequencing provider generated the HISAT2 alignments [28] and featureCounts gene counts [17]. Recounting used unstranded paired-fragment assignments, required MAPQ ≥ 10 and both mates mapped, and excluded multimapping reads. The reproduced provider counts were used for differential expression. Alternative Salmon quantification used the same genome and annotation with sequence-bias and GC-bias correction [18].

To compare ACTA2 expression with the clinical ordering of patients, we averaged the two eye-level $\log_2$(CPM + 1) values in each patient. This summary was separate from differential-expression analysis of summed bilateral counts. Absence of INHBA counts in donor controls was checked against the alignments. The scar matrix and small leucine-rich proteoglycan panels retained positive directions. The transparency panel was reversed for directional testing and displayed in its biological direction. Panel membership and direction were identical for featureCounts and Salmon. The broader 13-gene transparency module used for matrisome-category calibration was scored separately from the seven-gene transparency panel.

##### Rejection-score calibration and resampling

The five modules comprised 33 human genes, of which 19 strict one-to-one orthologs were available in both mouse calibration datasets and both in-house quantifications. Gene-effect ranks, with mean ranks assigned to ties, were scaled within each genome-wide contrast and averaged within module. The module-mean vector was projected onto the GSE275249 PBS-treated allograft-minus-isograft vector and divided by the reference vector's squared length. This defined a rejection score of +1 for the reference contrast. GSE181893 PBS-minus-anti-CD80/86 provided the independent positive reference [29]. Leaving out one library at a time within each GSE181893 group produced the same-state reference contrasts.

Intervals for the positive-reference-minus-failed-lenticule difference were based on 2,000 within-group bootstrap resamples of reference libraries and patient or donor units. Individual-patient intervals used 10,000 gene resamples stratified by module. Patient-label tests and leave-one-patient-out projections used the full human modules. Gene-wise standardized expression was averaged to summarize immune-module activity across specimens.

##### Immune-repertoire inputs

TRUST4 reconstruction used native-depth FASTQ and aligned BAM inputs [19]. For common-depth comparisons, nine of the ten primary specimens were independently downsampled without replacement to 16,028,223 read pairs for 20 runs per specimen. The remaining specimen defined that depth and retained its native input. The same chain and CDR3 nucleotide sequence had to be recovered from BAM reconstruction as well as satisfy the sequence and resampling criteria.

#### Tissue-program projection and individual-library comparisons

The 20 members of the scar matrix, stromal small leucine-rich proteoglycan and transparency panels retained their panel-level failure directions in public comparisons. Gene-effect ranks were scaled as $2r/(G + 1) - 1$, where r was the average-tie rank and G the number of ranked genes. We averaged direction-aligned ranks across available one-to-one orthologs and recorded member coverage. Gene-set permutations drew 10,000 sets of the same size without replacement from all ranked genes, retaining each panel's direction.

Rabbit repair and fibrosis comparisons used stromal pseudobulks reconstructed per animal from GSE328779 and GSE302936 [23,24]. The day-7 and day-28 freeze-injury comparisons retained their distinct injured–contralateral pairs. The direct time-by-injury contrast compared changes in injured and contralateral tissue over the same period. Rabbit culture contrasts used pairwise edgeR quasi-likelihood $\log_2$ fold changes from GSE337743 [30]. Each comparison, including the direct 100,000-versus-50,000-cell contrast, was ranked separately. Phenotype contributions were calculated from each gene's external weight and contrast rank using a common absolute-weight denominator for positive and negative weights.

For individual-library projections, each treatment library was compared with the comparator-arm mean, and the treatment-arm mean was compared with each comparator library. Both sets retained the treatment-minus-comparator orientation and were summarized separately from group-model estimates. This procedure was also applied to reconstructed rabbit tissue panels. The 69 intervention and induction comparisons retained their species, sampling time, comparator and replicate unit. GSE275249 comprised one pooled library of 4–6 corneas per condition [25]. GSE236626 included three corneas per arm [22]. GSE337743 included three culture libraries per condition [30], and GSE231796 included five culture replicates per group [20]. Day-5 feline drug-treated cultures were compared separately with the day-2 induced state and day-0 baseline. Direct day-5 comparisons used troglitazone, troglitazone plus GW9662 and UK-5099. Complete 19-member rejection scores and scores using available members were recorded separately.

INHBA, FST and FSTL3 retained the gene-effect scale of each dataset. TGF-β induction was compared with untreated human cells, serum-free rabbit cultures and day-0 feline cultures [31,32,20]. Human gene effects were differences in $\log_2$(author-normalized count + 1). Rabbit and feline group effects were edgeR $\log_2$ fold changes, and individual-library effects were differences in $\log_2$CPM. The serum response factor/myocardin-related transcription factor (SRF/MRTF) program was summarized from the mean ranks of its 20 target genes. Failure-signature and external-axis projections used continuous within-contrast ranks. The expanded-control projection retained the original reference, control center and signed arms. Patient values for these projections were the medians of paired-eye values; projections from summed bilateral counts were calculated separately. In label tests, the control center, query ranks, signed-arm membership and bidirectional projections were recomputed for every assignment.

#### Linkage of tissue outcomes and the ABT-263 experiment

Published optical, structural and graft-survival measurements were linked to the expression comparisons by treatment, comparator and tissue. RNA-collection and outcome-assessment times were recorded separately. In the eight independent in vivo experiments, reported outcomes were classified as better or worse than the comparator. A positive phenotype score matched a better outcome, and a negative score matched a worse outcome.

PRJDB43191/CRA014662 provided four whole-cornea libraries each from unoperated eyes, vehicle-treated allografts and ABT-263-treated allografts, sampled on postoperative days 10–12 [33]. Reads were quantified with Salmon against the mouse transcriptome [18]. All twelve libraries were TMM-normalized together with prior count 2 before the phenotype, failure-signature and scar matrix definitions were applied. The matched outcomes were the source study's slit-lamp observations and Kaplan–Meier graft survival for ABT-263 versus vehicle, with 14 grafts per group. Unoperated corneas supplied the common reference for library-level scores.

#### Anti-Gr-1 comparisons within annotated cell types

We analyzed the early, continuous and delayed anti-Gr-1-plus-tacrolimus conditions against tacrolimus alone in GSE275249 [25]. RNA was collected on day 15 after antibody exposure on days 0–7, 0–15 and 8–15, respectively. The corresponding survival experiments used days 0–7, 0–42 and 8–42. Each RNA condition comprised one pooled library from 4–6 corneas. GSM9020664, GSM9020665 and GSM9020666 represented early, continuous and delayed exposure, respectively; GSM8473611 was the tacrolimus reference. Continuous and delayed assignments followed the submitted sequencing-file and processed-matrix names. Each paired count matrix was TMM-normalized and converted to $\log_2$CPM with prior count 2 before direct comparison.

Counts were aggregated by the final cell-type annotations. We retained all annotated types and recorded cell counts and captured unique molecular identifiers (UMIs) separately. Within-type expression was compared only for types observed in both treatment and reference libraries. Types observed in only one arm remained in the full-tissue comparison, with their within-type contrasts left undefined. Counts were checked by summing final types back to the four broad populations and comparing gene totals and cell numbers with the original aggregates. The complete 32,285-feature mouse universe, one-to-one ortholog mapping and phenotype weights were retained.

For types observed in both arms, four reconstructed profiles combined reference or treated within-type expression with reference or treated captured RNA shares. F00 combined reference expression and composition. F10 replaced expression only, F01 replaced composition only and F11 replaced both. All four profiles were normalized, ranked and scored against the same shared-type reference. The state contribution was $[(F10 - F00) + (F11 - F01)]/2$, and the composition contribution was $[(F01 - F00) + (F11 - F10)]/2$. The support-set residual was the full observed score minus F11. This residual retained the effect of types outside the shared set and the difference between full and shared reference profiles. State, composition and the support-set residual were reported separately.

We repeated the calculation with consensus weights and both source weightings, retaining common-gene and all-eligible-gene variants. Direct phenotype comparisons within each type used that type's expression contrast. These scores were reported alongside captured RNA shares and cell counts. Whole-tissue contributions were calculated from reconstructed expression profiles. The 19-member rejection score was decomposed in the same way. Forward, reciprocal and symmetric A1 projections were reported for each reconstructed profile. An undefined F00 projection remained undefined.

#### Aire-deficient corneas after rewetting

GSE272700 provided single-nucleus RNA sequencing from wild-type mice, untreated Aire-deficient mice and Aire-deficient mice after 14 days of topical PBS rewetting, with one pooled library per condition [34]. We compared rewetting with untreated deficiency, and untreated deficiency with wild type. The source study's measurements of collagen arrangement, fibril spacing and stromal thickness were linked to these expression contrasts.

The primary analysis retained nuclei with at least 200 detected genes, at least 500 UMIs, no more than 6,500 detected genes and a mitochondrial fraction below 5%. Libraries were clustered separately with Seurat v5.3.0 [35]. Lineages were assigned by cluster-level enrichment of multiple marker genes. Module-score annotation and annotation accounting for ambient keratin signals were evaluated separately. Five broad lineages were retained: stromal/keratocyte, epithelial, immune, endothelial/vascular and other. Stromal nuclei were additionally integrated with Harmony v1.2.3 [36] and reclustered. Counts were summed within library and lineage or stromal subcluster.

Pseudobulk comparisons used TMM normalization, genes measured in both libraries, one-to-one ortholog mapping and the phenotype rank-projection procedure. When multiple entries mapped to a human gene, the entry with the largest absolute effect was retained. Gene rows absent from the rewetting library were excluded from the shared measured-gene set. RNA fractions calculated from all measured UMIs were recorded separately from fractions used for decomposition over the shared measured genes. Consensus, source-specific z-standardized and source-specific eligible-gene weights were all applied.

To equalize library depth, nonzero wild-type and untreated-deficient counts were independently thinned by binomial sampling, with retention probabilities of 0.590087694 and 0.476092436. Rewetting-library counts were unchanged. The target was the rewetting-library mean of 1,568.549845 UMIs per nucleus in the primary analysis. Thinning was applied to all prepared nuclei before repeating the same quality filters, separate-library clustering and lineage annotation. Stromal nuclei were again integrated and reclustered. Thinning and quality filtering were repeated with a second seed, retaining the first seed's stromal annotations. The two seeds were 272701 and 272702.

We also scored stromal pseudobulks separately within the UMI intervals [500,1000), [1000,2000), [2000,4000), [4000,8000) and [8000,Inf). Each interval was independently normalized and scored. Additional quality-filter analyses used a mitochondrial fraction below 10% or a minimum of 500 detected genes. Each analysis retained its nucleus counts and annotation results.

State and composition were separated at the five-lineage and stromal-subcluster levels by the four-profile procedure above. Shared populations were observed in both compared libraries. The full-minus-F11 residual was retained separately from state and composition. Nucleus counts, captured UMIs and lineage assignments were reconciled with the pseudobulk totals. Each condition retained its pooled sequencing library as the analysis unit.

##### Statistical estimators

For patient-level panel comparisons, the statistic was the patient-minus-control mean difference. The three panel P values were adjusted together within each quantification and control set. Phenotype mean-difference tests used the lower tail. Complete separation was evaluated with a binary statistic equal to one when the highest patient score was below the lowest control score and zero otherwise. Its upper-tail P value used the same complete set of label assignments. Mean differences and complete-separation statistics were recorded separately. Gene-set enrichment and expression-matched random-set analyses retained their gene-level units. Human group-level phenotype estimates and public-study estimates used the same within-comparison gene-effect ranking and weighted score, retaining each experiment's biological or pooled-library units.

#### Supplemental results

##### External implant-remodeling scores and the scar component

We compared the failed lenticules along externally defined transplant-response, repair and implant-remodeling axes. The genome-wide transplant-response axis, A1, was derived from the full costimulation-blockade contrast; it therefore measures a broader transcriptional response than the five-module rejection score. The implant-remodeling axis, A3, contrasted a fibrosis-associated synthetic implant with an extracellular-matrix scaffold. A3 placed every patient above every control, with a pooled failed-tissue score of +8.66 (Figure S6B). A1 showed overlap between the groups: failed tissue scored +4.85, and donor control 2 scored +6.18. Donor control 2 was also the control with the reproducible TRA clonotype. All controls were retained. The murine allograft-versus-isograft reference occupied a different position, with high A1 and low A3.

A3 retained patient–control separation after composition adjustment across two human corneal references and two deconvolution methods (Figure S6C) and after removal of variation shared with the other axes (Figure S6, B and E). A1 remained non-separating, and the repair axis A2 lost separation after mutual axis residualization (Figure S4C). The scar-matrix panel explained most A3 variance ($R^2 = 0.93$), and removing the scar-matrix component removed A3 separation (Figure S6A). Thus, A3 expressed scar burden as an externally derived remodeling score. The tissue-state interpretation comes from this scar response together with the independently measured loss of keratocyte phenotype.

##### One-eye-per-patient convergence enumeration

All eight one-eye-per-patient selections reached the 0.0286 patient-level floor before technical adjustment and remained directionally positive after it. All four selections containing one Patient B specimen were among the five that fell short of $P \leq 0.05$ after adjustment, and that specimen's adjusted margin was −0.022 (Figure S1C).

##### Corneal densitometry after uncomplicated implantation

Increased anterior and mid-stromal density has also been reported after uncomplicated allogeneic lenticule implantation [37]. That study included 14 allogeneic recipient eyes from 10 patients and 8 autologous recipient eyes from 8 patients, with no postoperative haze or infection reported (Supplemental Table S12, report L063).

##### Specular microscopy and anterior segment photography

Specular microscopy recorded endothelial cell densities of 2,339 to 3,296 cells/mm² at failure identification or exchange; morphology indices were outside the reference range in all three patients, so density rather than morphology is described as preserved. In the 41 anterior segment photographs that were reviewed for rejection signs (of 107 available), neither corneal neovascularization nor ciliary injection was seen.

##### Deconvolution reliability diagnostics

Sixteen of the twenty primary fits (ten specimens × two references) had negative reconstruction $R^2$ (median −0.035; Figure S2B). Whole-composition dispersion between pipelines was substantial (mean pairwise Jensen–Shannon distance 0.348 to 0.506, maximum 0.716) and was not confined to low-complexity libraries (Supplemental Data 1, Composition_diagnostics). The stromal proportion moved in opposite directions under the two methods (Figure S2D). Activated and resting keratocyte signatures were nearly identical in both references (correlation 0.98 and 0.97; Figure S2C), and the myofibroblast-like candidate correlated at 1.00 and 0.98 with other stromal signatures, as expected when the references are normal corneas that contain no true myofibroblasts.

The consensus phenotype score (12,526 external weights) applied to the same deconvolution outputs was lower in failed tissue than in donor controls under all four reference-method pipelines (observed difference −0.183 with GSE186433 and −0.179 with GSE199013; three patient units versus four donors). The composition prediction accounted for −0.057 (BayesPrism) and −0.138 (MuSiC) with GSE186433 and for −0.041 and −0.074 with GSE199013, which is 23% to 75% of the observed difference, and the model residual for the remainder (−0.126, −0.045, −0.138 and −0.105). BayesPrism lineage expression was lower in failed tissue in the stromal (−0.166 and

−0.174), myeloid (−0.154 and −0.155) and T/NK (−0.094 and −0.135) compartments for the two references; MuSiC provides no lineage expression estimate (Supplemental Table S5J).

##### Spatial co-occurrence of myeloid and stromal programs in external human corneas

In an external Visium data set of thirteen human corneas (eight keratoconus, five controls) [15], spots in the top 20% of the myeloid program within a cornea had more stromal-high neighbors than the within-cornea permutation expectation for the activated-stroma, myofibroblast-ECM and matrisome programs in all thirteen corneas (equal-cornea mean excess +0.103, +0.070 and +0.111), and dual-high spots were more frequent than expected in all thirteen (Figure S5A). The continuous cross-Moran statistic exceeded the null in every cornea for each of the three stromal programs (mean excess +0.163, +0.118 and +0.181; Figure S5B). Myeloid and stromal-matrix programs therefore share spatial neighborhoods in human corneal tissue.

##### Hallmark allograft-rejection enrichment and leading-edge annotation

The leading-edge analysis used for Figure 4C gave a positive Hallmark allograft-rejection enrichment score (NES = 1.45663918, nominal P = 0.0090955247, Hallmark-family adjusted P = 0.0324840168). The Figure 4C annotation lists the 45 leading-edge genes together with their patient-level edgeR $\log_2$ fold changes. INHBA ranked first, and LCP2 was the sole member assigned to the T-cell/TCR category. The other categories covered myeloid and innate activity, cytokine signalling, antigen presentation, leukocyte adhesion, stromal repair and metabolism.

The separate pathway-overview run gave NES = 1.45532755, nominal P = 0.0056656474 and Hallmark-family adjusted P = 0.0140150248. The immune-panel analysis gave NES = 1.45503 and adjusted P = 0.0732133 after joint BH adjustment across the 13 eligible immune and polarization sets. These values belong to their respective runs and multiplicity families; the Figure 4C annotation and its enrichment statistics are taken together from the leading-edge run.

##### TCR candidate attrition in the failed lenticules

Full-depth reconstruction returned 7 to 68 candidate clonotypes per failed lenticule (133 in total), 55 of which satisfied criteria (i) to (iv) before depth matching; in the two Patient C specimens all 33 candidates were removed at that stage (Figure S13A). One of the 55 was a T-cell receptor, the TRB clonotype from Patient B recovered in 10 of the 20 depth-matched resamples (Figure 4D and Figure S13A). A T-cell receptor sequence was therefore detectable in the failed tissue but did not meet the reproducibility criterion. Recovered receptor counts depend on read depth, which is why specimens were compared at a common depth: in one original unimplanted donor control, productive BCR clonotypes rose from a median of 341 at 5 million read pairs to 464 at 23 million, and reliable BCR clonotypes from 217 to 291, whereas the single TCR clonotype was productive at every depth and reliable from 10 million read pairs (Figure S13B).

##### BCR clonotype structure

Reliable BCR clonotype counts were 122, 263, 32 and 48 in the four controls (median 85, range 32 to 263) against 2, 4, 11, 7, 0 and 0 in the six failed lenticules (median 3, range 0 to 11). Every control exceeded every failed specimen, and this held when the specimen defining the common-depth floor was excluded. A lower reliable BCR count in the failed lenticules is as consistent with replacement of donor-derived cells as with a weaker humoral response. V-gene usage was almost non-repeating: nine distinct V genes among eleven clonotypes in one eye, seven among seven in another. Per-clonotype support was low (median 8 to 16 fragments), and no clonotype was shared between the two eyes of a patient or between patients. The clonotypes carried somatic hypermutation (median V-region identity 91.7% to 95.1%, minimum 83.0%), indicating antigen-experienced rather than naive B cells. With two to eleven clonotypes per eye, the absence of sharing is bounded by sampling and is not by itself strong evidence.

##### Glucocorticoid-receptor fingerprint, gene-level values

The median difference across the ten genes was −0.41 $\log_2$ units, with FKBP5 (+0.47) and PER1 (+0.65) marginally higher and KLF15 (−1.04), ANGPTL4 (−1.12), ZBTB16 (−0.82) and TSC22D3 (−0.44) lower. No gene reached significance, and the receptor gene itself was unchanged (NR3C1 −0.25).

Corticosteroid target engagement separated from the phenotype response in the public comparisons. In injured rat cornea, dexamethasone versus IgG at 48 h [38,39] increased FKBP5 and TSC22D3 by +1.5379 and +1.9394 on the source log-expression scale. The median difference across nine available glucocorticoid-receptor target genes was +0.0790, with ZBTB16 unavailable. Failure-signature similarity increased by +5.8795 and the phenotype score by +0.0222. In normal mouse cornea 6 h after dexamethasone [40], all ten target genes increased, with a median $\log_2$-expression difference of +0.6977, while the phenotype score decreased by −0.0495 (Supplemental Table S7).

#### Pathogen screen

Against an empirically determined stability floor of 5,158 non-host read pairs, each of the eight taxa passing the effect-size and read-count thresholds drew at least half of its failed-group reads from a single specimen (Figure S10, A and B). No between-group test is reported, because the enumeration available was at specimen rather than patient level. At this sequencing depth and with this database, no reproducible non-host signal accompanied the failure state.

#### Matrisome opposition split against two allorejection resources

Core-matrisome directions relative to the failed lenticules differed between the two allorejection resources. Among core-matrisome genes with usable effects, opposite changes accounted for 64.5% (109 of 169) in the anchoring rejection reference and 43.4% (76 of 175) in the second, independent allorejection resource, where 56.6% (99 of 175) were concordant (Supplemental Table S1).

#### Remodeling signals in the human failed lenticules

INHBA increased from alignment-verified zero counts in all four original unimplanted donor controls to 517–10,800 counts in the failed lenticules ($\log_2$ fold change +15.14; FDR = $4.25 \times 10^{-4}$). ACTA2 separated the groups and followed the B < A < C patient ordering of exchange-day reflectivity. SMAD-target transcripts increased without increased SMAD-factor expression. WWTR1/TAZ and its targets CCN2 and CCN1 increased. Collagen-category expression increased, and epithelial–mesenchymal transition was the top-ranked Hallmark pathway. The core-matrix-expression-minus-phenotype standardized difference was 4.02 in failed tissue and 3.48 in rabbit alkali-burn fibrosis. These findings connect the fibrotic tissue state with the inducible remodeling programs examined in Figure 8.

#### External reference context of the failed human tissue

Among the failed-tissue contrast and 18 external reference states, the failed lenticules had the highest core matrix expression score and the lowest externally weighted phenotype score ($z = +1.70$ and $-2.32$, respectively; Supplemental Table S1R; Figure S11A, which plots 17 of these states and shows GSE181893 by its PBS arm). The tested human TGF-β1-induced corneal myofibroblast state retained a positive phenotype score ($z = +1.06$). Both allorejection references had lower phenotype scores while differing in the direction of core matrix expression. The phenotype displacement persisted after deletion of individual HGNC families (Supplemental Data 1, Phenotype_family_omissions) and the leading one to three contributing genes (Supplemental Data 1, Phenotype_top_deletions). The high-information score from 3,098 genes in the top quartile of absolute external weights lay outside the distribution from 1,000 expression-matched random sets (two-sided $P = 0.001998$; Supplemental Data 1, Phenotype_null_summary; Supplemental Data 1, Phenotype_matched_sets). This reference comparison and matched-set test describe the tissue-state contrast; patient-level phenotype tests are reported separately in Supplemental Table S2.

#### Remodeling programs during scar-free repair and fibrosis

The freeze-injury series linked recovery of transparency to changing stromal programs within one injury model [23]. Haze peaked at day 7 and was lower at day 28, approaching normal inherent backscatter. In the corresponding injury-minus-contralateral contrasts, the phenotype score was −0.0414 at day 7 and −0.0146 at day 28, whereas failure-signature similarity was +3.621 and +12.993. Core matrix expression scores were +0.0332 and +0.1046. Over the same time points, INHBA changed from +1.332 to −0.319 on its source effect scale, and the SRF/MRTF target readout changed from +0.042 to −0.188. Day-14 alkali-burn fibrosis combined a phenotype score of −0.1405 with

INHBA +6.182 and SRF/MRTF +0.726 [24]. Each time point retains its own injury-minus-control contrast and original biological units. These comparisons distinguish sustained matrix expression from the remodeling signals accompanying the clearing of haze (Figure 5A; Figure 8A and Figure S11B; Supplemental Table S6). Day-28-minus-day-7 contrasts were computed directly after joint normalization of all eight libraries. Relative to the time-matched contralateral eyes, the consensus phenotype score increased by +0.040 and failure-signature similarity by +2.46, and the scar-matrix panel declined by −0.535. In injured eyes alone, the consensus phenotype score increased by +0.047 from day 7 to day 28. The meta-atlas weighting, which measures keratocyte quiescence relative to corneal fibroblasts, increased by +0.081 in the time-by-injury contrast. The GSE199013 weighting, which measures corneal specialization relative to fibroblasts from other ocular tissues, decreased by −0.018. The two day-28 rabbits had injured-minus-contralateral consensus phenotype scores of +0.011 and −0.047 (Supplemental Table S6).

##### Contributions to the culture-density difference in keratocyte phenotype

The rabbit culture experiment separated scar suppression from the direction of the phenotype response. Relative to coated 2D culture, the 50k and 100k 3D conditions both reduced the prespecified scar program, with similar positions in their gene-effect rankings (−0.9643 and −0.9646). On the original expression scale, the seven-member mean $\log_2$ fold changes were −3.2461 and −2.8436, and the core-matrisome means were −1.0409 and −0.5530. Thus, the 100k condition suppressed these matrix transcripts less strongly. The original direct 100k-versus-50k contrast had a scar-panel mean $\log_2$ fold change of +0.5576 across six available members; the common model retained all seven members and gave +0.4121 (Supplemental Table S10, A and B).

The direct 100k-versus-50k comparison had a lower phenotype score (−0.1461), comprising −0.1310 from genes with negative external phenotype weights and −0.0150 from genes with positive weights. The two source weightings retained the same direction and the predominance of negative-weight contributions. Fixing coverage to genes shared by the three focal contrasts gave contributions of −0.1287 and −0.0189, respectively. These contributions retained the original complete gene ranks and a shared absolute-weight denominator (Figure S7A; Supplemental Table S10C).

The negative-weight contribution included extracellular-matrix, cell-adhesion, surface-recognition and receptor-ligand families associated with the comparison fibroblast states. Core-matrisome members contributed −0.0184 through negative weights and +0.0065 through positive weights; the other annotated families are retained in Supplemental Table S10D. KERA itself increased in the direct comparison ($\log_2$ fold change +0.8934). The seven-gene transparency panel moved away from the failure direction at both 3D densities, whereas the genome-wide phenotype score increased at 50k and decreased at 100k. The density response therefore extended beyond the small transparency panel.

##### Induction and post-induction remodeling programs in corneal cells

INHBA and SRF/MRTF target ranks increased during TGF-β induction in human, rabbit and feline corneal systems. In feline fibroblasts [20], day-5 troglitazone and UK-5099 cultures had lower failure-signature similarity than the day-2 induced state (−6.907 and −6.471), together with higher phenotype scores (+0.0132 and +0.0217). Against day 0, phenotype scores remained lower (−0.0114 and −0.0182, respectively). INHBA and SRF/MRTF targets also decreased in both day-5-versus-day-2 comparisons. Day 2 and day 0 are the induction and pre-induction references for these time-separated comparisons (Figure 8, B and C; Supplemental Table S7).

The three same-day drug comparisons distinguished scar-matrix expression from the phenotype response within the day-5 cultures (Figure S14, A and B; Supplemental Table S10E). Troglitazone versus UK-5099 and troglitazone plus GW9662 versus UK-5099 both had lower scar-matrix expression (mean $\log_2$ fold changes −4.2351 and −6.2684). Their meta-atlas phenotype scores were lower (−0.0074 and −0.0074), whereas their GSE199013 scores were higher (+0.0162 and +0.0202), giving consensus scores of +0.0046 and +0.0068. Troglitazone versus troglitazone plus GW9662 had higher scar-matrix expression (+1.9958) and lower phenotype scores under both source weightings (−0.0072 and −0.0196), with a consensus score of −0.0156. INHBA increased in the troglitazone-versus-UK-5099 comparison (+0.6827), while the SRF/MRTF readout decreased (−0.6794). The original study used GW9662 to examine PPARγ dependence and interpreted the troglitazone response through a PPARγ-independent mechanism reproduced by mitochondrial pyruvate carrier inhibition with UK-5099 [20]. The shared day-5 groups each comprised five culture libraries.

#### Treatment courses in the published implantation record

The adjudicated set contained 285 report records, 284 after removal of one confirmed duplicate, and 4,938 reported tissue-implanted eyes. Opacity-family events were recorded in 95 records with 171 known affected eyes; an attributed cause was supplied for 53 eyes in 30 records. Rejection or suspected rejection was an author-assigned label for five eyes in two reports. The records covered intrastromal lenticules, stromal rings, surface and patch procedures, historical procedures and xenogeneic implants (Supplemental Tables S12 and S13).

Among 12 corticosteroid-exposed opacity-affected eyes, four resolved after corticosteroids alone and two after corticosteroids plus a procedure. A further resolving event was spontaneous. All seven resolving events were haze or diffuse lamellar keratitis within the first postoperative month. Three implanted-tissue whitening, fibrosis or melting events were unchanged or had mixed responses after corticosteroid exposure; the other two treated events involved an interface abscess or epithelial ingrowth. No report described corticosteroid reversal of established opacity within the implanted tissue. Across 80 report records, 346 specimens obtained by explantation or research biopsy were reported. Examinations included microscopy and other assays; no gene-expression profiling was reported. The molecular-level assays were HLA typing of cells cultured from removed tissue and matrix-metalloproteinase extraction from biopsy material. Modality and treatment counts are retained separately in Supplemental Table S13.

#### Extended controls and the scar component

The original-cohort scar, SLRP and transparency panels each reached exact P = 1/35 in both quantifications. With all eight controls, scar and transparency each retained complete patient–control separation (P = 1/165; BH FDR = 0.009091), while SLRP scores overlapped between patients and controls. The expanded-control SLRP mean difference was +0.4285, the separation margin was −0.9231 and exact P was 25/165 (Supplemental Table S1). For the prespecified implant-remodeling axis, all three patient medians and all six individual failed lenticules exceeded all eight controls (Figure S6D). The patient-median separation margin was +4.7206 (P = 1/165). Summing bilateral counts before projection gave a margin of +4.9039 with the same exact P. Both estimators are retained in Supplemental Table S1.

The consensus phenotype score placed every patient below every original control in both quantifications and below every control in the expanded featureCounts comparison (Figure 3D; Supplemental Table S2). In the original cohort, the mean-difference and complete-separation statistics each gave P = 1/35. In the expanded comparison, the mean-difference P was 2/165 = 0.01212121 and the complete-separation P was 1/165 = 0.00606061. The two separate common-gene source weightings each gave a mean-difference P of 2/165 in the expanded comparison; the GSE199013 weighting also retained complete separation. All source variants and the original unimplanted donor control and public control means are listed separately in Supplemental Table S2.

#### Public outcomes and phenotype readouts

Figure S9 places the six focal comparisons within all 69 intervention and induction comparisons retained for analysis, one point per dataset and contrast, with failure-signature similarity (Figure S9A) and core matrix expression (Figure S9B) each plotted against the phenotype score in the original treatment- or induction-minus-comparator direction. Early and continuous anti-Gr-1 treatment abolished the graft-survival benefit associated with tacrolimus in the source experiment, whereas delayed treatment did not [25]. Direct day-15 RNA comparisons with tacrolimus lowered the rejection score after early, continuous and delayed exposure (−0.2216, −0.3285 and −0.9509). The broad transplant-response axis A1 increased after early and continuous exposure and decreased after delayed exposure (+15.3241, +12.0855 and −3.1424). The consensus phenotype score decreased in the early and continuous arms and increased in the delayed arm (−0.0122, −0.0681 and +0.0148); both source weightings decreased after continuous exposure and diverged in the early and delayed arms. Failure-signature similarity decreased after early and delayed exposure and increased after continuous exposure, and the transparency panel shifted toward the patient failure direction in all three arms (Supplemental Table S11).

#### Within-population state and captured-RNA composition

Decomposition across T/NK, myeloid, stromal and other cells assigned the lower rejection score to a negative within-lineage state contribution in every arm (−0.753323, −0.894412 and −0.554325 after early, continuous and delayed exposure). The composition contribution opposed the decline after early and continuous exposure and reinforced it after delayed exposure. In contrast, the composition-only A1 projections reproduced its time-window pattern (+23.5428, +26.9167 and −15.7732). The consensus phenotype score combined a negative state contribution in every arm (−0.098503, −0.096690 and −0.096761) with a positive composition contribution (+0.086264, +0.028631 and +0.111558). The positive delayed-arm aggregate therefore arose from the change in captured-RNA composition. Each of the four lineages separately had a lower consensus phenotype score than the corresponding tacrolimus reference (Figure S15A; Supplemental Table S5, C–F). At final-cell-type resolution, the delayed-arm increase combined a common-support state contribution of −0.0911, a composition contribution of +0.1046 and a support-set residual of +0.0014. The keratocyte/stromal fibroblast population contained 529 cells and had a direct phenotype score of −0.0608 relative to tacrolimus alone (Figure S15B; Supplemental Table S5I). These results separate the within-lineage rejection response, the composition-sensitive broad transplant response and the aggregate phenotype response.

#### Cluster-specific phenotype responses during losartan-associated clearing

In the losartan study, day-3 haze was 32% lower and independent flow cytometry showed lower myeloid-cell frequency [26]. The consensus phenotype score decreased in the original four-cluster stromal aggregate (−0.0641) and increased in the Kera-enriched cluster 8 (+0.0169) and its Kera-positive, Krt12-negative subset (+0.0083). Clusters 12 and 14 had lower phenotype scores (−0.0443 and −0.0425). Both source weightings reproduced the aggregate decrease and Kera-enriched increase (Figure S8A; Supplemental Table S11). The quality-controlled myeloid fraction fell from 65.94% to 37.63% (Supplemental Table S5).

Relative to the shared-cluster reference, the original aggregate phenotype-score change comprised a within-cluster state contribution of −0.035586, a common-cluster composition contribution of +0.016670 and a cluster-17 entry contribution of −0.045140 (Figure S8C). The common-cluster composition change offset part of the decline, while within-cluster state and the added cluster lowered the aggregate. Cluster 17 represented 19.09% of stromal cells and 41.98% of captured stromal RNA after losartan. The Kera-enriched cluster's captured-RNA share fell from 55.05% to 5.89%, and its cell share from 80.10% to 32.90%. Supplemental Table S5, A and B, retains both the shared-cluster and full-aggregate results and separates cell from RNA representation. Acute clearing accompanied a population-specific phenotype response and altered tissue composition.

#### Rapamycin and resveratrol readouts

In the mouse allograft experiment, rapamycin versus PBS lowered the rejection score (−1.007) and the A3 implant-remodeling axis score (−7.766), while increasing failure-signature similarity (+4.349) and lowering the phenotype score (−0.1131). Each condition was one pooled RNA library from 4–6 corneas (Supplemental Table S7). In the rat allograft experiment, resveratrol versus saline lowered the available-member rejection score (−0.445). This estimate used 18 of the 19 prespecified genes, with JCHAIN unavailable, and retained three corneas per arm (Supplemental Table S7).

#### Phenotype scores relative to unoperated corneas after ABT-263

In the ABT-263 allograft experiment, library-level consensus phenotype scores relative to a common unoperated-cornea reference were lower for both ABT-263-treated and vehicle-treated grafts. The respective differences of group means were −0.209 and −0.210, with four libraries per group at postoperative days 10–12. The difference between the two allograft groups on this common reference was +0.0017 (Supplemental Table S4, note E).

#### Human and public phenotype differences under the same group-level estimator

Under the same group-level estimator, failed human lenticules had negative phenotype scores with all three weightings. Values are given in consensus, meta-atlas and GSE199013 order. Against four donor controls, the scores were −0.1843, −0.1450 and −0.1679. Against all eight controls, they were −0.2424, −0.1900 and −0.2215. With the two public-control libraries A1 and H1 merged, the three-patient-versus-seven-control scores were −0.2319, −0.1796 and −0.2143. Rabbit freeze-injury scores were −0.0414, −0.0681 and +0.0018 at day 7 and −0.0146, +0.0195 and −0.0472 at day 28. The direct day-28-minus-day-7 time-by-injury effects were +0.0399, +0.0806 and −0.0181. The corresponding injured-eye time effects were +0.0466, +0.0519 and +0.0254, and contralateral-eye time effects were +0.0137, −0.0296 and +0.0567. Rabbit alkali-burn fibrosis versus naive stroma gave −0.1405, −0.1334 and −0.1035. Mouse allograft PBS versus isograft gave −0.1176, −0.0738 and −0.1258. The human consensus scores were therefore lower than those in rabbit fibrosis and the clearing day-28 repair state (Figure 5C; public reference values in Supplemental Tables S4 and S6). The consensus group-level lower-tail exact P values were 5/35, 5/165 and 5/120 for the four-, eight- and seven-control comparisons, respectively. These test the group-level rank statistic and are separate from the unit-level complete-separation statistic reported in the main Results ($P = 0.0061$). Retaining all 12,377 genes shared by the count matrix and weights gave consensus scores of −0.1823, −0.2413 and −0.2308 for the same three human comparisons.

### Supplemental tables

**Supplemental Table S1. Prespecified tissue programs, patient-level comparisons and rejection calibration.** Panel and module membership, original-cohort featureCounts and Salmon scores, public-control extension comparisons, exact-label tests, BH adjustments, reference scores and bootstrap intervals retain their analytic units. The expanded featureCounts comparison uses four original unimplanted donor controls and four public controls. Prespecified A1–A3 axis comparisons retain paired-eye score medians and summed-bilateral-count projections as separate estimators. The final block lists the gene-level direction decomposition against the two allorejection references, including core-matrisome membership.

**Supplemental Table S2. Complete-estimator phenotype tests and observed group means.** Bilateral specimens are combined within each patient. The original cohort uses four original unimplanted donor controls; the expanded featureCounts comparison adds four public controls. Prespecified source weights are applied separately. Common-gene vectors contain 12,526 genes; the meta-atlas and GSE199013 full z-standardized vectors contain 15,108 and 21,015 genes. All exact tests include the observed allocation. Mean-difference P values use the lower tail of the complete estimator, rebuilding the center and all gene ranks within each allocation. The binary complete-separation statistic is S = 1 if every patient score is below every control score and S = 0 otherwise. Its upper-tail P is the number of separating allocations divided by all allocations when the observed S = 1, and is 1 when the observed S = 0. The count of separating allocations is retained separately. Fixed-score P values come from the permutation of unchanged scores. Values are reported at the precision of the analysis.

**Supplemental Table S3. Tissue and immune-program member scores in rejected and surviving corneal xenografts.** The public PXD031320 search result carries macaque-database proteins only. Scores summarize protein members mapped to human orthologs in six corneal grafts, four rejected (graft score 6; two GTKO and two WT donors) and two surviving (graft score 0; both WT donors). Differences are rejected minus surviving means for all grafts and the WT subset. P values are two-sided independent two-sample t tests with equal variance, as in the source study. Transparency scores are sign-reversed so positive values indicate the failure direction. Phenotype uses prespecified consensus weights; its absolute-weight coverage is shown. Empty modules are reported as no scorable members. Sheet S3 in the accompanying Supplemental Tables workbook gives unrounded scores and comparisons.

**Supplemental Table S4. Keratocyte phenotype scores and reported in vivo outcomes.** A Concordance compares the direction of the consensus phenotype score with the reported outcome. Better corresponds to a positive score and worse to a negative score. Each row follows its reported outcome direction without significance filtering. All phenotype and decomposition scores are dimensionless; definitions are in Dictionary. B Losartan scores summarize stromal clusters 8, 12, 14 and 17; the keratocyte-enriched cluster 8 has a positive score, while clusters 12 and 14 have negative scores. The state and composition components use shared clusters 8, 12 and 14; the cluster-17 entry term is the remainder relative to their joint observation, with cluster 17 present only in the losartan arm. Anti-Gr-1 decomposition uses four lineages and tacrolimus as the reference, as in Figure S15. C At day 28 after freeze injury, Same direction refers to residual haze and negative phenotype relative to the same-day contralateral cornea. Relative to day 7, haze improves and phenotype approaches the uninjured state; this recovery comparison is reported separately from the same-day reference. D RNase III and the allograft-minus-unoperated-cornea contrast have no matched control outcome and are Not evaluable. The three anti-Gr-1 rows are Not paired and present the existing decomposition. Neither status contributes to outcome concordance. E In the ABT-263 experiment the scores shown are group-effect projections. Library-level consensus phenotype, as differences of group means (four libraries per group), was −0.2086 for ABT-263-treated and −0.2103 for vehicle-treated allografts relative to unoperated corneas, and +0.0017 between the two allograft groups. GEO, Gene Expression Omnibus; PBS, phosphate-buffered saline; PMID, PubMed identifier; RNase, ribonuclease; SMA, smooth muscle actin.

F Human rows apply the same group-level estimator to three patients versus four donor controls, eight controls (four donor and four public controls), or seven controls after merging the two public-control libraries A1 and H1. These rows have no matched reported outcome, are Not evaluable and do not contribute to outcome concordance.

**Supplemental Table S5. Within-population state, captured-RNA composition and aggregate transcriptional responses during anti-Gr-1 and losartan treatment.** The decomposition uses the original cell annotations, expression matrices and prespecified scores. F10 combines treated within-population state with reference RNA shares; F01 combines reference state with treated RNA shares; F11 combines treated state and treated RNA shares. All corners are compared with the same reference F00. State and composition contributions use the symmetric two-factor decomposition described in Supplemental Methods. The losartan cluster-17 entry term adds that treatment-only cluster after the shared-cluster comparison. RNA fractions use captured UMI, and cell fractions use quality-controlled cells. These fractions have separate denominators. Source-specific common-gene and all-eligible variants retain the original decomposition weights; the full z-standardized patient-test variants are reported separately in Supplemental Table S2.

**Supplemental Table S6. Keratocyte phenotype, matrix expression and remodeling programs during rabbit repair and fibrosis.** Group-level readouts retain two injured–contralateral rabbit pairs at each of days 7 and 28 after freeze injury and three fibrotic and three naive rabbits at day 14 after alkali burn. Failure-signature similarity, phenotype scores, core matrix expression, INHBA and SRF/MRTF readouts retain their original gene coverage and scales. Direct day-28-minus-day-7 comparisons use all eight freeze-injury libraries after joint normalization; per-rabbit injury-minus-contralateral results are listed separately. SRF/MRTF, serum response factor/myocardin-related transcription factor.

**Supplemental Table S7. Public intervention and induction readouts.** The 69-comparison analysis lists its dataset, species, treatment and comparator, time, replicate or pooled-library unit, and member coverage. Group and per-library readouts include failure-signature similarity, external axes, keratocyte phenotype, core matrix expression, complete rejection score and available-member score, glucocorticoid-receptor fingerprint and remodeling-gene programs. Treatment-minus-comparator orientation is retained for both per-library sets, which are summarized separately. Gene-rank projection statistics accompany the readouts to which they apply. Direct anti-Gr-1 comparisons and the population-specific losartan analysis are provided separately in Supplemental Table S11. GSE291819 is reported by sequencing-library unit.

**Supplemental Table S8. External datasets used for the main tissue-state and intervention comparisons.** Accessions, species, biological contrasts, sample or library units, analytic roles and linked publications are listed for the reference systems discussed in the main text. The complete public perturbation index belongs to Supplemental Table S7. ‡ GSE328779 is not yet linked to a publication in GEO; the PMID given matches the Series title, authors and design.

| Accession | Organism | Design | Use in this study | PMID |
|---|---|---|---|---|
| PRJNA1533946 | Homo sapiens | Four public controls (human corneal lenticules) | Eight-control featureCounts panel and prespecified-axis extension | — |
| GSE291819 | Mus musculus | Day-3 nitrogen-mustard injury with saline or losartan; one sequencing library per condition | Aggregate and cell-population phenotype responses linked to published haze and flow cytometry | 41537756 |
| GSE236626 | Rattus norvegicus | Corneal allografts: resveratrol and saline, three corneas per arm | Resveratrol versus saline intervention | 37937143 |
| GSE337743 | Oryctolagus cuniculus | Corneal cells in 2D and 3D cultures, three libraries per condition | Culture geometry and density comparisons | — |
| GSE231796 | Felis catus | Corneal fibroblasts, TGF-β induction and drug treatment, five culture replicates per group | Induction and post-induction drug comparisons | 37870848 |
| GSE87330 | Rattus norvegicus | Injured cornea, dexamethasone, anti-VEGF and IgG control, four microarrays per group | Dexamethasone versus IgG after injury | 28811496 |
| GSE213571 | Mus musculus | Adrenalectomized mice, dexamethasone and vehicle, four pooled libraries per group, 4–6 corneas per library | Dexamethasone versus vehicle in normal cornea | — |
| GSE151631 | Homo sapiens | Corneal buttons: 19 keratoconus, 7 control (bulk RNA-seq, HiSeq 2500) | Keratoconus-versus-healthy positive separation control; within-KC quartiles as a same-state null | 32555404 |
| GSE175889 | Mus musculus | Implant microenvironment, 19 libraries across ECM/PCL/saline/naive (scRNA-seq) | A3 implant-remodeling axis; PCL-versus-ECM stratification | 34341534; 37079217 |
| GSE181893 | Mus musculus | High-risk corneal transplantation with anti-CD80/86: 3 PBS and 3 anti-CD80/86 graft libraries at 2 weeks (bulk RNA-seq) | Broad A1 transplant-response reference and independent positive control for the five-module rejection score | 35318419 |
| GSE186433 | Homo sapiens | Eight healthy donor corneas, 19,472 cells (scRNA-seq) | Deconvolution reference | 34741068 |

| Accession | Organism | Design | Use in this study | PMID |
|---|---|---|---|---|
| GSE199013 | Homo sapiens | Anterior segment of six non-diseased donors (snRNA-seq) | Deconvolution reference; keratocyte phenotype weights | 35858321 |
| GSE227908 | Mus musculus | Implant capsule, mechanically stimulated versus standard, 2 and 4 weeks (scRNA-seq) | Mechanical-force axis; external comparator for the mechanotransduction branch | 37749310 |
| GSE260476 | Homo sapiens | Human corneal stromal fibroblasts and TGF-β1-induced myofibroblasts, 12 samples (bulk RNA-seq) | Human TGF-β1 myofibroblast state; myeloid-gene response control | 39240550; 39243928 |
| GSE275249 | Mus musculus | Corneal transplantation under tacrolimus or rapamycin, nine libraries (scRNA-seq) | A1 direction calibration (allograft+PBS versus isograft); external true-rejection scores; second allorejection resource for the core-matrisome comparison | 40398562 (tacrolimus/anti-Gr-1 design correspondence; GEO does not supply this PMID link) |
| GSE285060 | Homo sapiens | Five keratoconus and three control corneas (Visium spatial) | Second human corneal cohort for spot-module direction check | 39837422 |
| GSE287915 | Oryctolagus cuniculus | Rabbit corneal cells: serum-free, TGF-β1, FBS, 8 replicates each (bulk RNA-seq) | External pure-culture matrix for the myeloid-gene panel | 40072446 |
| GSE302936 | Oryctolagus cuniculus | Alkali-burn corneal fibrosis: day-14 fibrotic versus naive corneas, 3 versus 3 (scRNA-seq) | Rabbit alkali-burn fibrosis reference state (fibrotic versus naive stromal lineage) | 41147949 |
| GSE303036 | Homo sapiens | Eight keratoconus and five control corneas (Visium spatial) | Spatial localization of the failure signature; myeloid–stromal adjacency | 41931003 |
| GSE303805 | Mus musculus | Limbal injury time course: uninjured, day 2, 5, 10, 40; one pooled library per state (scRNA-seq) | A2 repair-versus-resolution axis (all-cell day 2 minus day 40; stromal day 5 and day 10 minus day 40); repair-corridor anchors | 42034629 |
| GSE306147 | Mus musculus | T cells in muscle injury and foreign-body response, 15 libraries (scRNA/TCR-seq) | Comparator for the adaptive branch in sorted T/γδ layers | 41997939 |
| GSE328779 | Oryctolagus cuniculus | Transcorneal freeze injury with non-fibrotic repair: day 7 and day 28, 2 injured and 2 paired control corneas each (scRNA-seq) | Rabbit nonfibrotic repair reference states (day 7 and day 28 injury versus control stromal lineage) | 41421445‡ |
| GSE39394 | Homo sapiens | Human fibroblasts, TGF-β-treated versus control, 4 samples (expression microarray) | Additional human fibroblast TGF-β system for the myeloid-gene panel | 23153532 |
| GSE6676 | Mus musculus | Mouse corneas, wild-type versus high-dose TGF-β, 8 samples (expression microarray) | Mouse cornea TGF-β system for the myeloid-gene panel | no linked publication in GEO |

| Accession | Organism | Design | Use in this study | PMID |
|---|---|---|---|---|
| GSE97358 | Homo sapiens | 168 human cardiac fibroblasts: 84 control, 84 TGF-β1 (bulk RNA-seq) | Second fibroblast system bounding the myeloid-signal interpretation | 29160304 |
| Human cornea meta-atlas (no single accession; integrated from four published scRNA-seq studies) | Homo sapiens | 65,036 cells, 21 cell states | Keratocyte phenotype weights (quiescent stromal keratocyte versus corneal fibroblast) | 40838019 |

**Supplemental Table S9. Prespecified tissue-program projections in public repair, fibrosis and perturbation systems.** Sheet S9 in the accompanying Supplemental Tables workbook retains each dataset, contrast, panel, mapped and total member count, ranked-gene universe, unaligned mean rank, failure-oriented mean rank and gene-set permutation summary. Positive scores follow the prespecified panel-level directions in the failed human lenticules. The transparency projection therefore has the opposite orientation to the biological display of transparency in Figure 3C. Null percentiles describe gene-rank position. Individual experimental units and group-level failure-signature similarity and phenotype readouts are identified in Figures 5–8, Figure S7 and Supplemental Table S7. Per-animal panel ranges are provided in Supplemental Table S11.

**Supplemental Table S10. Original-scale matrix effects and keratocyte phenotype responses within the rabbit and feline culture experiments.** Rabbit contrasts use GSE337743, with three culture libraries per condition; the common model includes all 15 libraries in the five original conditions. Original pairwise effects and the common-model effects are shown side by side. Positive scar ranks and positive original expression effects denote the failure-associated panel direction. Core-matrisome means retain their own member universe. The direct phenotype decomposition uses the original complete gene ranks and a common denominator for positive and negative weights. Feline day-5 contrasts use GSE231796, with five culture libraries in each of the three compared treatment arms. The three contrasts share these arms. Scores and $\log_2$ fold changes are reported with their original estimators and precision. The same-day feline comparisons are shown in Figure S14.

**Supplemental Table S11. Rabbit panel reconstructions and source-linked tissue outcomes.** Part A gives the per-animal panel ranges for the rabbit references. Parts B and C link the direct anti-Gr-1 and losartan transcriptional comparisons to their reported graft outcomes. Failure-signature similarity is symmetric normalized rank enrichment; the phenotype score is an externally weighted rank score. Panel values are mean ranks aligned with the prespecified panel direction, and the rejection score is the five-module calibrated projection. A positive transparency-panel score denotes change toward the transparency loss observed in failed lenticules. Source-specific phenotype definitions, treatment windows and sampling units are retained.

**Supplemental Table S12. Adjudicated clinical-report, treatment-sequence and specimen records.** Report identifiers and links retain procedure, implantation arm, event phenotype, attributed cause, treatment sequence, clinical outcome, tissue examination and known or unreported eye/specimen counts. Report, eye and specimen units remain separate. Duplicate and cohort links identify the relationship between the reviewed records and the final aggregation. Additional worksheets with the suffixes _02 to _07 list the same records with one row per implantation arm, event, treatment, specimen, eye, and duplicate or cohort link.

**Supplemental Table S13. Published corneal stromal implantation records, opacity courses and tissue examinations.** Counts retain report-record, eye and specimen units; modalities and event categories can overlap. One of the 285 adjudicated report records duplicates another and was removed, leaving 284 reports; the displayed counts are tallied over the 284 reports. Specimen totals sum known counts. Eleven specimen records did not report a specimen number. Treatment-sequence rows distinguish corticosteroid-only resolution, resolution after corticosteroids plus a procedure and spontaneous resolution. Source-level adjudication and report linkage are assigned to Supplemental Table S12.

**Supplemental Table S14. Software, references and analytic parameters.** Versions refer to the analysis environment used to generate the reported results. Alignment and quantification settings, repertoire references and dataset references identify the original in-house analysis and its external projections.

| Step | Tool | Version | Reference / database | Key parameters |
|---|---|---|---|---|
| Read QC | FastQC | v0.11.9 | — | default |
| QC aggregation | MultiQC | v1.35 | — | default |
| Alignment | HISAT2 | v2.2.1 | GRCh38.p12 | default |
| BAM handling | SAMtools | 1.13 | — | — |
| Variant utilities | BCFtools | 1.13 | — | — |
| Interval utilities | BEDtools | v2.30.0 | — | — |
| Gene counting | featureCounts (Subread) | v2.1.1 | GRCh38.p12 | -T 8 -p --countReadPairs -B -Q 10 -s 0 -t exon -g gene_id |
| Transcript quantification | Salmon | 1.4.0 | GRCh38.p12 | -l A (returned IU for all twelve libraries) |
| Differential expression | edgeR | v4.6.2 | — | patient-level pseudobulk; the unit of inference is the patient |
| Gene-set enrichment | fgsea | v1.34.0 | MSigDB Hallmark | — |
| Module scoring | UCell | v2.12.0 | — | — |
| Deconvolution | BayesPrism | v2.2.3 | GSE186433; GSE199013 | broad and fine label tiers |
| Deconvolution | MuSiC | v1.0.0 | GSE186433; GSE199013 | broad and fine label tiers |
| Spatial deconvolution | RCTD | v2.2.1 | GSE303036 | reference-anchored cell abundances |
| Immune repertoire | TRUST4 | v1.1.10-r639 | IMGT reference human_IMGT+C.fa distributed with TRUST4 (SHA-256 f9bb920092a2b4aab4b17d81057f8e1cda35403e743123d4f9c39b02a4c2a8a2) | common depth 16,028,223 read pairs; six-part clonotype reliability gate (Supplemental Methods) |
| Taxonomic screen | Kraken2 | v2.17.1 | PlusPF, build pluspf_20260626 | — |
| Statistical environment | R | 4.5.0 | — | — |
| Scripting | Python | 3.10.12 | — | BLAS threads capped at 1 (OMP/MKL/OPENBLAS/VECLIB) |

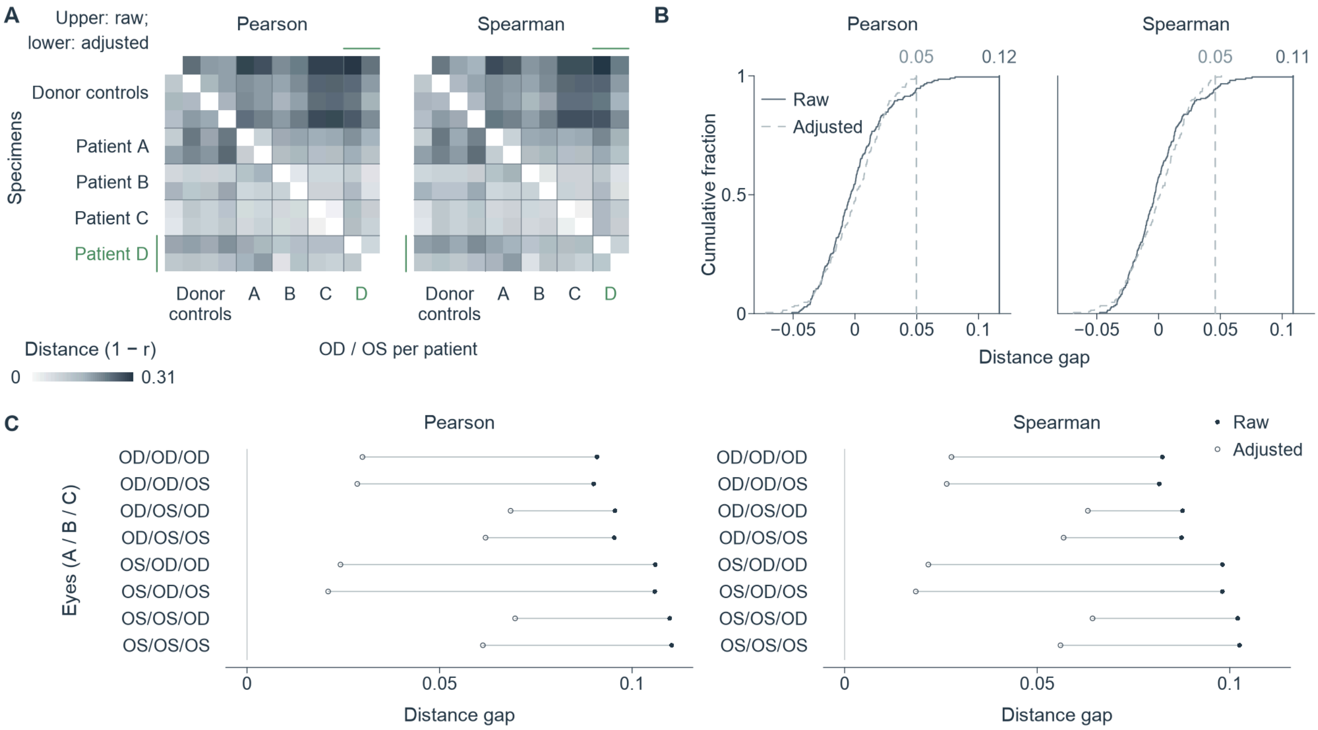


**Supplemental Fig. S1. Failed lenticules remain transcriptomically distinct across single-eye selections.**

(A) Pearson and Spearman distances (1 − correlation) among four donor controls and bilateral specimens from Patients A–D. Upper/lower triangles show raw distances/distances adjusted for mitochondrial fraction and expression complexity, on a shared white-to-dark scale. Patient D, shown in green, supplied two lenticules removed for another indication and sequenced separately. (B) Exact label-allocation cumulative distributions for Patients A–C and the four controls, with 210 allocations per metric and adjustment. The gap is mean failure–control distance minus mean within-failure distance. Solid/dashed curves and observed vertical lines denote raw/adjusted values. (C) Raw (filled) and adjusted (open) gaps for all eight selections of one eye per patient, shown separately for Pearson and Spearman distances. Row labels specify eyes in Patient A/B/C order. OD/OS, right/left eye.

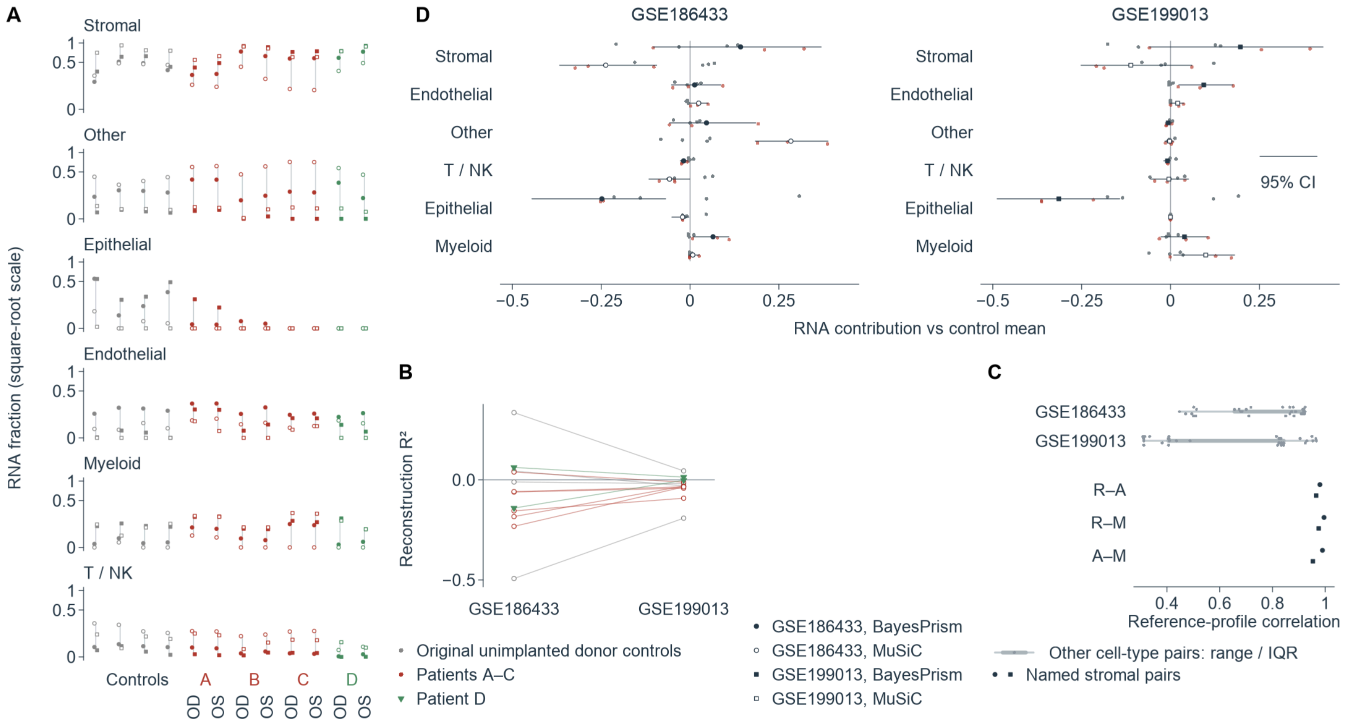


**Supplemental Fig. S2. Captured-RNA contribution estimates vary with reference and method.**

(A) Six-class captured-RNA fractions for 12 specimens: four original unimplanted donor controls and both eyes of Patients A–D. Patient D supplied two lenticules removed for another indication and sequenced separately. Circles/squares identify GSE186433/GSE199013; filled/open symbols identify BayesPrism/MuSiC. Lines span four estimates per specimen; square-root axes retain fraction labels. Gray, red and green identify controls, Patients A–C and Patient D. (B) MuSiC reconstruction $R^2$, paired across references for each specimen. Green inverted triangles identify Patient D. Sixteen of 20 primary-cohort fits and two of four Patient D fits have negative $R^2$. (C) Pearson correlations between log-transformed counts-per-million reference profiles. Dark symbols show the three named stromal pairs in each reference; gray points, thin ranges and thick interquartile segments summarize other pairs. R, resting keratocyte; A, activated keratocyte; M, myofibroblast candidate. (D) Failed-minus-control RNA fractions for each reference–method–class combination, with 95% percentile bootstrap intervals resampling three patients and four controls after combining eyes within patient. Small red/gray points show those patient/control fractions centered on the corresponding control mean. Each reference dataset is shown as a facet, with six cell types and paired method rows identified by the shared filled/open-symbol key. NK, natural killer; OD/OS, right/left eye.

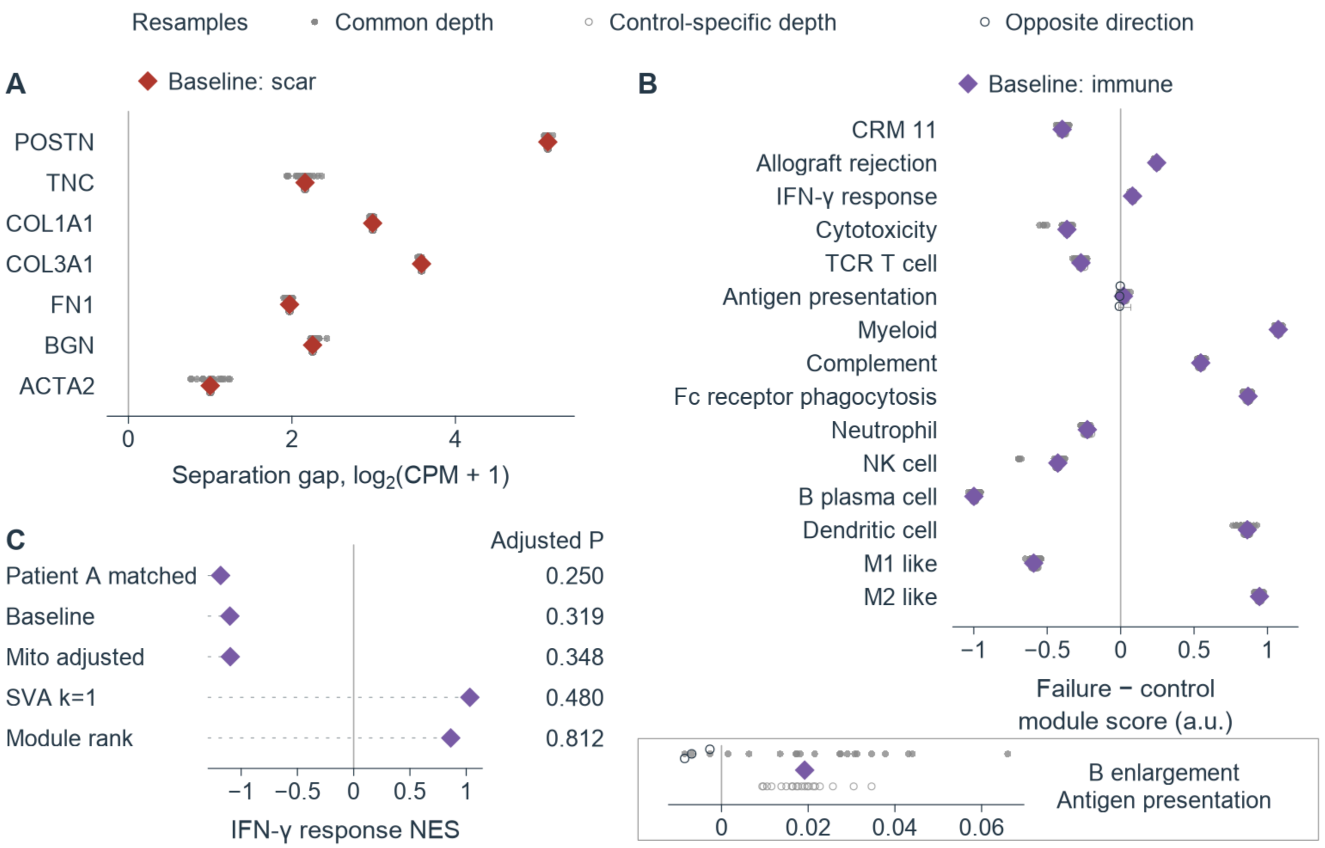


**Supplemental Fig. S3. Scar matrix separation persists after sequencing-depth resampling.**

(A) Minimum failed-lenticule minus maximum control expression for seven scar genes in six failed lenticules and four original unimplanted donor controls, using trimmed-mean-of-M-values-normalized log2(CPM + 1). (B) Failure-minus-control differences in 15 immune-program mean gene z scores. Filled/open gray points in A and B denote common-depth/control-specific-depth resamples; red/purple diamonds denote scar/immune baseline estimates. There are 20 technical resamples per gene or program and scenario. Thin lines in A span observed resampling ranges. Dark-gray open circles mark antigen-presentation estimates opposite to baseline; the boxed enlargement in B shows the same antigen-presentation resamples and baseline. (C) IFN-γ response normalized enrichment scores under five conventions: Patient A batch-matched, baseline, mitochondrial-fraction-adjusted, one-surrogate-variable adjustment and module enrichment on edgeR gene ranks. Purple diamonds show each estimate; three are negative and two positive. The adjacent column gives the adjusted P value for each convention. CPM, counts per million; TCR, T cell receptor; NK, natural killer; CRM 11, 11-gene common rejection module; Fc, fragment crystallizable; IFN-γ, interferon-γ; Mito, mitochondrial; SVA, surrogate variable analysis; k, number of surrogate variables; NES, normalized enrichment score; a.u., arbitrary units.

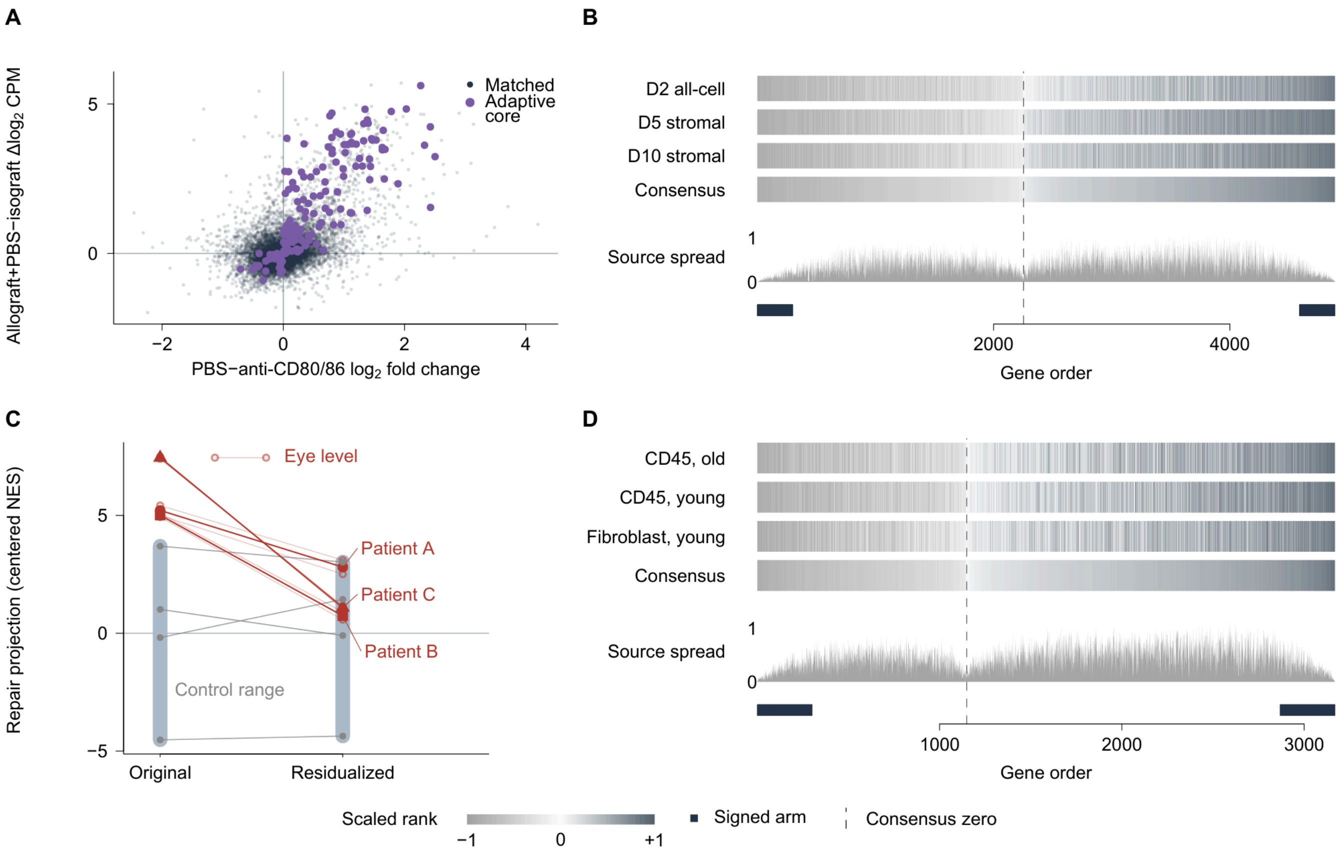


**Supplemental Fig. S4. Repair projections overlap controls after shared reference structure is removed.**

(A) Effects for 11,390 matched genes in PBS minus anti-CD80/86 (horizontal, edgeR log2 fold change) and allograft plus PBS minus isograft (vertical, Δlog2CPM). Purple points identify 145 concordant adaptive-core genes; gray points show the remaining matched genes. (B) Repair-reference ranks from day-2 all-cell, day-5 stromal and day-10 stromal contrasts against day 40. (C) Original and residualized repair projections in three paired-eye patient medians and four original unimplanted donor controls. Red circles, squares and triangles identify Patients A, B and C; pale red lines with open circles show the six eyes (eye level), gray points show controls, and blue-gray bars span control ranges. Lines join the same unit across projections. Scores average forward and reciprocal normalized enrichment scores and subtract the control mean. Reference weights were mutually residualized in their shared gene space before projection. (D) Implant-remodeling reference ranks from polycaprolactone minus extracellular matrix in CD45-enriched old/young and young-fibroblast strata. In B and D, aligned bands show source ranks and their consensus; gray, white and dark denote −1, 0 and +1. Gray areas show maximum-minus-minimum source rank; dark rugs identify the 300 genes in each signed arm, and dashed lines identify consensus ranks nearest zero. PBS, phosphate-buffered saline; CPM, counts per million; CD45, cluster of differentiation 45; NES, normalized enrichment score.

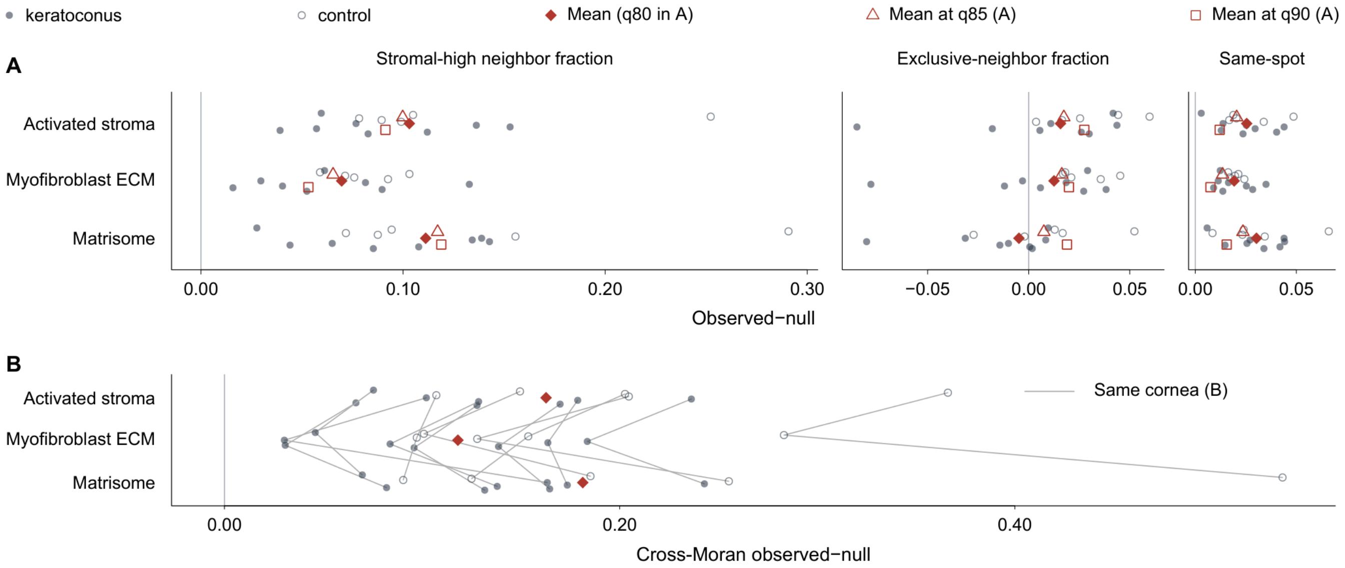


**Supplemental Fig. S5. Myeloid and stromal programs share spatial neighborhoods in human corneas.**

(A) Spatial associations between the myeloid/phagocytosis/complement program and activated-stroma, myofibroblast extracellular matrix (ECM) or matrisome programs in 13 corneas. Filled/open gray circles denote keratoconus/control corneas. Stromal-high neighbor fraction measures directed edges from myeloid-high to stromal-high spots; exclusive-neighbor fraction removes dual-high spots from both endpoint classes; same-spot fraction measures dual-high spots among all spots. Individual corneas and red diamonds show values and equal-cornea means at the 80th-percentile threshold; open red triangles/squares show means at the 85th/90th percentiles. Values subtract within-cornea permutation means. (B) Continuous cross-Moran observed-minus-null values for the same programs and corneas, calculated from standardized scores across neighboring spots. Lines connect programs within cornea; red diamonds show equal-cornea means. Vertical lines mark zero.

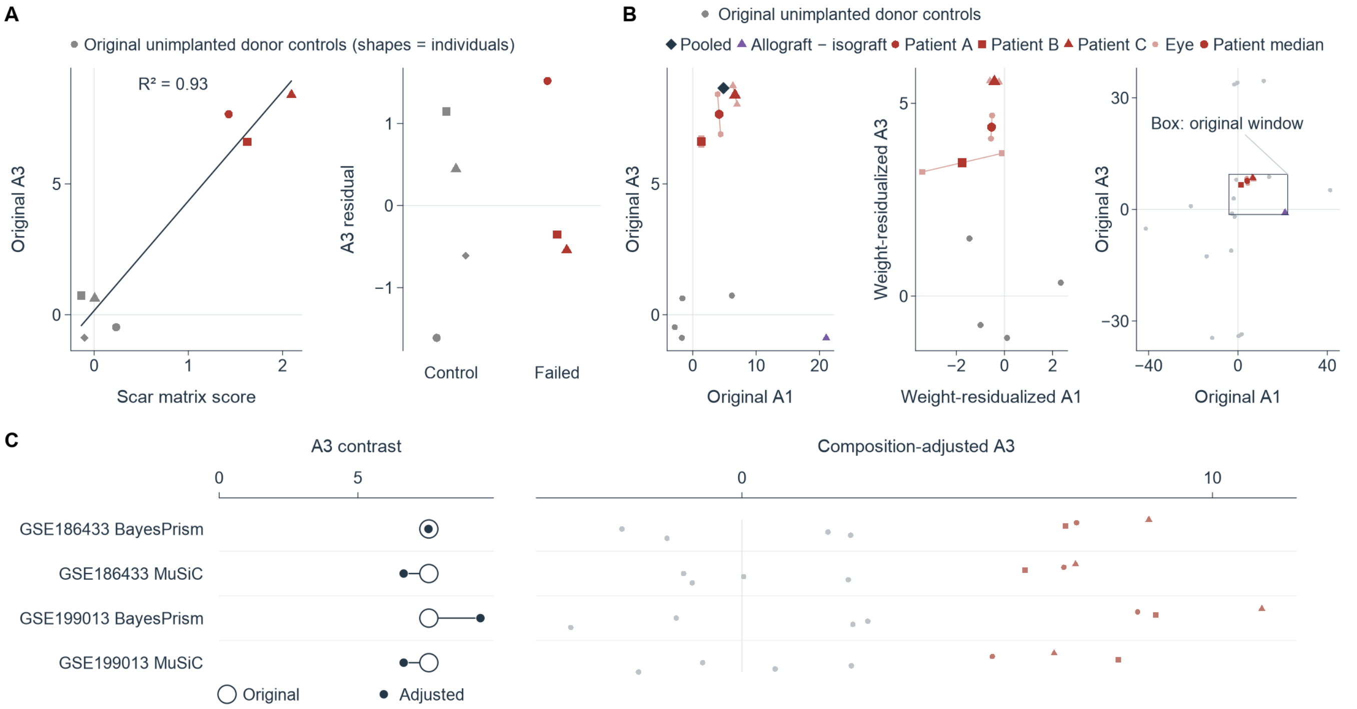


**Supplemental Fig. S6. External implant-remodeling scores track scar matrix and retain separation after composition adjustment.**

(A) Implant-remodeling score (A3) versus scar matrix score in three paired-eye patient medians and four original unimplanted donor controls. The line shows linear regression ($R^2 = 0.93$); the adjacent strip shows residuals. Gray shapes distinguish controls; red circles, squares and triangles identify Patients A–C. (B) Broad transplant-response (A1) versus implant-remodeling (A3) scores before and after mutual residualization of the three external reference weights. Pale/dark red symbols show six eyes/three patient medians; lines join eyes within patient. In the original view, the dark diamond shows the pooled failure contrast and the purple triangle the allograft-minus-isograft reference. The right plot places 18 external states on the original scores; the box marks the original display window. (C) Original (open rings) and composition-adjusted (filled points) failure-minus-control contrasts for two references and two methods. The aligned right-hand rows show seven adjusted patient or donor scores per reference–method combination. Zero lines indicate no change.

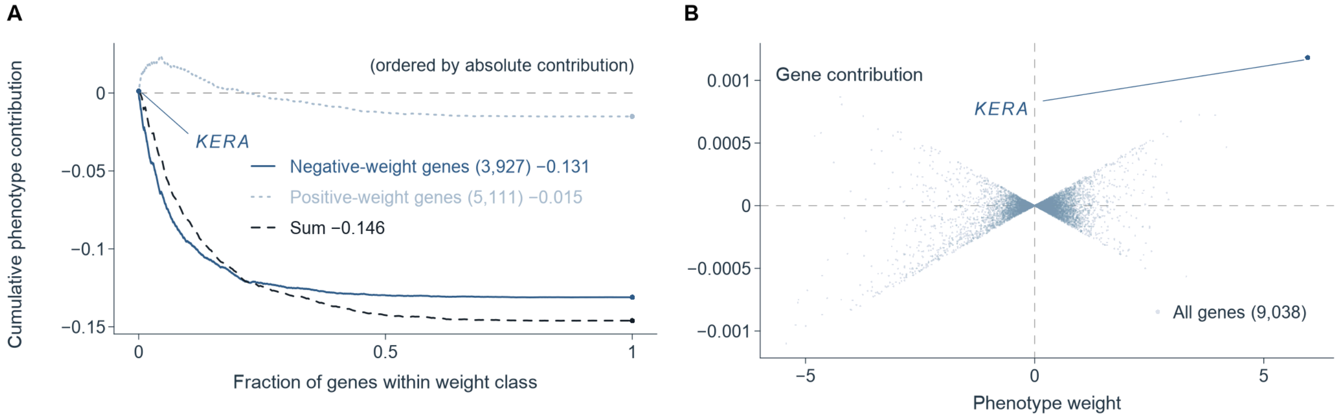


**Supplemental Fig. S7. Phenotype decline at higher culture density is distributed across genes with both weight signs.**

(A) Cumulative contributions to the consensus phenotype change for 100,000 minus 50,000 rabbit corneal cells seeded per three-dimensional collagen matrix, with three culture libraries per condition. Each gene contributes its phenotype weight multiplied by its scaled effect rank, divided by the sum of available absolute weights. Genes are ordered by absolute contribution within the negative- and positive-weight classes. The dark-blue solid and light-blue dotted curves show the 3,927 negative-weight genes and 5,111 positive-weight genes, respectively; the darker dashed curve sums both classes at matched within-class fractions. Endpoints are −0.131 and −0.015, totaling −0.146. KERA is marked on the positive-weight curve. (B) Phenotype weight versus signed contribution for the same 9,038 genes, with KERA labeled. Dashed lines mark zero. The aggregate decomposition is in Fig. 7C.

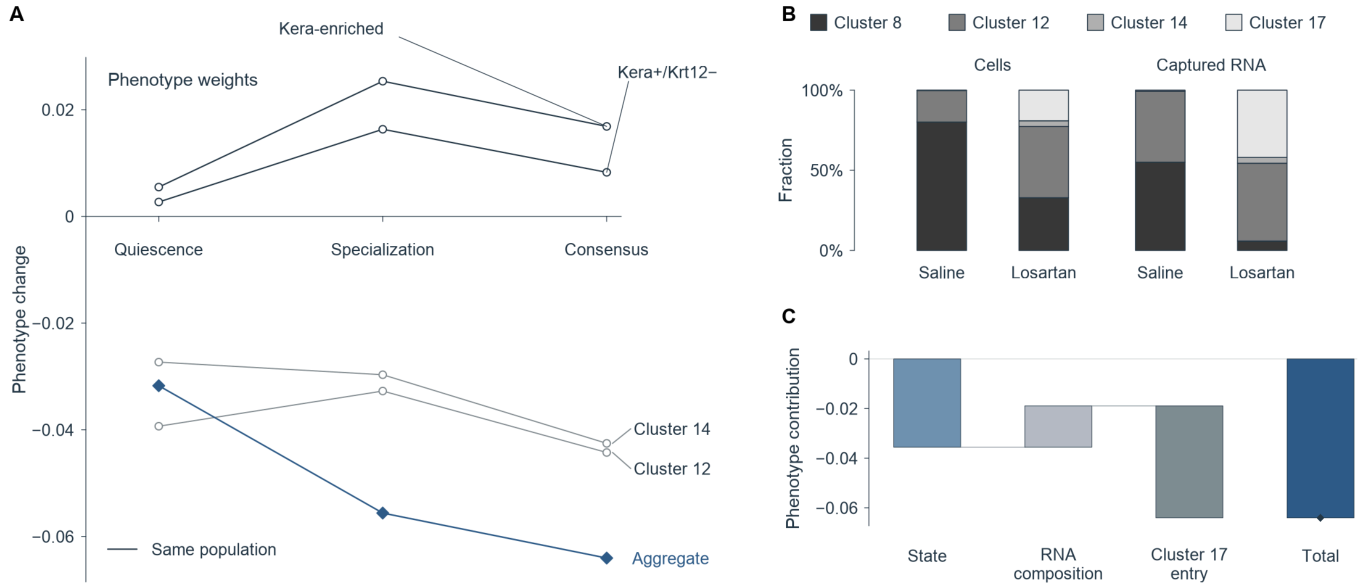


**Supplemental Fig. S8. RNA composition and cluster entry explain the lower stromal aggregate phenotype after losartan.**

(A) Losartan minus saline at day 3 after nitrogen mustard injury, with one sequencing library per condition. Lines connect five populations across quiescence, specialization and consensus weights: the four-cluster stromal aggregate, keratocyte-rich cluster 8, its Kera-positive/Krt12-negative subset, and clusters 12 and 14. The subset has Kera > 0 and Krt12 = 0 captured unique molecular identifiers. Blue diamonds/open circles show the aggregate/individual populations. (B) Cell fractions and captured-RNA fractions within the four-cluster aggregate, using each condition's cell or captured-RNA total as its denominator. Gray shades identify clusters 8, 12, 14 and 17. (C) Signed additive bridge from state (blue) through RNA composition (light gray) and cluster-17 entry (dark gray) to the total phenotype change (dark blue), marked by a filled diamond. State and composition average marginal changes over both replacement orders for shared-cluster profiles and RNA shares; cluster 17 is added last. Horizontal lines mark zero and connect cumulative levels. The aggregate comparison is in Fig. 5D.

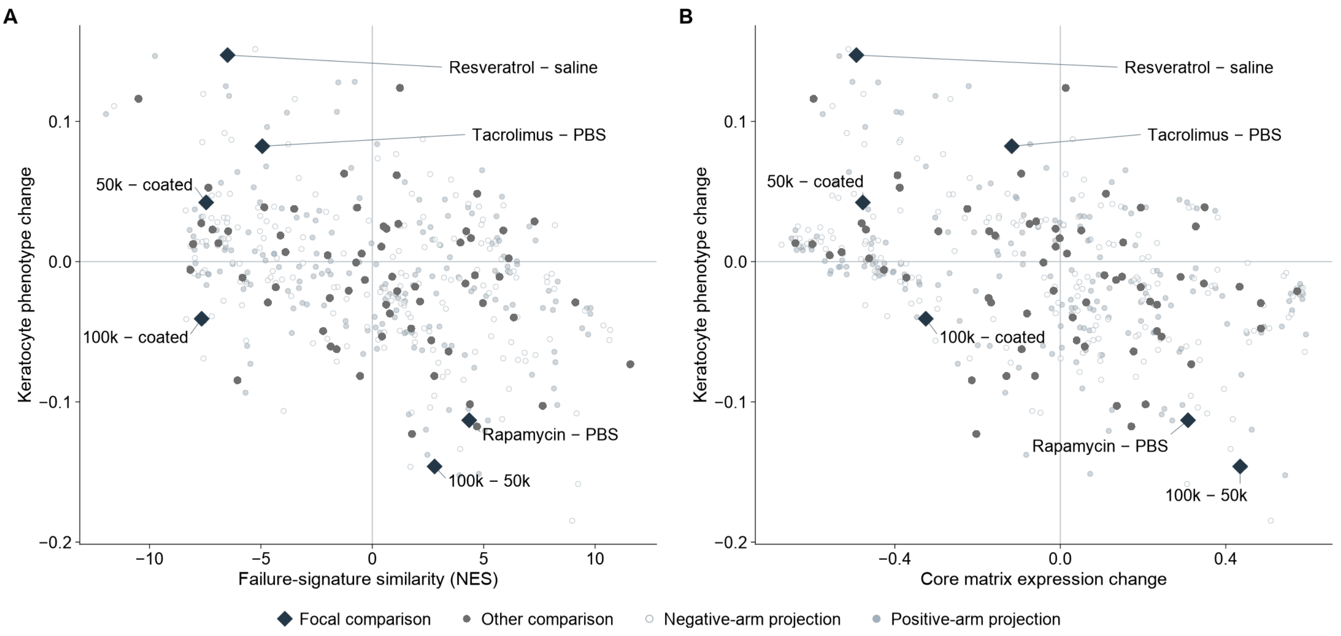


**Supplemental Fig. S9. Phenotype responses separate from failure-signature similarity and matrix responses across interventions.**

(A) Failure-signature similarity and (B) core matrix expression change versus phenotype change for 69 intervention and induction comparisons. Each large point represents one dataset–contrast group estimate in its original positive-minus-negative-arm direction. Dark diamonds identify six focal culture and drug contrasts; gray points show the other comparisons. Here 50k/100k denote 50,000/100,000 cells per collagen matrix, and coated denotes the coated culture comparator. Pale filled/open points show positive-/negative-arm library projections: 362 projections from 43 contrasts in each panel. Projections compare a positive-arm library with the negative-arm mean or the positive-arm mean with a negative-arm library. Group estimates and library projections are scored separately; shared libraries retain their unit identity across contrasts. Zero lines mark no change. NES, normalized enrichment score; PBS, phosphate-buffered saline.

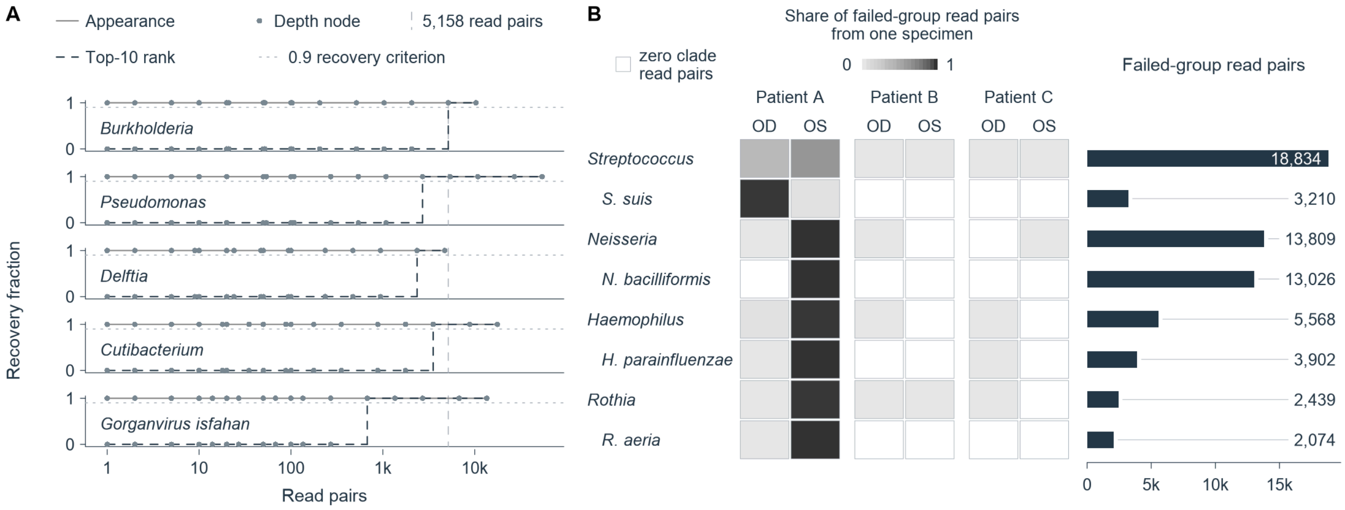


**Supplemental Fig. S10. Non-host taxon recovery depends on read depth and enriched reads concentrate in individual specimens.**

(A) Recovery of five taxon–source series under real-read resampling: Burkholderia, Pseudomonas, Delftia and Gorganvirus isfahan from original unimplanted donor controls, and Cutibacterium from Patient A OS. Solid/dashed steps show appearance/top-10-rank recovery fractions; each node summarizes ten technical resamples. The dotted horizontal reference marks 0.9 recovery; the dashed vertical reference marks the common floor of 5,158 non-host read pairs. Read-pair depth uses a logarithmic axis. (B) Eight taxa enriched in the six failed eyes and annotated as known background in the source classification. Each cell shows one eye's share of that taxon's failed-group clade read pairs, from white to black at 0 to 1; blank outlined cells denote zero clade read pairs. Aligned bars show failed-group totals. Genus and species rows represent nested taxonomic ranks; clade read-pair counts are not additive across these rows. Each taxon's largest share came from Patient A. N. bacilliformis, Neisseria bacilliformis; H. parainfluenzae, Haemophilus parainfluenzae; S. suis, Streptococcus suis; R. aeria, Rothia aeria; OD/OS, right/left eye.

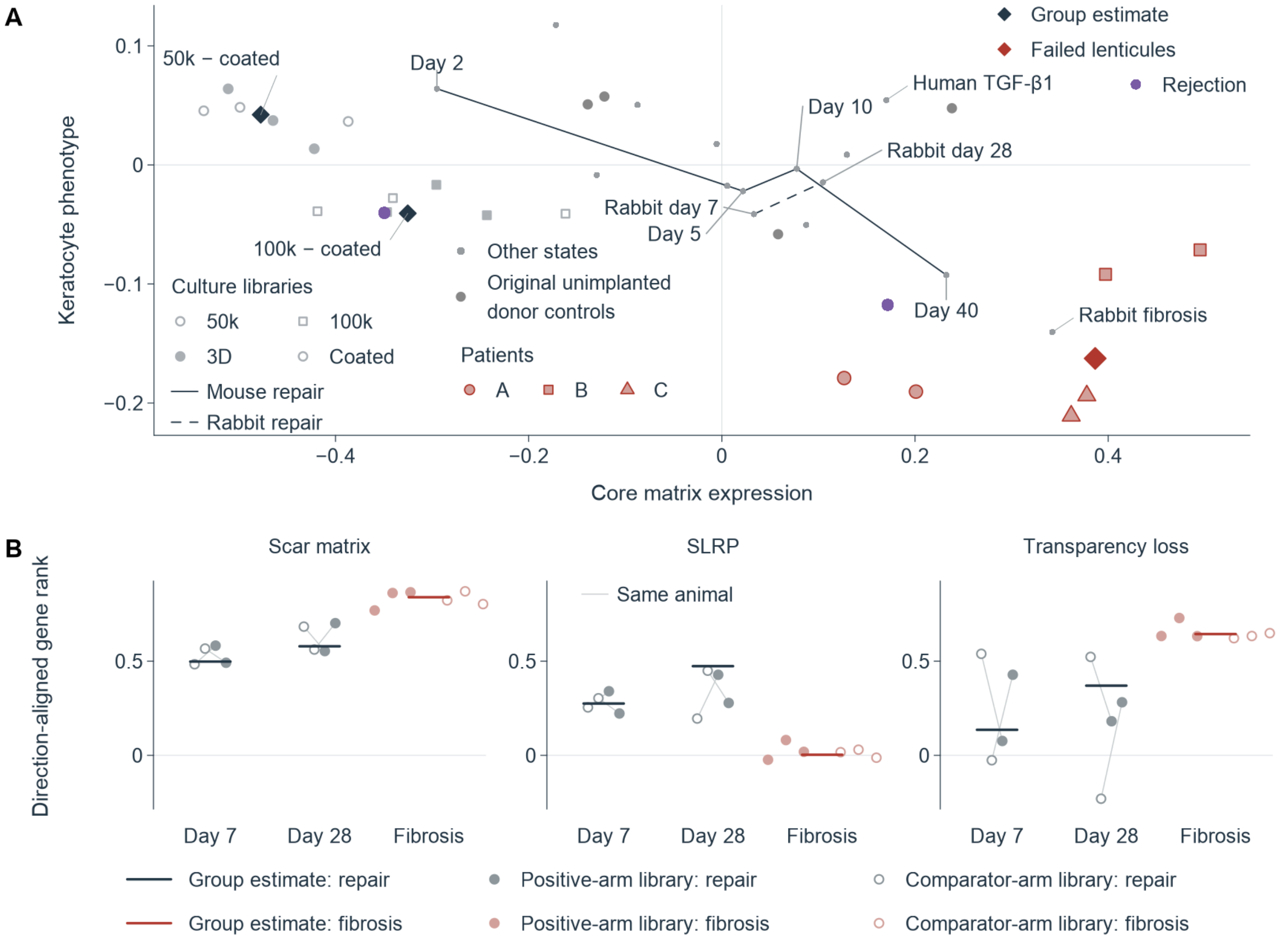


**Supplemental Fig. S11. Tissue phenotype separates from core matrix expression across repair and disease states.**

(A) Core matrix expression versus phenotype for 17 external states, comprising 15 other states and two rejection references, together with failed lenticules, four original unimplanted donor controls and two rabbit culture-density contrasts. Purple/gray points identify rejection references/other states; the red diamond identifies the failed-lenticule group contrast, and pale red circles, squares and triangles identify Patient A, B and C eyes. Dark diamonds show culture group estimates; library circles/squares denote 50,000/100,000 cells per matrix, and filled/open symbols denote 3D/coated projections. Three libraries per condition yield 12 projections from nine unique libraries. Solid/dashed lines connect mouse/rabbit repair group estimates. (B) Direction-aligned ranks for scar matrix, stromal small leucine-rich proteoglycan (SLRP) and transparency loss in rabbit freeze repair at days 7 and 28 and day-14 alkali fibrosis. Each repair time has two injured–contralateral animal pairs linked by pale lines; fibrosis has three libraries per arm. Dark/red marks show repair/fibrosis group estimates; pale filled/open points show positive-library minus comparator-mean/positive-mean minus comparator-library projections. Positive transparency ranks denote transparency loss. TGF-β1, transforming growth factor-β1; 3D, three-dimensional.

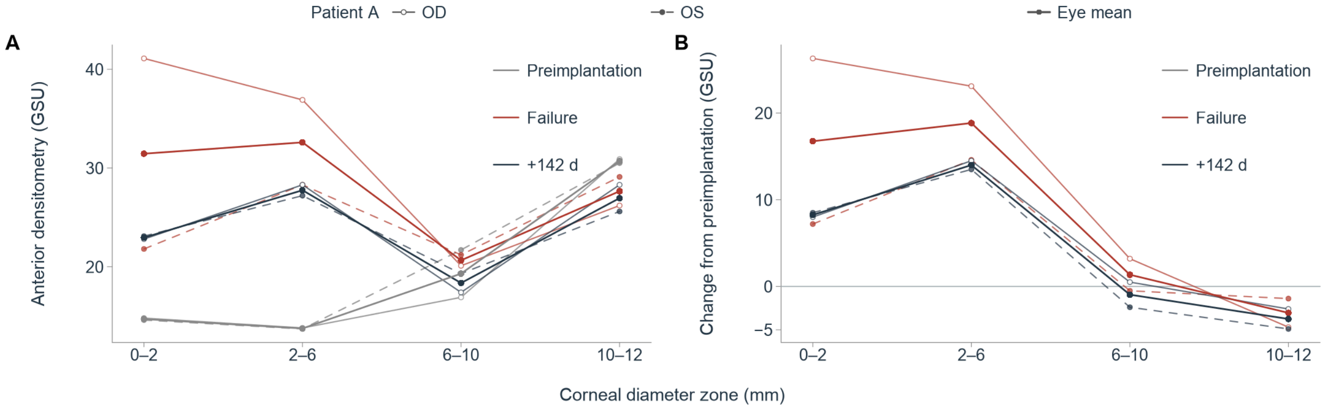


**Supplemental Fig. S12. Anterior corneal density remains elevated centrally after exchange in Patient A.**

(A) Anterior 120 µm Scheimpflug densitometry in both eyes of Patient A before implantation, at failure and +142 days after exchange. Gray, red and dark profiles identify the three stages. Pale points and lines show individual eyes; dark points and lines show their equal-eye means (two eyes, one patient). Open circles/solid lines denote OD; filled circles/dashed lines denote OS. The measured diameter zones are 0–2, 2–6, 6–10 and 10–12 mm. (B) Failure and late values minus each eye's preimplantation density in the same zone. Dark profiles average bilateral changes; the horizontal line marks zero. Central depth profiles are in Fig. 2D. OD/OS, right/left eye; GSU, grayscale units.

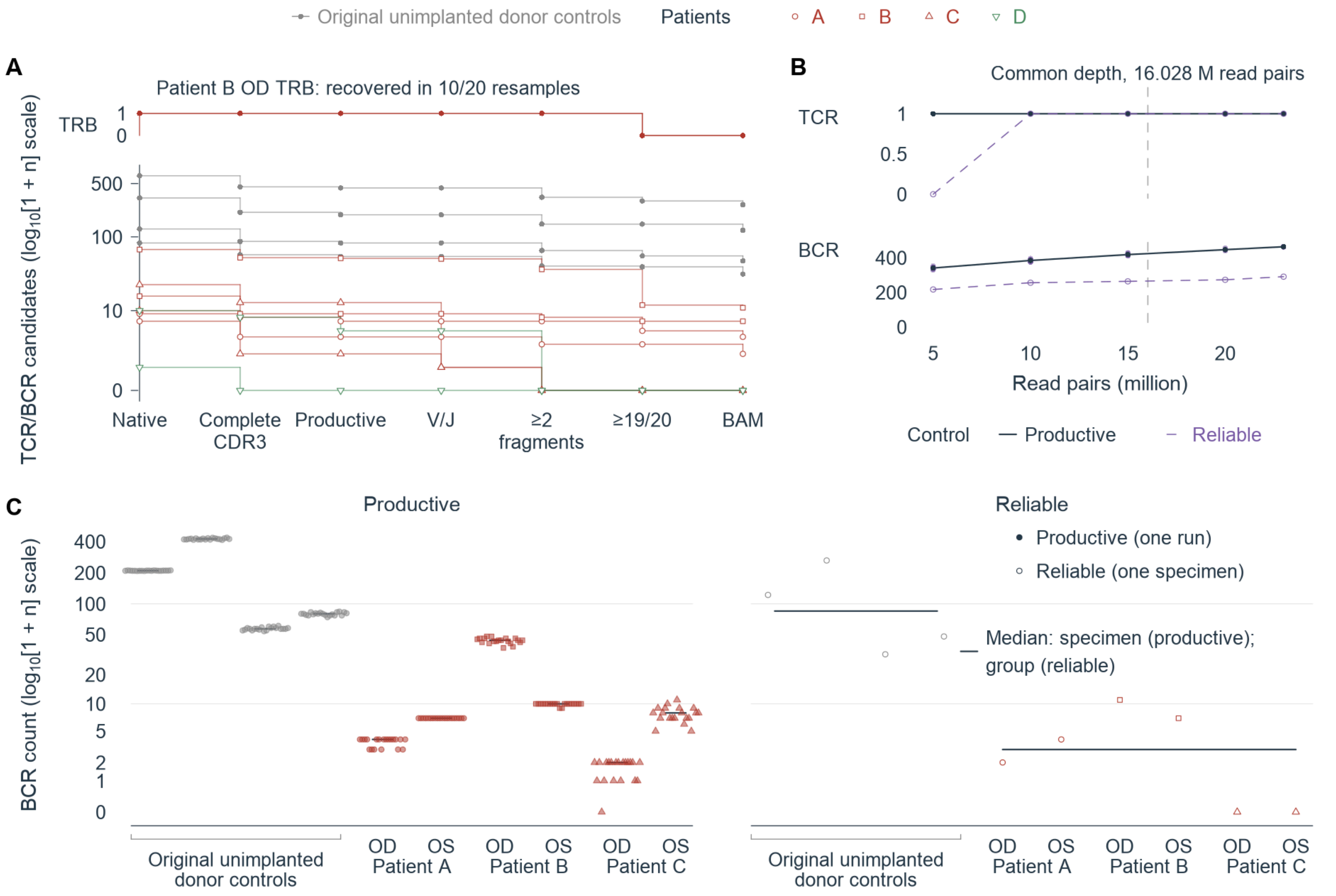


**Supplemental Fig. S13. Failed lenticules yield fewer reliable receptor clonotypes than donor controls.**

(A) T cell receptor (TCR) and B cell receptor (BCR) candidate retention across seven reliability stages in 12 specimens: four original unimplanted donor controls and both eyes of Patients A–D. Patient D supplied two lenticules removed for another indication and sequenced separately. Gray curves identify controls; red circles, squares and triangles identify Patients A–C; green inverted triangles identify Patient D. Stages proceed from native reconstruction through complete complementarity-determining region 3 (CDR3), productive sequence, chain-consistent variable/joining (V/J) assignments, support from at least two fragments, recovery in at least 19 of 20 common-depth technical runs, and matching receptor sequence from binary alignment/map (BAM) input. The upper track follows the Patient B OD TCR β candidate, recovered in 10 of 20 resamples. Counts use $\log_{10}(1 + \text{count})$ spacing. (B) Productive and reliable TCR and BCR counts at five sequencing depths in one donor control. Pale points show 20 technical runs per depth and receptor class; dark lines/ranges show productive medians/extrema; purple dashed lines/open points show reliable counts. The vertical line marks 16.028 million read pairs. (C) Productive and reliable common-depth BCR counts in four controls and six failed eyes from three patients. Filled points show 20 productive technical runs per specimen with specimen-median marks; open points show reliable counts per specimen. Horizontal lines show reliable group medians. Patient D appears in A; C uses Patients A–C and controls. OD/OS, right/left eye; TRB, TCR β.

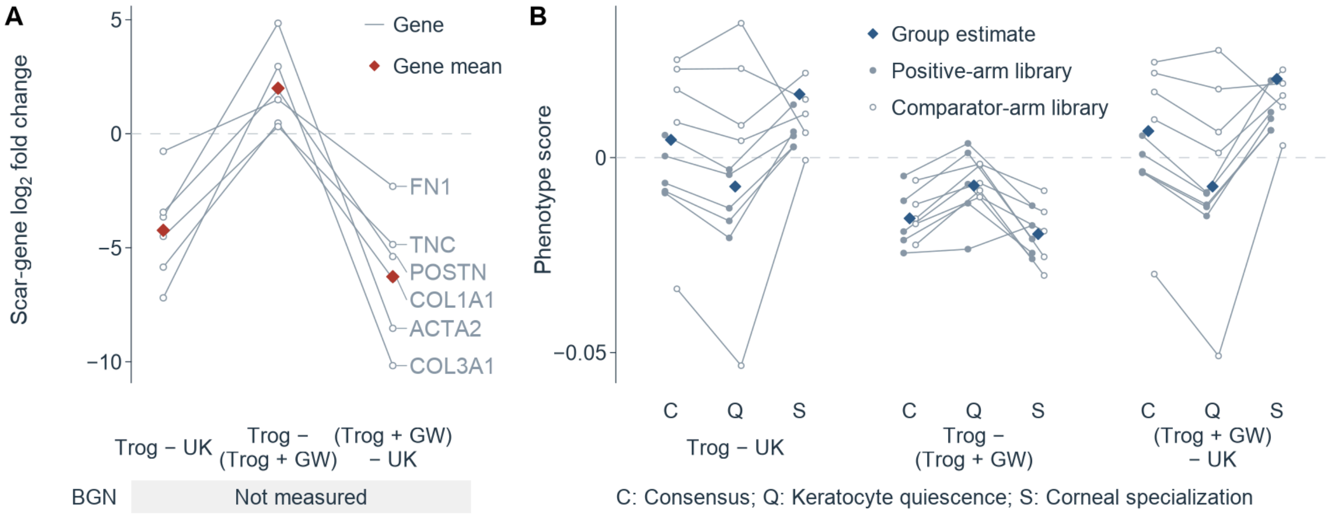


**Supplemental Fig. S14. Same-day drug contrasts separate scar matrix changes from phenotype responses.**

Three day-5 feline corneal-cell comparisons reuse 15 culture libraries, five per treatment arm. (A) Scar matrix gene $\log_2$ fold changes for troglitazone minus UK-5099, troglitazone minus troglitazone plus GW9662, and troglitazone plus GW9662 minus UK-5099. Lines connect the same six measured genes across contrasts, with gene names at their right endpoints; red diamonds show gene means. The BGN row identifies unmeasured expression. (B) Phenotype under consensus (C), keratocyte-quiescence (Q) and corneal-specialization (S) weights. Diamonds show group estimates; small filled/open points show positive-library minus comparator-mean/positive-mean minus comparator-library projections, with lines connecting the same library across weights. Each contrast has ten projections per weighting. Dashed lines mark zero. Trog, troglitazone; GW, GW9662; UK, UK-5099.

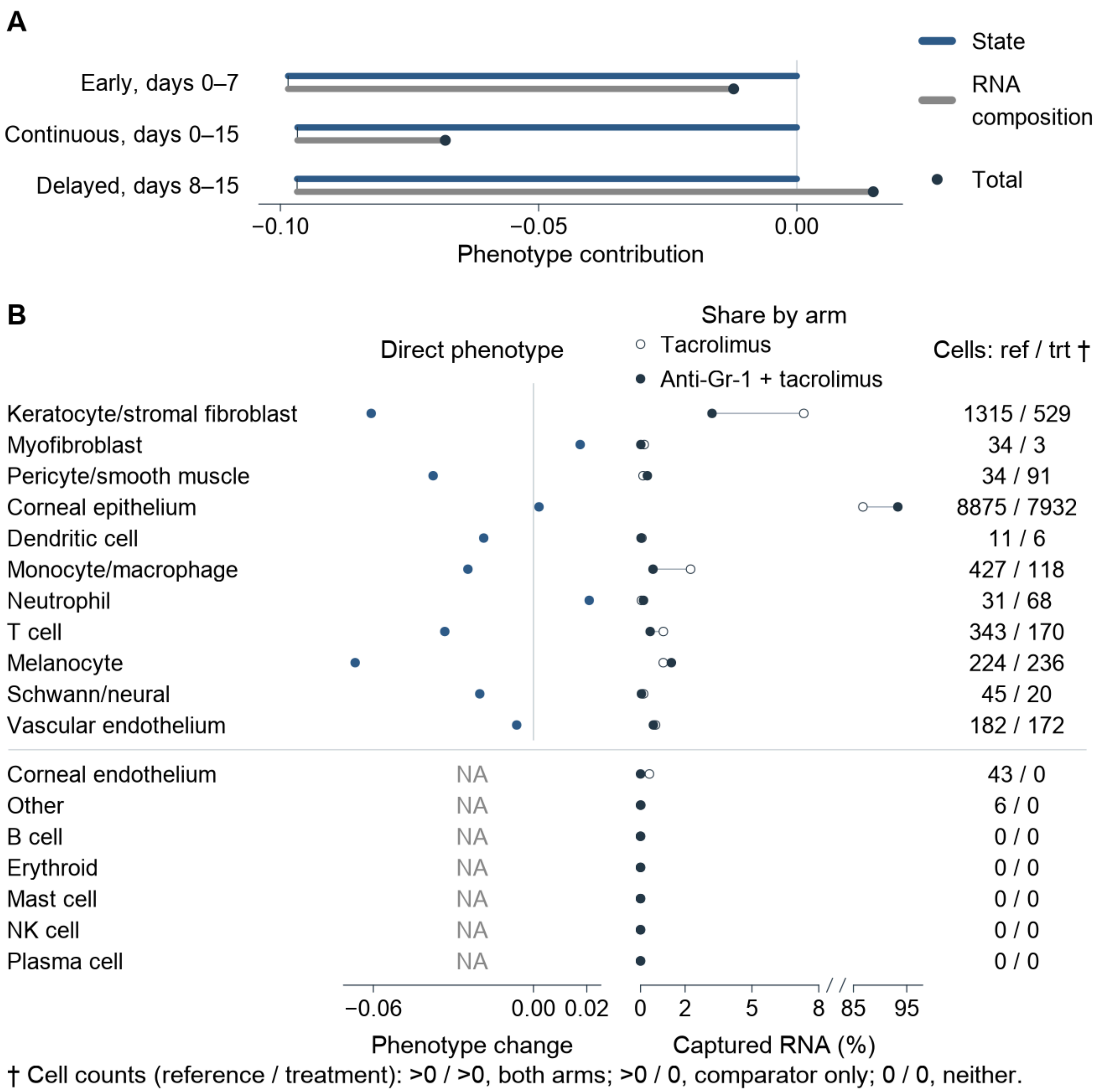


**Supplemental Fig. S15. Captured-RNA composition accounts for the higher aggregate phenotype after delayed anti-Gr-1 depletion.**

(A) Four-population decomposition of phenotype after early, continuous or delayed anti-Gr-1 plus tacrolimus versus tacrolimus alone. Blue state segments begin at zero; gray composition segments extend to totals marked by dark points. Contributions average marginal changes over both replacement orders for profiles and captured-RNA shares in T/NK, myeloid, stromal and other populations. RNA was collected on day 15 after days 0–7, 0–15 or 8–15 exposure. Each condition is one library pooled from 4–6 corneas. (B) Direct within-type phenotype effects, captured-RNA fractions, cell counts and observed support for all 18 annotated cell types in the delayed comparison; the upper block shows the 11 types with direct estimates, and the lower block retains the seven types without a direct estimate. Open/filled fraction points denote tacrolimus/anti-Gr-1 plus tacrolimus; lines join the same type. Captured-RNA shares are displayed on a broken axis, omitting the unobserved interval from 8% to 85%; tick labels give the original percentages. Cell counts are comparator/treatment. The † footnote below the horizontal-axis titles defines cell-count patterns for types present in both arms, in the comparator only or in neither arm; individual NA labels identify types without a direct score. Fine-cell-type attribution and the common-type scatter are in Fig. 6C and D. NK, natural killer; NA, not calculable from both arms.

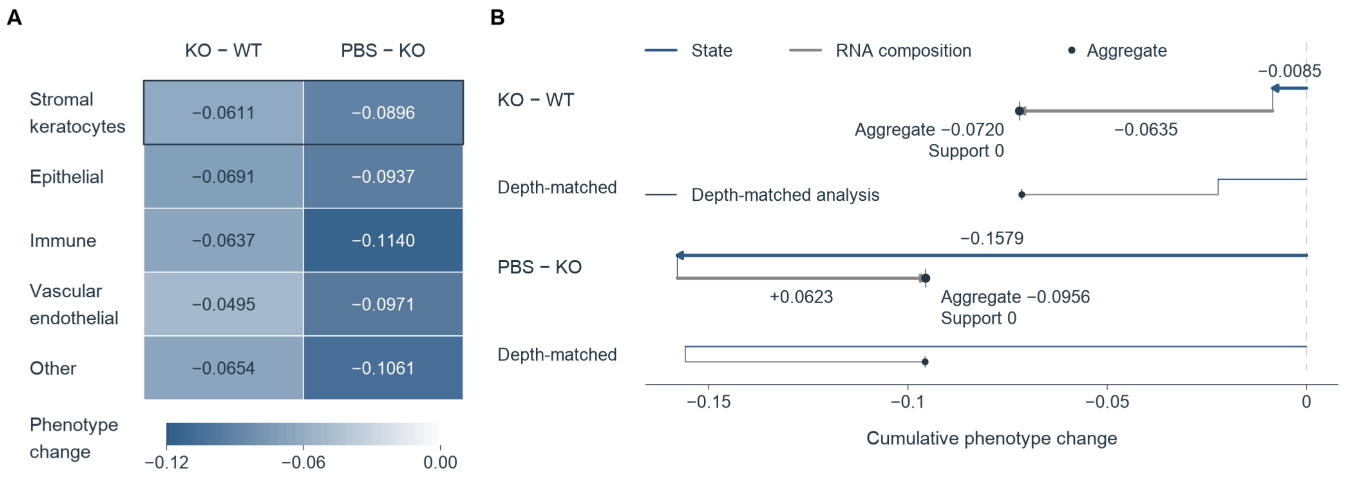


**Supplemental Fig. S16. Rewetting lowers stromal-lineage phenotype in Aire-deficient corneas despite restored matrix organization.**

(A) Consensus phenotype changes across five lineages in Aire knockout (KO) minus wild type (WT) and PBS-rewetted minus untreated KO corneas. The stromal-keratocyte lineage is outlined. Blue and white indicate negative and zero changes, respectively, on a shared scale from −0.12 to 0. (B) Whole-tissue phenotype decomposition into state (blue), captured-RNA composition (gray) and support terms. Thick/thin cumulative traces show main/depth-matched analyses. Arrows preserve signed addition, filled circles mark aggregate changes, and terminal vertical marks denote zero support terms. Each condition is one pooled nuclear library. PBS, phosphate-buffered saline.

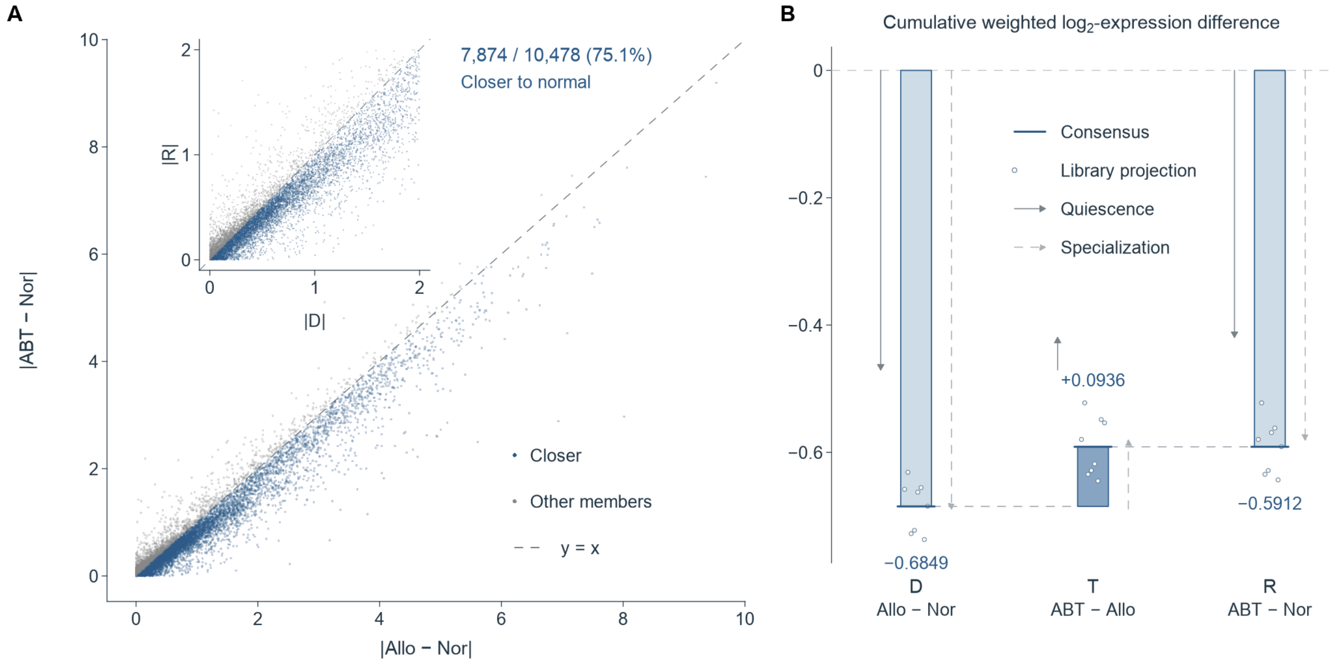


**Supplemental Fig. S17. ABT-263 moves most measurable phenotype genes toward normal while a residual expression difference remains.**

(A) Absolute allograft-minus-normal differences (D) versus absolute ABT-263-minus-normal differences (R) for all 10,478 measurable phenotype members. Blue points satisfy $|R| < |D|$: 7,874 members (75.1%), accounting for 75.3% of measurable absolute consensus weight. Gray points identify other members; the diagonal denotes equal distance from normal. The inset enlarges members with $|D| \leq 2$ and $|R| \leq 2$ on the original expression scale; the full scatter retains all 10,478 measurable members. (B) Fixed cumulative weighted $\log_2$-expression differences for D, treatment change T (ABT-263 minus allograft), and R, with D + T = R. Blue bars show consensus values; thin solid/dashed arrows show quiescence/specialization weights. Each treatment-change trace begins at its own D endpoint. Open points show eight library projections per comparison, reusing 12 libraries, four per arm. T library projections are displayed at the consensus D endpoint plus the corresponding library projection; D and R library projections are displayed without a baseline shift. Projections compare a treatment library with the comparator mean or the treatment mean with a comparator library. Each weighting uses the sum of measurable absolute weights as its denominator. Allo, allograft; Nor, normal; ABT, ABT-263.

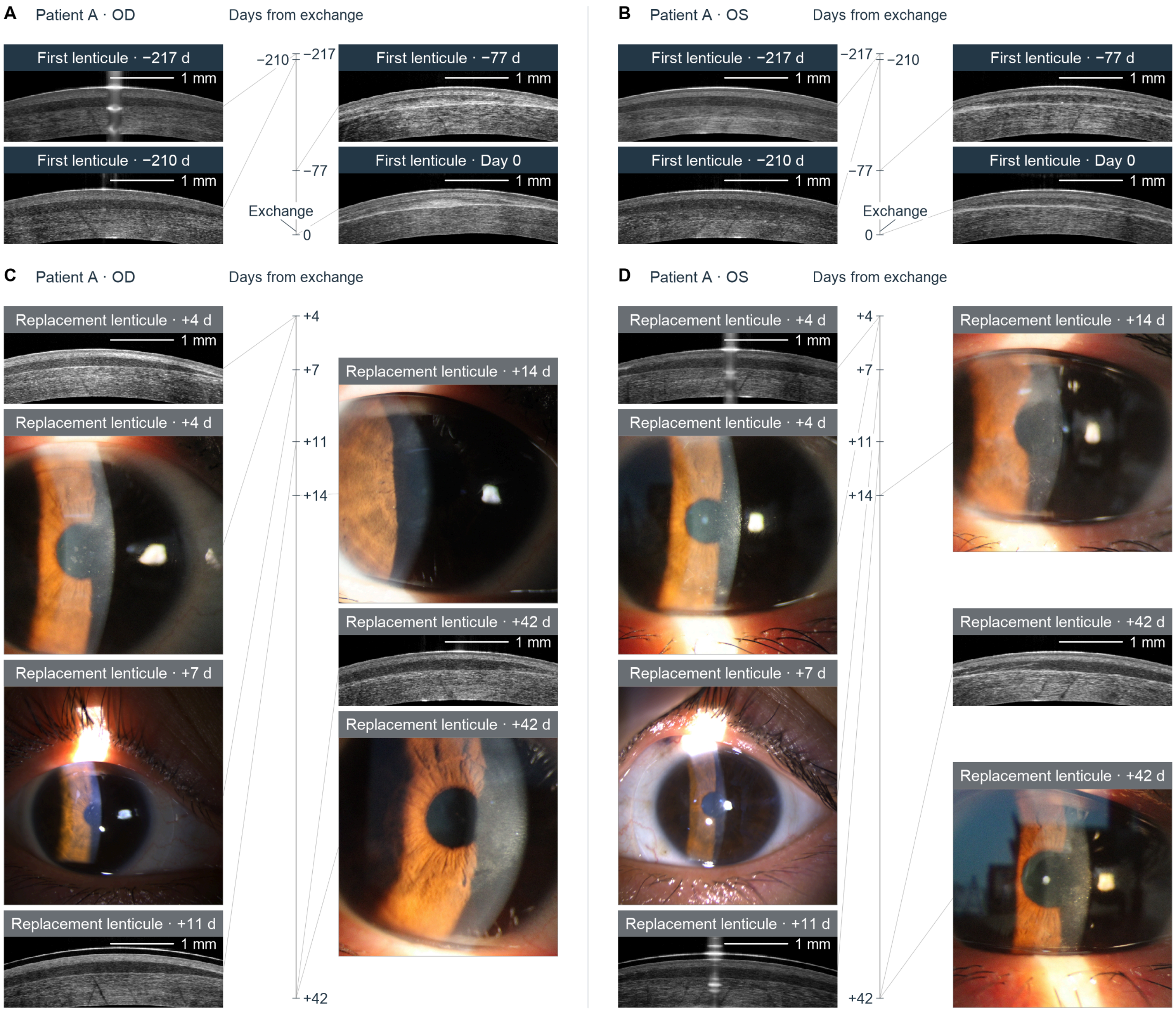


**Supplemental Fig. S18. Patient A's bilateral image series records first-lenticule opacity and changes after exchange.**

(A and B) First-lenticule OCT sections from OD and OS, respectively, at −217, −210, −77 and 0 days relative to exchange. (C and D) Replacement-lenticule series from OD and OS, respectively, with OCT at +4, +11 and +42 days and anterior-segment photographs at +4, +7, +14 and +42 days. The series contains seven bilateral OCT visits (14 eye-visits and 14 sections) and eight photographs. Dark/gray bands identify first/replacement lenticules; Exchange labels point to day 0 on the vertical axes in A and B. Images are arranged in two compact columns, with connectors linking each image to its actual day on a linear vertical axis. OCT scale bars, 1 mm. OD/OS, right/left eye; OCT, optical coherence tomography.

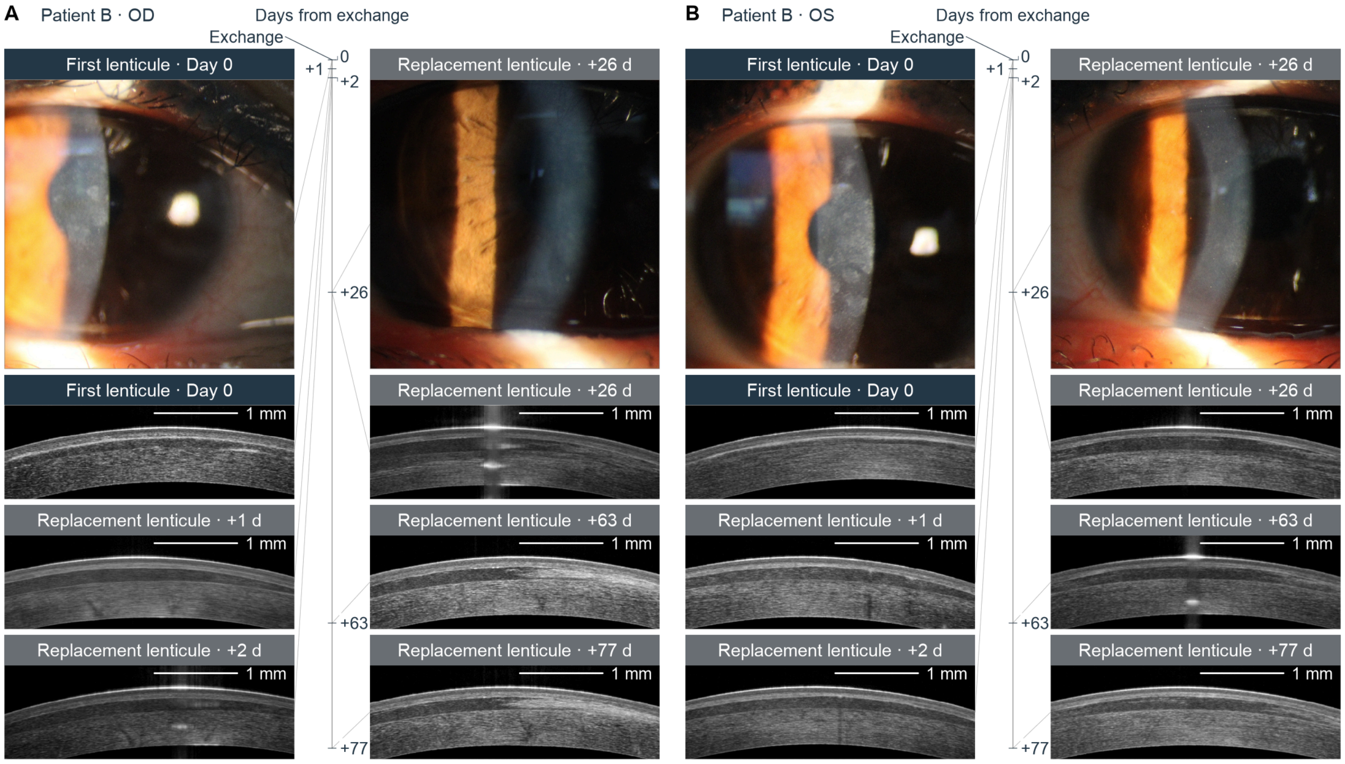
A
Patient B · OD
Days from exchange
Exchange
First lenticule · Day 0
Replacement lenticule · +26 d
First lenticule · Day 0
Replacement lenticule · +26 d
Replacement lenticule · +1 d
Replacement lenticule · +63 d
Replacement lenticule · +2 d
Replacement lenticule · +77 d
1 mm
0
+1
+2
+26
+63
+77
B
Patient B · OS
Days from exchange
Exchange
First lenticule · Day 0
Replacement lenticule · +26 d
First lenticule · Day 0
Replacement lenticule · +26 d
Replacement lenticule · +1 d
Replacement lenticule · +63 d
Replacement lenticule · +2 d
Replacement lenticule · +77 d
1 mm
0
+1
+2
+26
+63
+77

**C** Patient B · OD

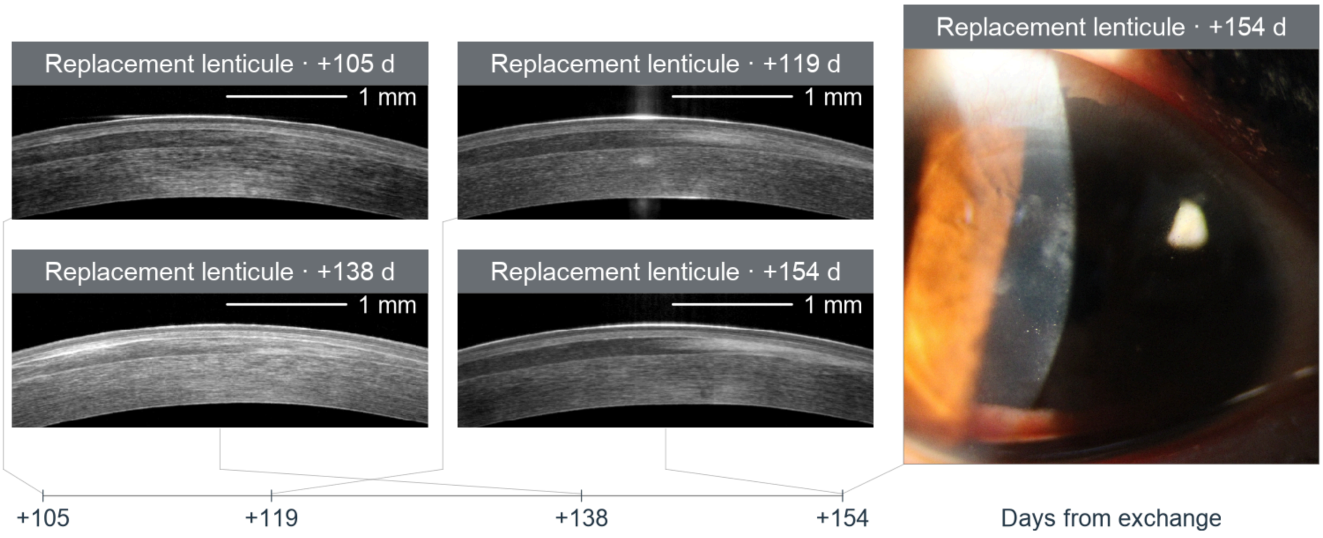


**D** Patient B · OS

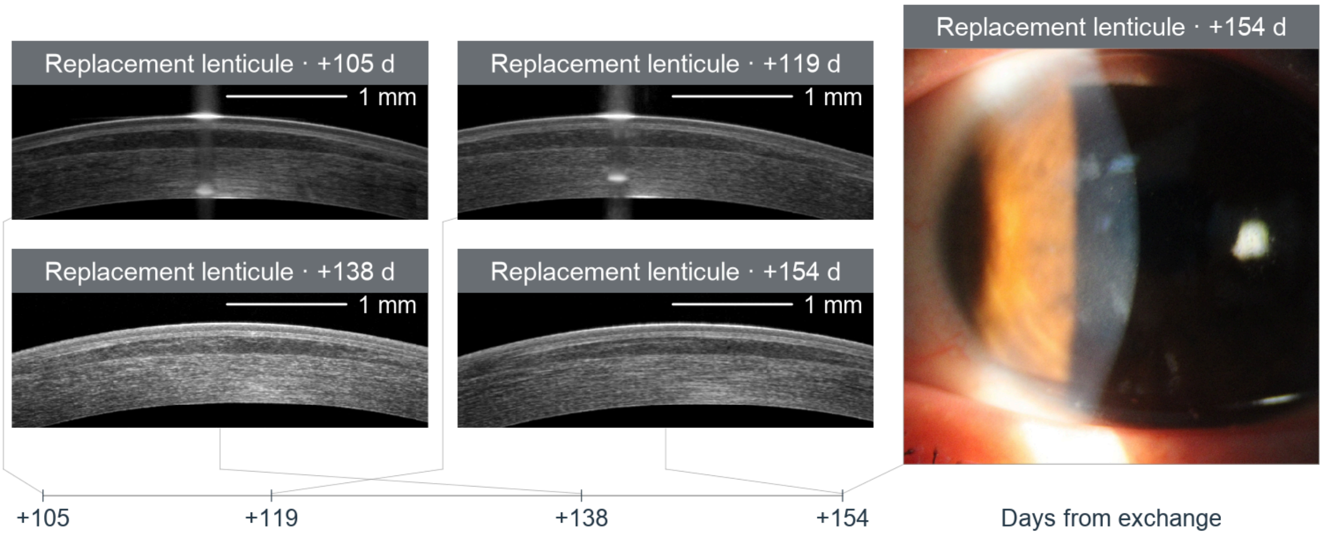


**Supplemental Fig. S19. Patient B's bilateral image series records early clearing and recurrent replacement-lenticule opacity.**

(A and B) OD and OS series from day 0 to +77 days. Day-0 images show the first lenticules before exchange; subsequent images show replacement lenticules. Exchange is labeled at day 0 on the vertical time axes in A and B. (C and D) OD and OS replacement-lenticule series from +105 to +154 days. Across both pages, ten bilateral OCT visits contribute 20 eye-visits and 20 sections, with six anterior-segment photographs. Dark/gray bands identify first/replacement lenticules. In A and B, images from days 0, +1 and +2 occupy the left column, and images from days +26, +63 and +77 occupy the right column; connectors link each image to its actual day on a linear vertical axis in each panel. In C and D, the four OCT sections form a two-by-two array, with +105 and +119 days in the upper row and +138 and +154 days in the lower row, alongside the same-eye photograph at +154 days. Images in C and D connect to their actual days on linear horizontal axes below the OCT arrays. OCT scale bars, 1 mm. OD/OS, right/left eye; OCT, optical coherence tomography.

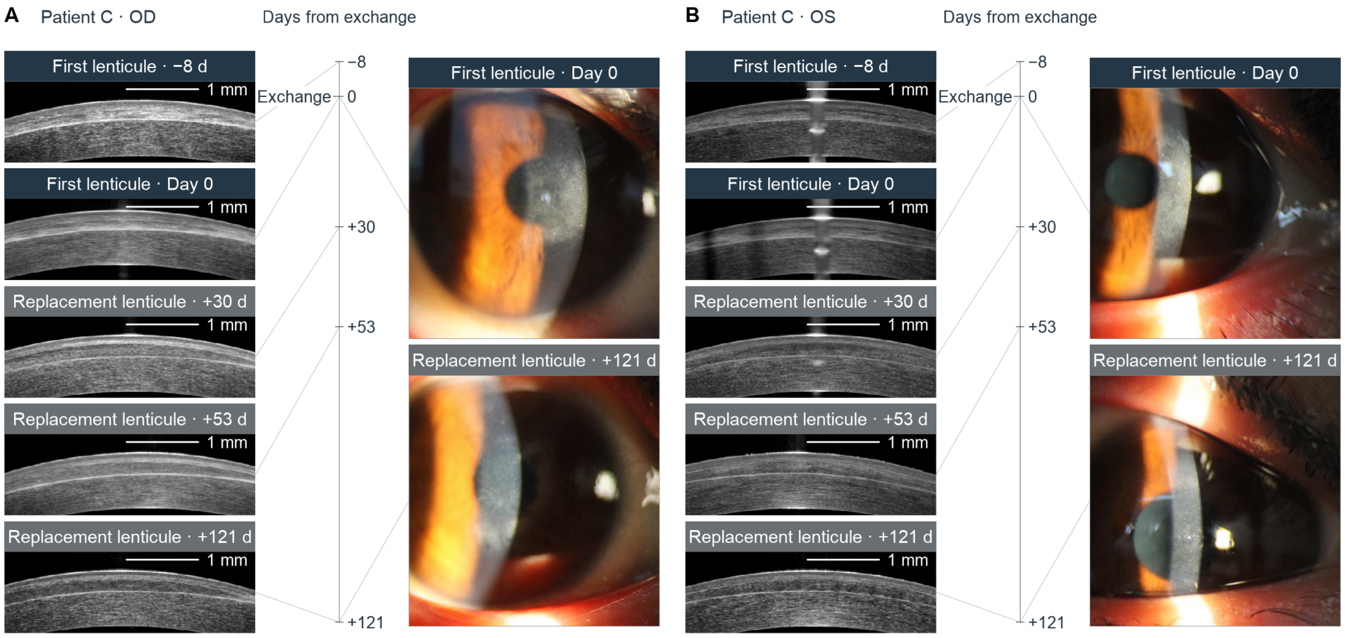


**Supplemental Fig. S20. Patient C's bilateral image series records opacity in both first and replacement lenticules.**

(A and B) OD and OS, respectively, at −8, 0, +30, +53 and +121 days relative to exchange. Dark bands identify first lenticules at −8 and 0 days; gray bands identify replacement lenticules at positive days. Exchange is labeled at day 0 on the vertical time axes in A and B. Five bilateral OCT visits contribute ten eye-visits and ten sections; four anterior-segment photographs show day 0 and +121 days in both eyes. OCT sections and photographs occupy separate columns linked to their actual days on a single linear vertical axis in each panel. OCT scale bars, 1 mm. OD/OS, right/left eye; OCT, optical coherence tomography.

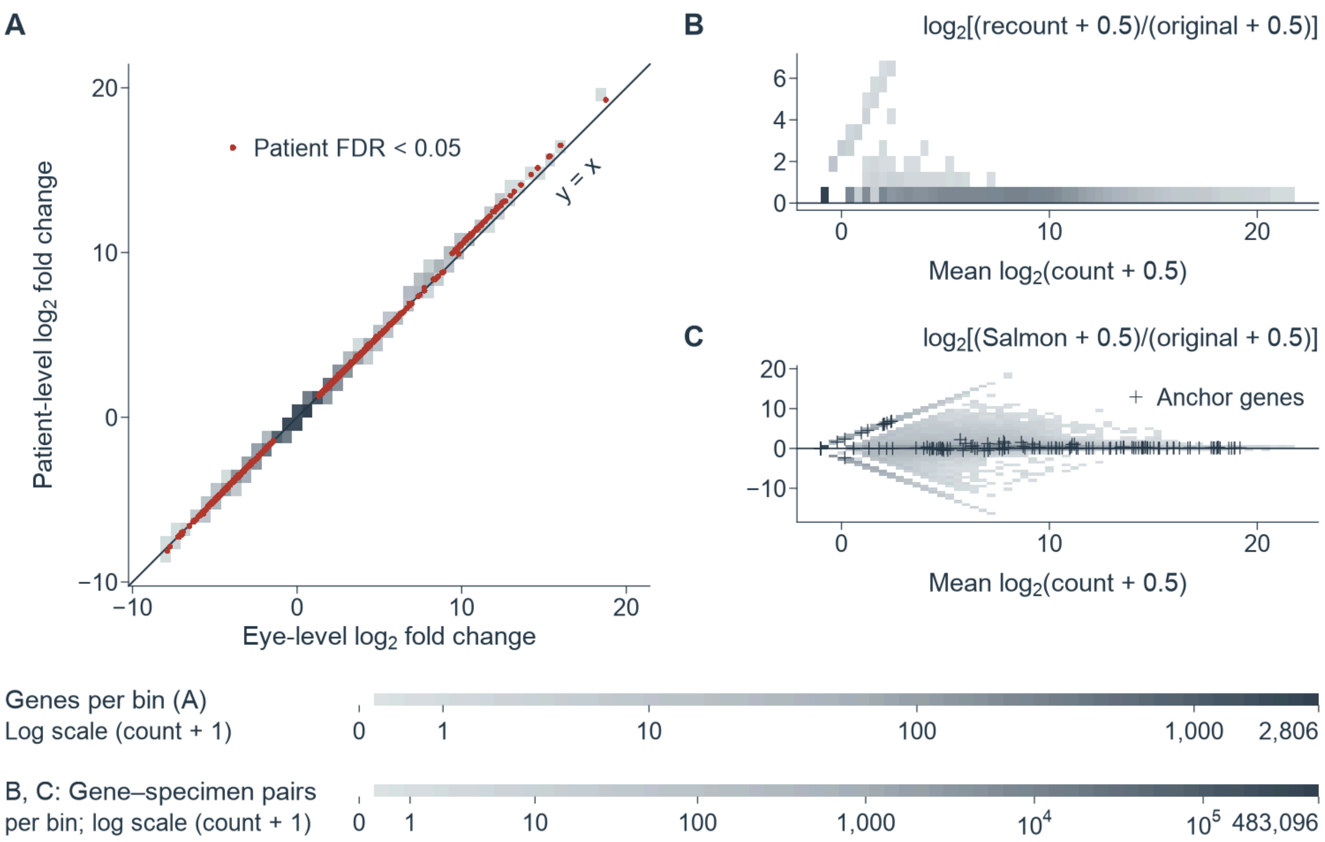


**Supplemental Fig. S21. Patient-level gene effects agree with eye-level effects, and recounting reproduces original counts.**

(A) Eye- versus patient-level log2 fold changes for 15,338 matched genes, with bilateral counts summed within patient. Red points identify 1,039 genes with patient-level false discovery rate (FDR) < 0.05; gray bins represent all matched genes. The diagonal marks equal effects. (B) Same-settings featureCounts recount and (C) Salmon estimates versus vendor original counts, with 704,820 gene–specimen pairs per panel. Horizontal positions are mean log2(count + 0.5); vertical positions are log2[(estimate + 0.5)/(original + 0.5)]. Crosses in C identify 15 anchor genes across 12 specimens. Gray shading encodes bin counts on a log(1 + count) scale, with a separate range for genes in A (0–2,806) and a shared range for gene–specimen pairs in B and C (0–483,096); horizontal lines mark zero in B and C.

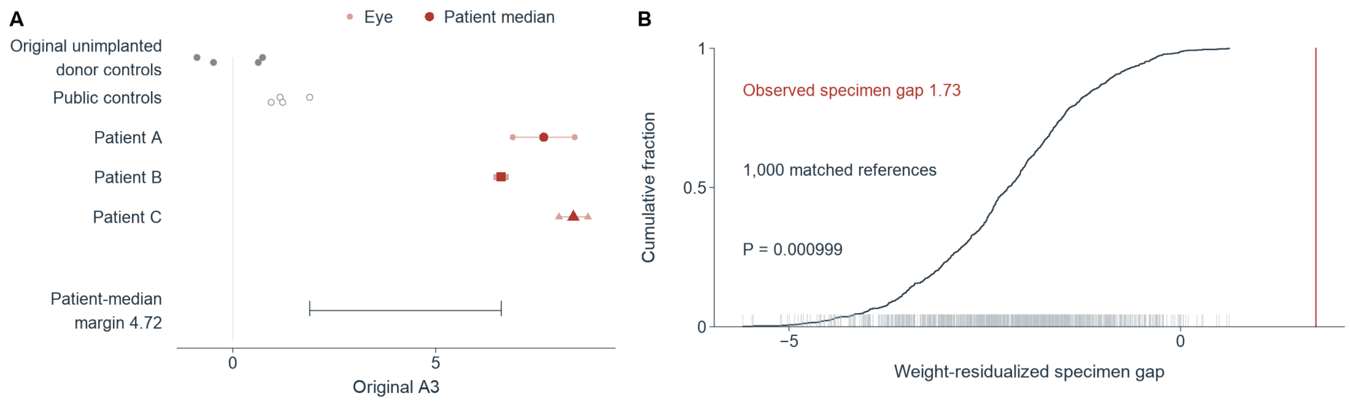


**Supplemental Fig. S22. Implant-remodeling scores separate failed lenticules from expanded controls and expression-matched references.**
(A) Original implant-remodeling scores (A3) for six failed eyes, three patient medians, four original unimplanted donor controls (filled gray) and four public controls (open gray). Pale points show eyes and dark red circles, squares and triangles identify medians for Patients A–C. The bracket marks the lowest patient median minus the highest control, 4.72 arbitrary units. (B) Empirical cumulative distribution and rugs for 1,000 expression-matched random-reference specimen gaps. The red line marks the observed lowest-eye-minus-highest-original-control gap on weight-residualized A3, 1.73. The empirical upper-tail P value is 0.000999.
